# Hierarchical clustering optimizes the tradeoff between compositionality and expressivity of task structures for flexible reinforcement learning

**DOI:** 10.1101/2021.07.20.453122

**Authors:** Rex G Liu, Michael J Frank

## Abstract

A hallmark of human intelligence, but challenging for reinforcement learning (RL) agents, is the ability to compositionally generalise, that is, to recompose familiar knowledge components in novel ways to solve new problems. For instance, when navigating in a city, one needs to know the location of the destination and how to operate a vehicle to get there, whether it be pedalling a bike or operating a car. In RL, these correspond to the reward function and transition function, respectively. To compositionally generalize, these two components need to be transferable independently of each other: multiple modes of transport can reach the same goal, and any given mode can be used to reach multiple destinations. Yet there are also instances where it can be helpful to learn and transfer entire structures, jointly representing goals and transitions, particularly whenever these recur in natural tasks (e.g., given a suggestion to get ice cream, one might prefer to bike, even in new towns). Prior theoretical work has explored how, in model-based RL, agents can learn and generalize task components (transition and reward functions). But a satisfactory account for how a single agent can simultaneously satisfy the two competing demands is still lacking. Here, we propose a hierarchical RL agent that learns and transfers individual task components as well as entire structures (particular compositions of components) by inferring both through a non-parametric Bayesian model of the task. It maintains a factorised representation of task components through a hierarchical Dirichlet process, but it also represents different possible covariances between these components through a standard Dirichlet process. We validate our approach on a variety of navigation tasks covering a wide range of statistical correlations between task components and show that it can also improve generalisation and transfer in more complex, hierarchical tasks with goal/subgoal structures. Finally, we end with a discussion of our work including how this clustering algorithm could conceivably be implemented by cortico-striatal gating circuits in the brain.

## 1 Introduction

Humans have a remarkable ability to quickly solve novel problems by leveraging knowledge from past experiences that bear some meaningful similarity to the problem at hand. One important way by which we do this is through structural generalization: most naturalistic tasks have an underlying latent structure which we can extract and transfer to new problems in novel ways [1, 2]. For instance, when travelling to work, we can abstract out knowledge of the underlying road network – recognising that it does not depend on the particularities of a single commute, such as the cloud patterns in the sky or the foliage pattern in the trees along the route – and transfer it to other commutes on the same network. Such capacity to abstract out and transfer latent task structures has also been observed in non-human primates, as first noted by Harlow in his account of “learning set” acquisition [3]. In contrast, artificial agents, in spite of their impressive ability to now outperform humans on a wide range of difficult tasks [4, 5, 6, 7], are notoriously brittle and struggle with similar knowledge transfer [8, 9].

Task structures play a central role in model-based reinforcement learning. Reinforcement learning has provided one of the most successful frameworks for studying and modelling intelligent decision-making, both in natural [10, 11, 12] and artificial intelligence [13, 4, 6]. In model-based reinforcement learning, an agent uses an internal predictive model of the task to simulate different possible courses of action in search of the best one, a process known as planning. In essence, the agent “thinks” before acting. Such models are needed for planning, but how an agent acquires such models to efficiently plan across contexts – and especially how they can be combined in novel ways to support generativity – remains an open question.

Compositional generalisation affords one mechanism. Indeed, the structures underlying naturalistic tasks are often compositional: they can often be decomposed into independent subcomponents that an intelligent agent can learn and recombine in novel ways. In the example of travelling to work, this task can be decomposed into two subcomponents: (i) knowing the location of the destination (a reward function) and (ii) operating a vehicle (such as a bike or car) to follow a route there (a mapping from primitive muscle actions to vehicle movements in physical space). The knowledge of how to drive is independent of the destination and can be transferred to tasks with new destinations; conversely, knowing how to get to a specific destination by car should imply knowing how to get there by bike.. The same principle applies to other domains of flexible behavior: for example, a musician can readily transfer instrument fingerings across different songs and songs across different instruments. Compositional generalisation allows agents to rapidly construct new models by composing familiar structural components together into novel combinations that may not have been seen before. Humans are known to rely on this mechanism to rapidly infer the correct model for novel tasks, yet the computations needed to support compositional generalisation have only started to be explored [14, 15]. On the other hand, artificial agents struggle with compositional generalisation; they instead treat entire tasks as monolithic structures, preventing them from leveraging knowledge about subcomponents that may recur in other tasks.

Compositional generalisation has been extensively studied in other domains, such as natural language processing [16, 17, 18] and concept learning [19, 20, 21, 22]. Yet in reinforcement learning, the study of compositional generalisation has been restricted primarily to compositional policy structures in the domain of hierarchical reinforcement learning [23, 24, 25, 26, 12, 27, 28, 29, 30, 31, 32]. Here, an overall policy is constructed by composing together sub-policies in the form of skills or solutions to subtasks. However, such approaches are restricted to situations in which an agent can sequentially apply subtasks, and typically apply in the alternative domain of “model-free” reinforcement learning. Because model-free agents require extensive trial-and-error interactions with the environment [13], they do not afford the potential to flexibly plan in new contexts that can leverage compositions of transition and reward functions. Indeed, most state-of-the-art deep reinforcement learning algorithms are model-free algorithms [4, 33, 34, 35, 36, 37, 38, 5] and thus suffer from the same limitations [9].

In this work, we address the problem of compositional generalisation over task structures using non-parametric Bayesian inference. When solving a new task, the agent will attempt to infer what latent task structure is generating the observed transitions and rewards, in the form of a Markov Decision Process (MDP). If the observed statistics are consistent with previously seen MDPs, the agent can transfer its knowledge of that structure to quickly solve the new task. Otherwise, a new structure can be created and added to the pool of task structures that can be leveraged in the future. At its core, this framework is a clustering algorithm, as task contexts that share a common structure are clustered together. Previous work has shown that this framework can account for how humans are able to reuse previously learned structures while also acquiring new ones with minimal interference [39, 40, 1]. Such models have been further extended to sequential decision making scenarios [41, 14] and to situations in which agents can reuse deeper latent state abstractions in which new tasks can be learned by analogy with previous tasks [42]. Note that this framework is normative; approximate resource-constrained implementations are also possible. For example, models of hierarchically nested cortico-striatal gating circuits have offered a mechanistic account for how the brain might approximate such structure learning and transfer [43, 40], and human studies have found signatures of latent task structures in the brain [44, 45, 1, 46, 2, 47].

Until recently, however, these frameworks have only considered how agents could learn and reuse task structures as a whole and therefore could not support compositionality. Although an agent that memorises and reuses entire structures (combinations of transition and reward functions) can eventually solve any task, this strategy suffers from two main limitations. The first issue is one of scalability. If a structure is compositional, then the number of distinct structures possible will scale exponentially with the number of components that make up the structure; that is, if a structure consists of *n* components, each with *d* possibilities, then the number of possible structures scales as 𝒪(*d*^*n*^), which can quickly tax a finite memory capacity. The second issue is one of transfer and sample efficiency. An agent that memorises entire structures will be unable to transfer repeating structural components. Even if two MDPs have a substantial number of overlapping components, the agent will relearn each structure from scratch, making this approach highly inefficient. Moreover, this combinatorial explosion implies that many structures are never seen twice in exactly the same way in an agent’s lifetime, further reducing the benefits of memorising entire structures.

Recent work has extended the Bayesian framework to allow for compositionality and to compare to patterns of human generalization [14, 15]. In this extension, the agent maintains a factorized representation of structural components (for example, the *action-outcome mapping function*, akin to the mapping of primitive leg actions to bike movements, and the *reward function*, akin to the desired destination). When encountering a novel context, the agent attempts to infer each of the structural components independently of the others, allowing it to recompose new task structures, akin to biking for the first time to a destination one normally drives to. This is also a clustering algorithm, but now a task is assigned to multiple clusters, one for each component that makes up the structure. Because the components in these structures are treated statistically independently of each other, we refer to these structures as *independent structures*.

Nevertheless, it is not always appropriate to treat each task component as completely independent of the others. For one thing, it can still help to learn and transfer entire *joint structures* particularly when these recur in natural tasks. For instance, one might always bike to ice cream parlours but drive to work, making these specific joint structures more privileged and therefore worth memorising for future reuse. The set of tasks natural agents encounter throughout the course of their lifetimes, however, are never strictly joint or independent but are better characterised as recurring joint structures existing amongst otherwise independent statistics. Moreover, the correlational structure between structural components can be hierarchical. For instance, when travelling, the choice of transport method can depend not only on the destination within a city but also on higher-order contexts like the city itself. In Venice or Amsterdam, one could consider boating to their favourite canal-side restaurant, but not if they are in the suburbs of Toronto; alternatively, one could consider driving to many destinations in most cities, but not in Venice; and one can often rickshaw to many destinations in Asian cities but not elsewhere. The city thus constrains the possible correlations that can exist between the mode of transport and the destination.

Here, we develop a novel computational strategy to accommodate a wider range of correlational structures between model components. Instead of independent or joint agents, we introduce a single, more powerful, clustering agent based on the hierarchical Dirichlet process [48] that can learn the appropriate degree of covariance in a context-dependent fashion. Like the independent agent, our agent maintains a factorized representation over task components, allowing for components to be learnt and generalized independently of each other. Simultaneously, it also estimates the covariances between these components, thereby affording generalization over a wider range of compositional structures. We validate our approach on a variety of tasks with independent, joint, and mixed statistics and show that only the hierarchical agent is able to perform well across all tasks. Finally, we show that the model’s hierarchical structure can be immediately applied to improve generalization and transfer in more complex tasks with hierarchical goal/subgoal structures.

### 1.1 Paper layout

This paper is organised as follows. In the first half, we focus on non-hierarchical tasks to illustrate the utility of our framework in simple settings. In section 2, we explain the structure of the tasks we shall consider as well as the architecture of the various agents we shall use. Section 2.1 introduces the compositional gridworld tasks and contextual Markov decision process framework that we work with, while section 2.2 explains the agent architectures. Section 2.3 explains four versions of our compositional gridworlds intended to elucidate the relative adaptability of the different agents we consider and presents the results of our simulations. In section 3, we scale our approach up to hierarchical tasks of greater complexity, again presenting the task structure, the agent architectures, and the results of our simulations. We compare our work to related works in section 4, considering the limitations and other issues that future work will need to address. And finally, we conclude in section 5.

## 2 Task structure and agent architectures

### 2.1 Gridworld task structure

We consider a continual learning setting where each task has an explicitly compositional structure, based on tasks described in [14]. The agent must navigate towards a hidden goal in a series of 6 × 6 gridworld rooms (Fig. 1), where it has to learn both (i) the action-outcome mapping function (which actions will take it from one location to another, analogous to the relationship between primitive muscle actions and the resulting locomotion on a bike) and (ii) the reward function (the rewarding locations or goals). A flexible agent will be able to recognize when it can reuse a previously observed mapping function in the service of reaching a different goal (i.e., if these are independent across rooms) but could also infer when a given mapping function is informative about the likely goal location.

**Figure 1:**
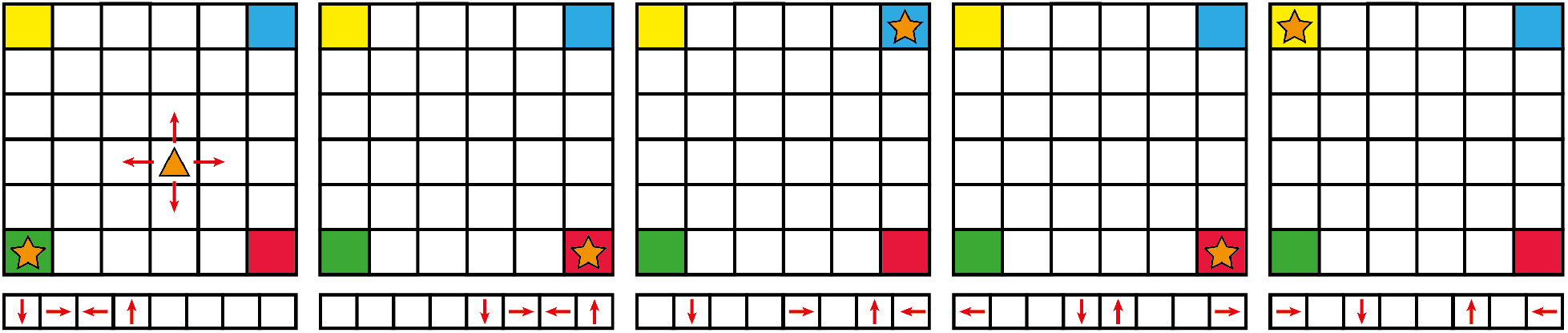
Gridworld task: the agent (triangle) navigates a series of 6 × 6 gridworld rooms where it can move in any of the four cardinal directions (red arrows). Each room has a hidden goal state (star) located in one of the corners. The agent moves by pushing one of eight possible keys, but the mappings from keys to cardinal movements (bottom) vary from room to room. A mapping is generated by randomly assigning arrows to keys, and pressing a blank key (no arrow) results in no movement. This mapping assignment is hidden from the agent, and thus must be inferred from its interactions with the gridworld.

The agent starts from a random location in each room and must navigate to a hidden goal by moving along the four cardinal directions. To move, the agent selects one of eight possible primitive actions (Fig. 1, bottom), hereafter referred to as “keys” based on corresponding human experiments in which subjects had to move by pressing keys on a keyboard [15]. Only four of the keys will move the agent while the others will do nothing. Moreover, the mapping from keys to movements (i.e., which key takes the agent up, down, left and right) varies from room to room and is unknown to the agent – it must be learnt from its interactions with the gridworld. The goal is also hidden from the agent so it must also learn the reward function from its interactions with the rooms. The goal appears in one of the four corners (Fig. 1, star), but the agent is not endowed with that information either. In sum, the composition in each of our gridworlds is of a mapping function paired with a reward function, and the agent has to infer both in order to plan in novel contexts.

Formally, each task can be modelled as a contextual Markov decision process (cMDP) [49] consisting of the sextuple (𝒞, 𝒮, 𝒜, 𝒯(*c*), ℛ(*c*), *γ*); here, 𝒞 is a set of contexts, *c* ∈ 𝒞, 𝒮 and 𝒜 are spaces of states and *primitive* actions, *γ* ∈ [0, 1) is a discount factor, and 𝒯 (*c*) and ℛ(*c*) are context-dependent transition and reward functions. Here, contexts are *observed* and denote room identity; in our implementation, contexts are discrete, although in more naturalistic settings, they could be a distinguishing visual feature of the room like wall or floor colours or some sign posted somewhere in the room. The state *s* ∈ 𝒮 corresponds to the agent’s position in the room, *s* = (*x, y*). The space 𝒜 is the set of eight possible keys. In our tasks, the agent is rewarded if it enters a goal state; as this is a function of position and context only, the reward function ℛ(*c*) takes the form

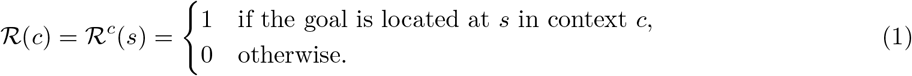

The transition function 𝒯 (*c*) expresses the probability of reaching state *s*′ when taking action *a* in state *s*; that is, 𝒯 (*c*) = 𝒯^*c*^(*s*′, *a, s*) = *p*^*c*^(*s*′ | *s, a*). In our tasks, this can be factorized into two components. The first component expresses state transitions in terms of the cardinal movements, namely *𝒜*_*card*_ = {*up, down, left, right, stay*} and abstracts out the transition’s dependence on the room’s spatial layout. Because all rooms share the same spatial layout, this component does not depend on context and hence takes the form *p*(*s*′|*s, a*_*card*_), where *a*_*card*_ ∈ *𝒜*_*card*_. The second component describes how primitive actions map onto cardinal movements and takes the form

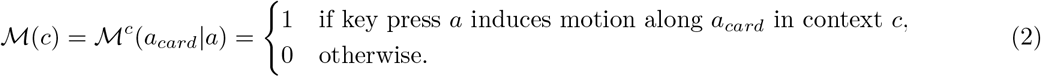

This component introduces context-dependence, reflecting the fact that mappings can depend on environmental conditions or other contextual factors; for instance, one needs to execute different motor actions depending on whether one is driving a car or pedalling a bike, and an aerial drone navigating to a goal needs to adjust its motor outputs depending on wind conditions. This is the mapping function that the agent must infer. Composed together, the full state transition function is

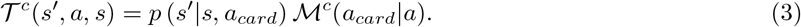

In this paper, we shall refer to the Markov decision process (MDP) for an individual context (that is, a specific room or gridworld) as a *task* and the entire cMDP (that is, the entire set of rooms or gridworlds that an agent must solve) as the *environment*.

#### Planning in compositional cMDPs

In these tasks, the agent must learn ℛ^*c*^ and ℳ^*c*^ from its interactions with the rooms (all other MDP components are given). If the goal is known, the agent can plan the optimal sequence of cardinal movements to reach it. The optimal policy is obtained by choosing movements that maximize the action-value function 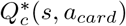,

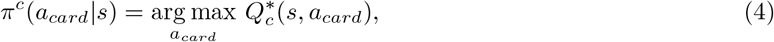

where 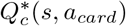 is the solution to the Bellman optimality equation [13]

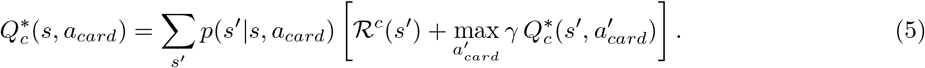

Both 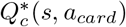 and *π*^*c*^(*a*_*card*_|*s*) can be obtained by model-based policy iteration [13]. Note that the only room-dependent component here is the reward function ℛ^*c*^; hence only the goal location is needed for the agent to plan a policy in terms of cardinal actions.

If ℳ^*c*^ is known, the agent can then implement this policy by selecting the appropriate keys. Combining (2) with (4), one can show that this solves the Bellman optimality equation in terms of the key presses *a* ∈ *𝒜*,

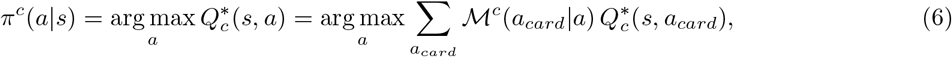

where, by an abuse of notation, we use *π*^*c*^(*·*) and 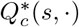 to denote the policy and action-value functions in terms of cardinal movements or key presses according to whether the argument is an action from *𝒜*_*card*_ or *𝒜*. The action-value functions for *a* are easily obtained from those for *a*_*card*_,

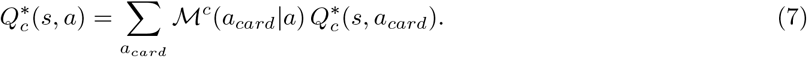

Thus, the reinforcement learning problem critically reduces to one of inferring the correct ℛ^*c*^ and ℳ^*c*^.

As the agent moves from room to room, functions ℳ^*c*^ and ℛ^*c*^ may repeat independently of each other, and agents that leverage knowledge about familiar ℳ^*c*^ and ℛ^*c*^’s will exhibit an advantage to quickly solve novel rooms. However, depending on the task statistics, ℳ^*c*^’s and ℛ^*c*^’s can also be correlated, so there will also be a distinct advantage towards agents that can learn and exploit this correlational structure.

We have designed four different versions of our compositional environment testbed. Each is designed to highlight the relative expressivity of the different agents we consider, each of which uses a different strategy for clustering and transferring task structures. Before discussing our environments, we first introduce these agents.

### 2.2 Clustering agents

Past work in both Pavlovian [39] and instrumental conditioning [40, 43] have cast context-dependent generalisation as a clustering problem. In this framework, different contexts are clustered together into a common, latent generative task state if they share similar task statistics. These latent states are not to be confused with MDP states but can be regarded as higher order states that condition entire MDPs or stimulus-response mappings – indeed, models of the hierarchical cortical-striatal circuitry suggest that higher, more anterior, levels of the circuitry select for different MDP structures, conditioning how lower level circuitry selects actions [43, 44, 40, 1]. Clustering allows an agent to maintain distinct task representations in different clusters without them interfering with one another, but also allows for rapid transfer of knowledge between tasks when they are inferred to be in the same cluster. This framework has been extended to allow for compositional generalisation [14, 15] and for reuse of compressed representations of abstract task states even when reward and transition functions change [42].

Formally, cluster assignment is framed as a Bayesian inference problem where the agent must infer which latent cluster is generating the current task statistics. To allow for the possibility of an infinite number of clusters, a Dirichlet process (DP) prior is used for cluster assignment. This does not require specifying the total number of clusters beforehand, making it a non-parametric Bayesian model. A DP is defined formally in terms of a base distribution *H* and a concentration hyperparameter *α* that controls the probability of generating new clusters [50],

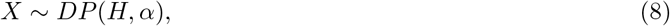

where *X* is a clustering drawn from the DP.

To impose parsimony in the prior cluster assignment, we assume that those clusters that are most popular across contexts are more likely to apply in a new context [39, 40], consistent with human subject priors during generalization [1], and reflected in the Chinese restaurant process (CRP) [51, 52]. The total number of clusters *K* at any moment in time is finite, but new clusters can be created and added to the set as needed, and can theoretically approach infinity in the limit of infinite contexts. However, the parsimony prior gives the agent a higher chance of reusing an existing cluster, in proportion to its past popularity, with more popular clusters receiving a greater probability of assignment. If contexts *c*_1:*n*_ have already been assigned to clusters {1, …, *K*}, where *K* ≤ *n*, then the prior probability that new context *c*_*n*+1_ belongs to cluster *k* is given by

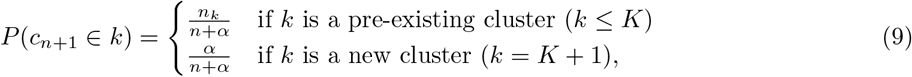

where *n*_*k*_ ≥ 1 is the total number of contexts already assigned to cluster *k* and *α*, here, controls the probability of assignment to a novel cluster. As data 𝒟 is accumulated, cluster assignment is updated via Bayesian inference,

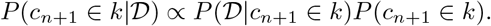

The data distribution for each cluster *P* (𝒟|*c*_*n*+1_ ∈ *k*) is determined by a likelihood function, the parameters for which must also be inferred; the base distribution *H* specifies its prior distribution. The number of clustering hypotheses grows exponentially with the number of contexts, making representation of all hypotheses impractical. Instead, we only retain a subset which gets filtered each time a new context is added. Exact details of our particle filtering can be found in the Supplement.

We shall now describe the various agents that we shall be comparing, each of which uses a different strategy to cluster contexts. Not that the clustering strategy is the only difference between these agents; it is the structure of the clustering hypothesis space that determines the Bayesian priors used to support transfer to novel contexts, and it is this alone that is responsible for any performance differences between the agents in our simulations below. At the level of individual reward and mapping clusters, all agents operate identically.

#### 2.2.1 Independent clustering agent

Compositional generalisation requires the ability to generalize task components (e.g., mapping and reward functions) independently of one another, and a factorized representation of task components is necessary to support this. Previous work [14, 15] has accomplished this by maintaining two independent clusterings of contexts (that is, two separate but parallel Dirichlet processes); one clusters according to whether contexts share the same mapping function ℳ^*c*^ and another according to whether they share the same reward function ℛ^*c*^ (Fig. 2a). Each mapping cluster *k*_ℳ_ defines a mapping likelihood function given by (2), with all contexts in the same cluster sharing a common ℳ^*c*^. However, the parameters of this common likelihood is unknown and must be inferred; the posterior is given by

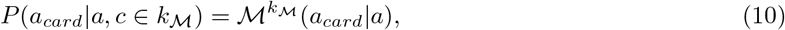

where 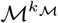 takes the form of a Bernoulli distribution. Likewise, each reward cluster *k*_ℛ_ defines a reward likelihood function ℛ^*c*^(*s*) given by (1). The likelihood parameters are also unknown and inferred using the posterior

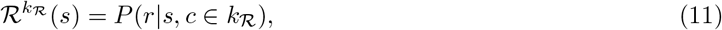

where 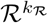 is also a Bernoulli distribution and specifies the probability of observing reward *r* in state *s*. Following [14], we shall refer to an agent which clusters contexts in this way as an *independent clustering agent*. Note that the entire inference process can be viewed in analogy to Bayesian model comparison, with inference of the correct clustering analogous to model inference based on available evidence, and inference of ℳ^*c*^ and ℛ^*c*^ analogous to inference of parameters for a given model.

**Figure 2:**
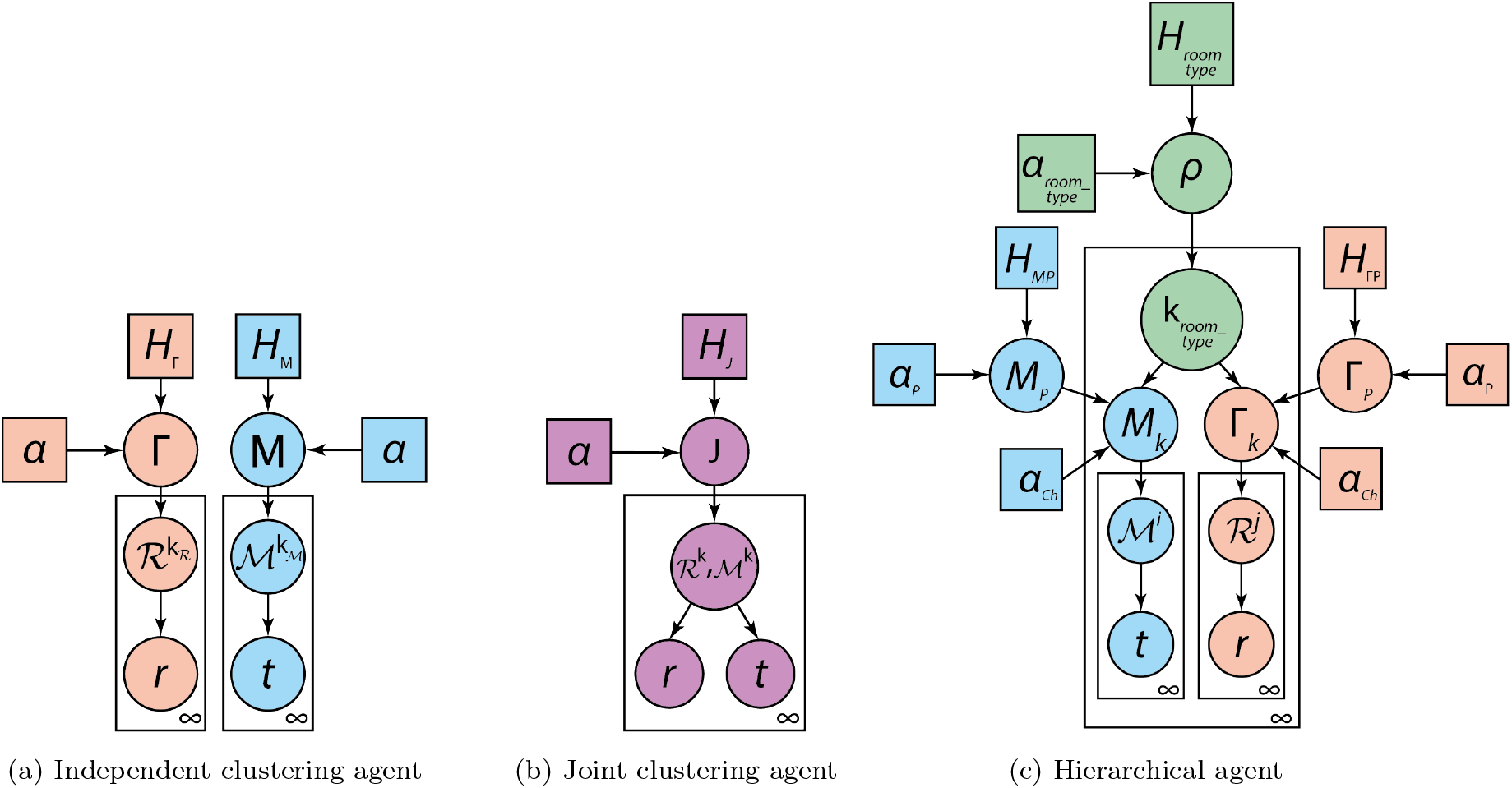
The independent clustering agent (a) maintains separate clusterings of rewards (red) and mappings (blue). A base distribution *H*_Γ_ generates a clustering Γ of reward functions 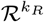, which in turn generates the rewards *r* observed in the room. A separate base distribution *H*_*M*_ generates clusterings *M* of mappings 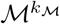, which in turn generates the observed transitions *t*. In the joint clustering agent (b), a single base distribution *H*_*J*_ generates clusterings *J* of reward functions ℛ^*k*^ paired with mappings ℳ^*k*^. These, in turn, generate *r* and *t*. The hierarchical agent (c) consists of three main components: (i) a DP of room types (green), (ii) an hDP for mappings (blue), and a separate hDP for reward functions (red). A base distribution *H*_*room_type*_ generates clusterings *ρ* of different room types *k*_*room_type*_, each defining different statistical associations between mappings and rewards. Outside room clusters, base distributions *H*_*MP*_ and *H*_Γ*P*_ generate environment-wide clusterings, *M*_*P*_ and Γ_*P*_, of mapping ℳ and reward ℛ functions, respectively. As with the independent agent, this separate clustering factorises mappings and rewards across the environment so that they can be recombined in new ways inside new room types. *M*_*P*_ and Γ_*P*_ in turn generate children clusterings *M*_*k*_ and Γ_*k*_ within room clusters (outer plate), thereby defining room-specific distributions over ℳ and ℛ. Each cluster in *M*_*k*_ and Γ_*k*_ defines a mapping ℳ^*i*^ and reward ℛ^*j*^ function, which in turn generates the observed transitions *t* and rewards *r*.

#### 2.2.2 Joint clustering agent

An alternative to independent clustering is to cluster ℳ^*c*^ and ℛ^*c*^ jointly in a single DP (Fig. 2b), much like what many previous non-compositional agents do. In this case, the likelihood function for a transition (*s, a, a*_*card*_, *s*′, *r*) is

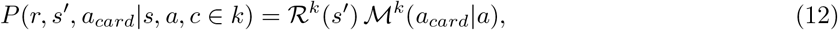

where, again, ℳ^*k*^(*a*_*card*_|*a*) and ℛ^*k*^(*s*′) are inferred using Bernoulli posteriors. As in [14], we shall refer to this as a *joint clustering agent*.

#### 2.2.3 Meta-agent

As noted above, an agent that clusters components independently can compositionally generalize to new tasks but will fail to leverage statistical structure when the reward and transition functions are informative of each other. For instance, in navigating a city, bike routes might correlate more with destinations such as ice cream parlours or parks but car routes might correlate more with work destinations, and an agent could theoretically recognise and exploit such correlations. In gridworld tasks with sparse rewards, an agent will generally learn ℳ^*c*^ first and only learn ℛ^*c*^ when it reaches a goal. There will therefore be a distinct advantage towards agents that can predict ℛ^*c*^ from ℳ^*c*^, when such a predictive relationship exists. In the extreme case where ℳ^*c*^ is entirely predictive of ℛ^*c*^ (for instance, when each ℳ is paired with exactly one goal, making the goal entirely predictable from the mapping), the joint clustering agent will outperform the independent clustering agent, as shown in prior work [14] and as we shall see below. Humans are capable of flexibly adapting to both types of task statistics [15], which raises the question of how this flexibility can be captured in artificial agents. To address this, [14] developed a mixture model that infers whether environment-wide task statistics are better modelled by independent or joint clustering models and then acts accordingly,

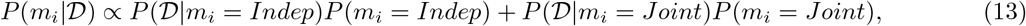

where *m*_*i*_ ∈ {*Indep, Joint*} denotes the model type. This was called a *meta-agent* because it performed inferences over two lower level (joint and independent) inference architectures. The main limitation with this approach is that all tasks get projected onto either joint or independent statistics, yet most naturalistic environments have statistics that fall somewhere in between. In particular, tasks may actually include both joint task structures existing alongside several contexts with independent structural statistics, and such an agent would fail to discover and reuse these different types of structural statistics. For instance, in our earlier navigation example, one might always drive to work destinations, making this task special and its joint structure something worth learning and transferring as a whole, but one might otherwise favour biking or driving equally for any other destination. The meta-agent would be suboptimal in such cases, and a more powerful algorithm is therefore needed to learn mixed statistics like these.

#### 2.2.4 Hierarchical agent

Here, we develop a novel, more expressive *hierarchical agent*, based on the hierarchical Dirichlet process. In contrast to the meta-agent, which can only be optimal if the environment is uniformly joint or independent, our agent is capable of learning and exploiting a wider range of statistical task structures that may exist alongside each other within the same environment, a situation more typical of naturalistic environments. Importantly, our agent performs inference over a more expanded hypothesis space covering a broader range of statistical structures between ℳ and ℛ, but which includes independent and joint clustering as special cases.

The key idea behind the hierarchical agent is to have it learn and transfer task *components* as well as any statistical relationship between those components (generative *structures* that may produce them jointly). It does this by inverting a non-parametric, hierarchical generative model for these components (Fig. 2c). In our navigation task, the different rooms can be regarded as belonging to different “room types” that generate different statistics in ℳ and ℛ. More precisely, different room types will generate different conditional distributions over ℳ and ℛ. Some types can generate a deterministic distribution over ℳ and ℛ to yield joint structures, while other types can generate more independent statistics, thus allowing both joint and independent generative structures to coexist alongside each other in the same environment. The idea behind the hierarchical agent is to invert this generative model and cluster contexts not just according to mapping and reward functions (which the independent agent does) but also according to the room types that generated them. In this way, the hierarchical agent clusters contexts not just according to task components but also according to generative structures.

The hierarchical agent introduces, at the highest level, a DP (Fig. 2c, green nodes) for clustering contexts by room types (note the same framework could be used for any type of cMDP, but here we focus on rooms in gridworlds). Each room-type cluster defines its own pair of DPs for clustering mapping and reward functions (Fig. 2c, blue and red nodes inside plates). These room-specific DPs model distributions over ℳ and ℛ that are conditional on the room type. Because different room types can induce different distributions over ℳ and ℛ, statistical associations between ℳ and ℛ may be prevalent in some room types (that is, ℳ may be predictive of ℛ). The agent can also leverage information about the prevalence of each mapping and reward function across the entire environment (all rooms); indeed, it additionally maintains a pair of DPs outside the rooms that cluster mapping and reward functions across the environment (Fig. 2c, blue and red nodes outside plates), and the room-specific clusters are assumed to be drawn from these environment-wide clusters, hence forming a hierarchical Dirichlet process (hDP). In the terminology of hDPs, the room-independent DPs serve as *parental* processes while the room-specific DPs serve as *children* processes. The room-independent DPs serve the essential role of factorising task components across the entire environment so that they can be recombined in novel ways in novel room types. Moreover, the same room-independent parental cluster can be inherited by multiple room-specific children DPs, mirroring the way in which the same task components can be used in multiple different structures, which is the essence of compositional generalisation. Indeed, it is important to note that all children clusters that inherit from the same parental cluster will model the same ℳ or ℛ function, in that any Bayesian update to a function made in one room will immediately update the function for all other rooms that use it.

Note that the hierarchical agent is a fundamentally different solution than the meta-agent, which only tries to infer whether an entire task domain is independent or joint, but cannot discover any substructure within the domain. As an analogy, in the navigation example, the hierarchical agent can discover that travel by bike tends to co-occur with ice parlour destinations and therefore treat this as one generative structure, while simultaneously recognising that most other destinations and modes of transport tend to be uncorrelated and therefore independent of each other; the meta-agent is unable to recognise this, instead treating the relationship between modes of transport and destinations as always independent or always joint according to whichever strategy gives the best performance.

In sum, the hierarchical agent (Fig. 2c) consists of several components, each serving a distinct purpose:

i. a DP, *P* (*c* ∈ *k*_*room*_*_*_*type*_), that clusters contexts according to room types (allowing for different latent generative structures within the same environment). *P* (*c* ∈ *k*_*room*_*_*_*type*_) is governed according to (9) with hyperparameter *α* = *α*_*room*_*_*_*type*_ and corresponds to the green nodes in Fig. 2c. Note that although we use “room types” here to capture an environment made up of multiple rooms, the formulation is general and could equally apply for example to a “musical instrument type” in an environment of instruments, “sport types” in a decathalon, etc. Formally, the DP is given by

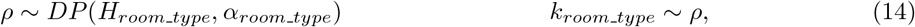

where base distribution *H*_*room*_*_*_*type*_ generates a clustering *ρ* of contexts according to room type and is controlled by a concentration hyperparameter *α*_*room*_*_*_*type*_. This clustering *ρ* in turn generates specific room-type clusters *k*_*room*_*_*_*type*_.
ii. a pair of parental DPs (blue and red nodes, respectively, outside plates in Fig. 2c) for clustering task components ℳ and ℛ across the entire environment. As described above, these parental DPs play the essential role of factorising task components ℳ and ℛ across the environment so that ℳ and ℛ can be learned independently of each other and reused in novel combinations inside novel structures (i.e. the “room types” in our framework). The room-specific children DPs (which we describe below) will inherit their clusters from these DPs.^1^ Formally, these DPs are given by

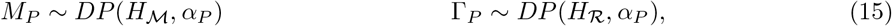

where base distributions *H*_ℳ_ and *H*_ℛ_ generate clusterings *M*_*P*_ and Γ_*P*_ of contexts according to whether the contexts share ℳ and ℛ; clustering is controlled by the concentration hyperparameter *α*_*P*_. (Further information on the specification of *H*_ℳ_ and *H*_ℛ_ is given below.)
iii. within each room cluster, a pair of DPs, *P* (*c* ∈ *k*_ℳ_|*c* ∈ *k*_*room*_*_*_*type*_) and *P* (*c k*_ℛ_ *c* ∈ *k*_*room*_*_*_*type*_), for clustering contexts according to common ℳ and ℛ functions respectively (blue and red nodes, respectively, inside plates in Fig. 2c). These clusterings inherit their clusters from the parental processes above and allow the agent to represent potential covariances between task components according to the corresponding generative structure. They are also governed by (9) with hyperparameter *α* = *α*_*Ch*_. Each cluster in this clustering determines corresponding likelihood functions ℳ and ℛ, for which the parameters are unknown and inferred through Bayesian posteriors (10) and (11) respectively – these define the base distributions *H*_ℳ_ and *H*_ℛ_ for the parental processes introduced above. Formally, the DPs are given by

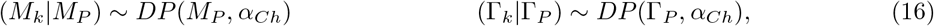

where parental clusterings *M*_*P*_ and Γ_*P*_ from (15) now serve as the base distributions for these DPs. Clusters in *M*_*P*_ and Γ_*P*_ in turn define likelihood functions for mappings and rewards

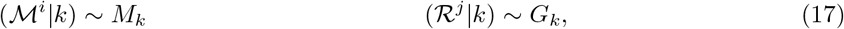

where *i* and *j* index clusters in *M*_*k*_ and *G*_*k*_. These functions, in turn, generate the observed transitions *t* = (*a*_*card*_|*a*) and rewards *r*

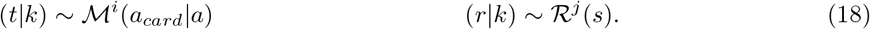 As mentioned above, the parameters for ℳ and ℛ are unknown to the agent and must be inferred. Note that if mapping cluster *i*_1_ in room *k*_1_ and mapping cluster *i*_2_ in room *k*_2_ inherit from the same parental cluster *k*_ℳ_ in *M*_*P*_, then their corresponding posteriors for the parameters of ℳ will be the same (i.e. 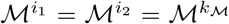) even though the clusters belong to different room types; likewise for ℛ. In this way, information about shared task components gets pooled across different room structures.

For ease of reference, all variables appearing in the hierarchical agent has been summarised in Table 1.

**Table 1:**
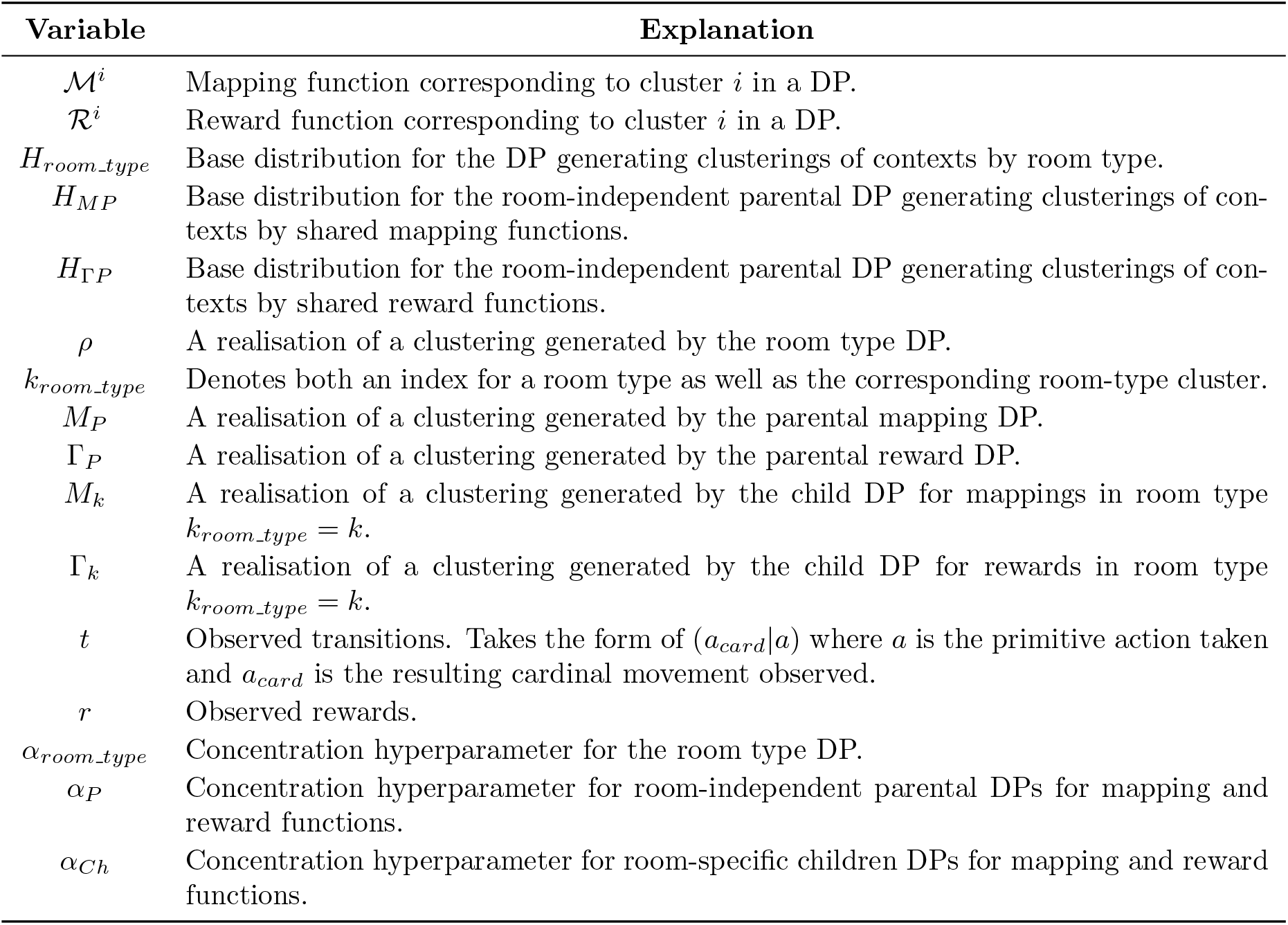
Summary of variables appearing in the hierarchical agent and their explanation.

It is important to note that room clusters in the hierarchical agent offer a generalisation of the purely joint structures defined in the joint agents. Indeed, as a special case, these clusters can comprise specific pairings of ℳ^*i*^ and ℛ^*j*^ (Fig. 3, yellow clusters) – as in joint agents. On the other extreme, the agent may recognise a room cluster where the components are completely unpredictive of each other, and hence clustered independently (Fig. 3, brown cluster). These independent room clusters also permit situations in which some of the ℳ^*i*^ or ℛ^*j*^ clusters can be more popular than others (Fig. 3b). Finally, room clusters can also take on intermediate forms, if a given component is typically paired with a restricted set of other task components (Fig. 3, orange clusters). For instance, in room cluster 4, the dark purple ℳ^*i*^ is always applicable but can be paired with either the red ℛ^*j*^ or the pink ℛ^*j*^. Thus one can consider this like a noisy joint structure.

**Figure 3:**
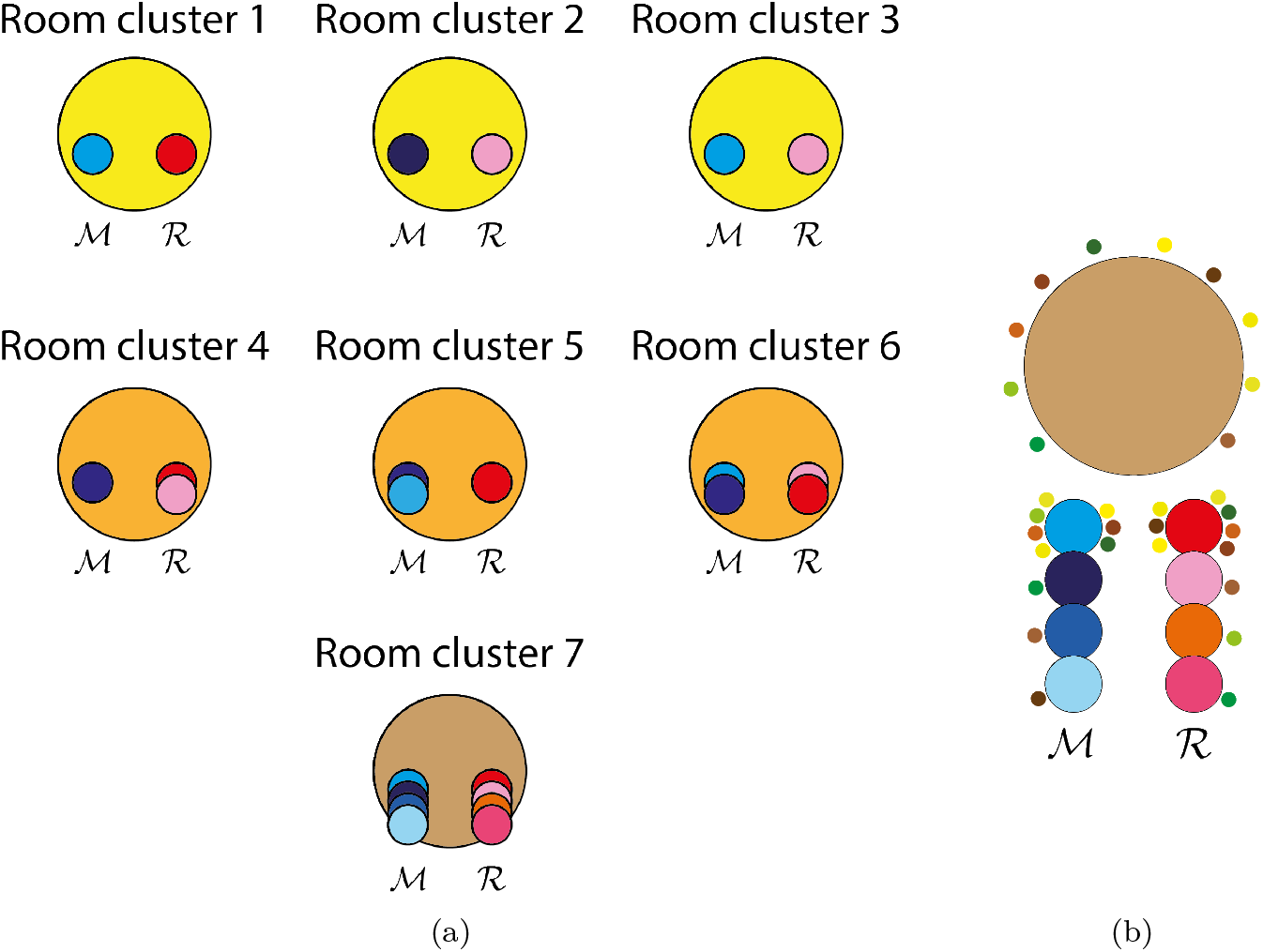
(a) Room clusters (large circles) can capture a variety of structures from purely joint (yellow clusters, wherein a given mapping function is always paired with a corresponding reward function) to those with completely independent components (brown cluster). Orange clusters represent an intermediate case, wherein a given mapping function is typically applied with a small set of possible reward functions or vice versa. Each room cluster contains its own pair of children CRPs for ℳ and ℛ, represented by the chains of small circles, with each small circle representing a single cluster within the CRP. (b) Even within an independent room cluster, one of the mappings or reward functions could be more popular than the others. Here, a variety of distinct contexts (small colored dots) are assigned to this room cluster (large brown circle). Each context is also assigned a ℳ cluster (small colored dots surrounding the ℳ circles) and an ℛ cluster (small dots circles surrounding the ℛ circles). For this cluster, the top blue ℳ cluster has been assigned far more contexts than all the other ℳ clusters; likewise for the top ℛ cluster.

Note that the hierarchical agent confers a potential generalization advantage over the joint clustering agent even within purely joint room clusters. In particular, as we mentioned above, the agent can readily leverage knowledge of familiar individual task components to construct a novel joint structure; only the structure (i.e., the linking from which ℳ goes with which ℛ) would have to be learnt from scratch. As an example, suppose the agent had already learnt room clusters 1 and 2 in Fig. 3 and was adding in room cluster 3, which reuses the blue ℳ^*i*^ from room cluster 1 and the pink ℛ^*j*^ from room cluster 2. Here, the agent can simply transfer these components over, even if the pairing is novel. For instance, in a new context, if the agent sees that pushing key 0 takes it up (and if this transition is unique to a given mapping), then it could immediately infer the keys for the other directions and hence transfer entire mapping functions to new structures. In contrast, the joint agent is unable to support such combinatorial transfer and would have to relearn even familiar components again when they reappear in novel ways.

More broadly, we note that the hierarchical agent subsumes the hypothesis spaces of both the independent and joint clustering agents, making the hierarchical agent more general than either of the two. Indeed, by an appropriate choice of hyperparameters, we can actually force the hierarchical agent to behave exactly like either of the two agents, showing that they are special cases. We discuss this further in the Supplement.

In principle, each DP could have its own concentration hyperparameter *α*_*•*_; but to reduce the model complexity, we have simplified the number of hyperparameters to just three: *α*_*room*_*_*_*type*_ for the room type DP, *α*_*P*_ for all parental DPs, and *α*_*Ch*_ for all children DPs. Also, we found in practice that simply setting *α*_*P*_ to 1.0 and *α*_*room type*_ as well as *α*_*Ch*_ to 0.5, without significant finetuning, was sufficient to achieve good performance in our simulations.

#### 2.2.5 Flat agent

For comparison, we also simulated an agent that does not transfer any prior knowledge and instead constructs a new model for each room. In each new context, it will learn both ℳ^*c*^ and ℛ^*c*^ from scratch. One can think of this as a joint or independent agent where contexts *c* map one-to-one onto clusters *k*; that is, *k* = *c*. Following [14], we call this the *flat* agent (to contrast it with the other agents that are hierarchical, in the sense that they cluster latent states which then define the mapping and reward functions).

### 2.3 Four variants of compositional gridworld environments

To explore (ecologically motivated) situations that might benefit from the different agents, we constructed four versions of our cMDP environment. The first environment involves task structures that are composed independently from the task components, making this environment better suited to the independent agent but not the joint one. Conversely, the second environment involves tasks in which mapping functions are paired with specific reward functions, making it better suited to the joint agent instead. The final two environments involve mixtures of correlational structures that can co-exist in various (and more naturalistic) ways. Although the meta-agent can handle the first two environments (because it arbitrates between independent or joint clustering strategies), we find that only the hierarchical agent is adaptive across all four environments.

#### 2.3.1 Independent environment

We refer to the first environment as the *independent environment* : mapping and reward functions are counterbalanced to give statistical independence between them (Fig. 4a). Two distinct mapping functions are paired with two distinct goal locations, giving a total of four unique pairings and hence four unique contexts. A given mapping is not predictive of a given goal (hence independence).^2^ Contexts appear in random order and are visited a total of four times across the entire environment. Again, contexts (and hence room identity) are observed by the agent. In this environment, an agent that can exploit the tasks’ compositional structure by independently clustering and transferring the reward and mapping functions will exhibit an advantage, as shown by [14]. In contrast, agents that only transfer the entire model (the joint mapping-reward functions) would perform suboptimally when these functions reappear in novel combinations.

**Figure 4:**
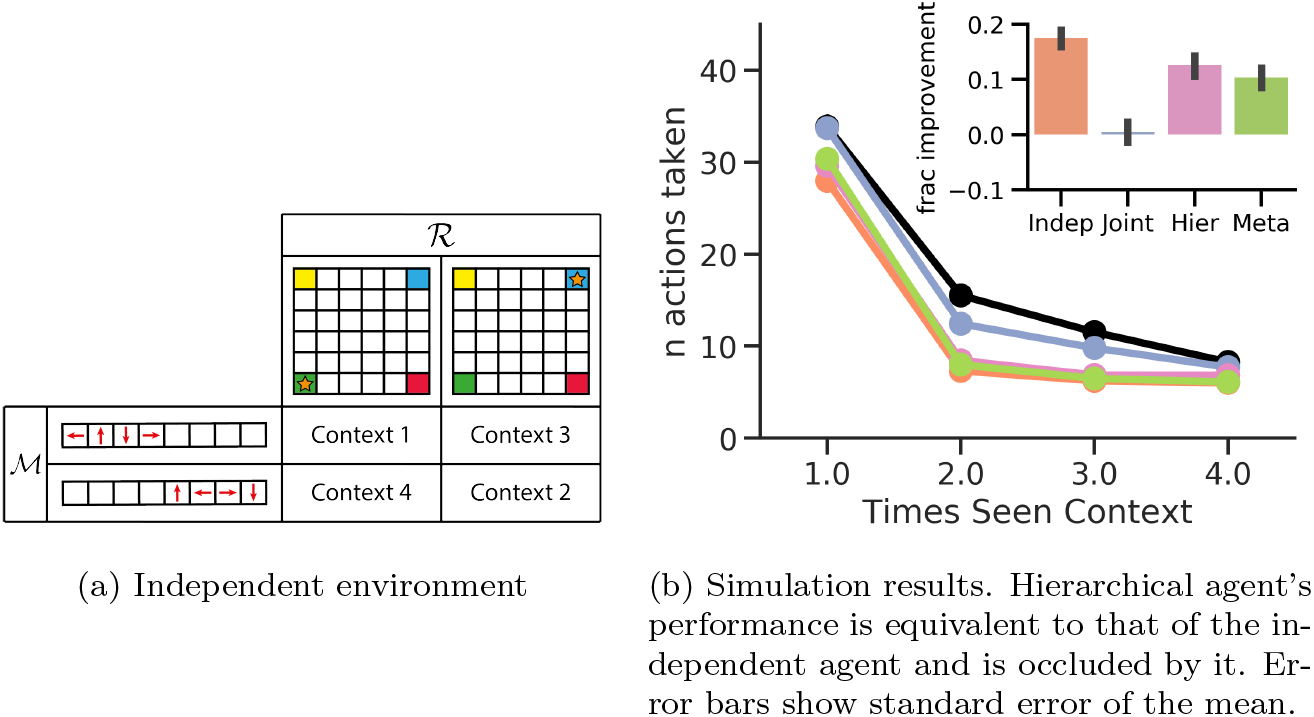
Independent environment. (a) Example of mappings ℳ and goals ℛ (orange star) used in each contexts of the four environments. Goals, mappings, and context numberings, as well as the order of context appearance, were randomised across trials. (b) Average number of steps taken to reach the goal in a context plotted against the number of times the agent has visited that context. Inset shows the average fractional improvement of each agent over the flat agent baseline (black graph in main plot) when visiting the context for the first time (bar colours for each agent has been matched to their counterparts in the main graph). In the main graph, each point is averaged across all distinct contexts and across 150 trials. The first point corresponds to the agent’s average performance when visiting a context for the first time, providing a metric for how effectively the agent leverages prior knowledge. A lower number of actions implies more effective transfer to reach the goal. Likewise, a higher bar in the inset also implies better transfer. Compared to the joint agent, the independent, hierarchical, and meta-agent’s fractional improvement were significantly greater (one-sided Welch’s t-test: *t* = 13.7, 9.4, 7.8; *p* = 2.8 × 10^−42^, 3.8 × 10^−21^, 4.2 × 10^−15^; and *df* = 4758, 4798, 4790, respectively).

Fig. 4b shows how many steps each agent needed, on average, to reach the goal in a context as a function of how many times the agent has seen that context. As expected, the independent clustering agent shows the most effective transfer, needing the fewest actions on average to reach the goal in a novel context, while the joint agent is almost as suboptimal as the flat agent. The independent agent can leverage the knowledge from previous contexts about the likely ℳ and ℛ functions (in contrast to the flat agent) without assuming that knowledge of one is indicative of the other (in contrast to the joint agent). As shown in [14], the meta-agent can also transfer effectively, as its hypothesis space subsumes that of the independent agent. The same property emerges from the hierarchical agent here, confirming that it can also easily identify independent structure even though (unlike the meta-agent) it is not given this structure as one of two *a priori* hypotheses.

#### 2.3.2 Joint environment

We next consider an environment where mappings are paired in a one-to-one manner with goals to make one entirely predictive of the other (Fig. 5a). We refer to this as the *joint environment*. Here, we use four distinct mappings and four distinct goals, but to get the one-to-one correspondence, each mapping is paired with only one of the goals, giving four unique mapping-goal pairings. Each pairing now appears as two distinct contexts to test the ability of structural transfer from one context to another that uses the same mapping-goal pairing. Contexts again appear in random order and are visited a total of four times each. This environment will favour an agent that can exploit the one-to-one correspondence between mappings and goals, and an agent that learns goals and mappings as a single structure will outperform one that treats them as independent components [14].

**Figure 5:**
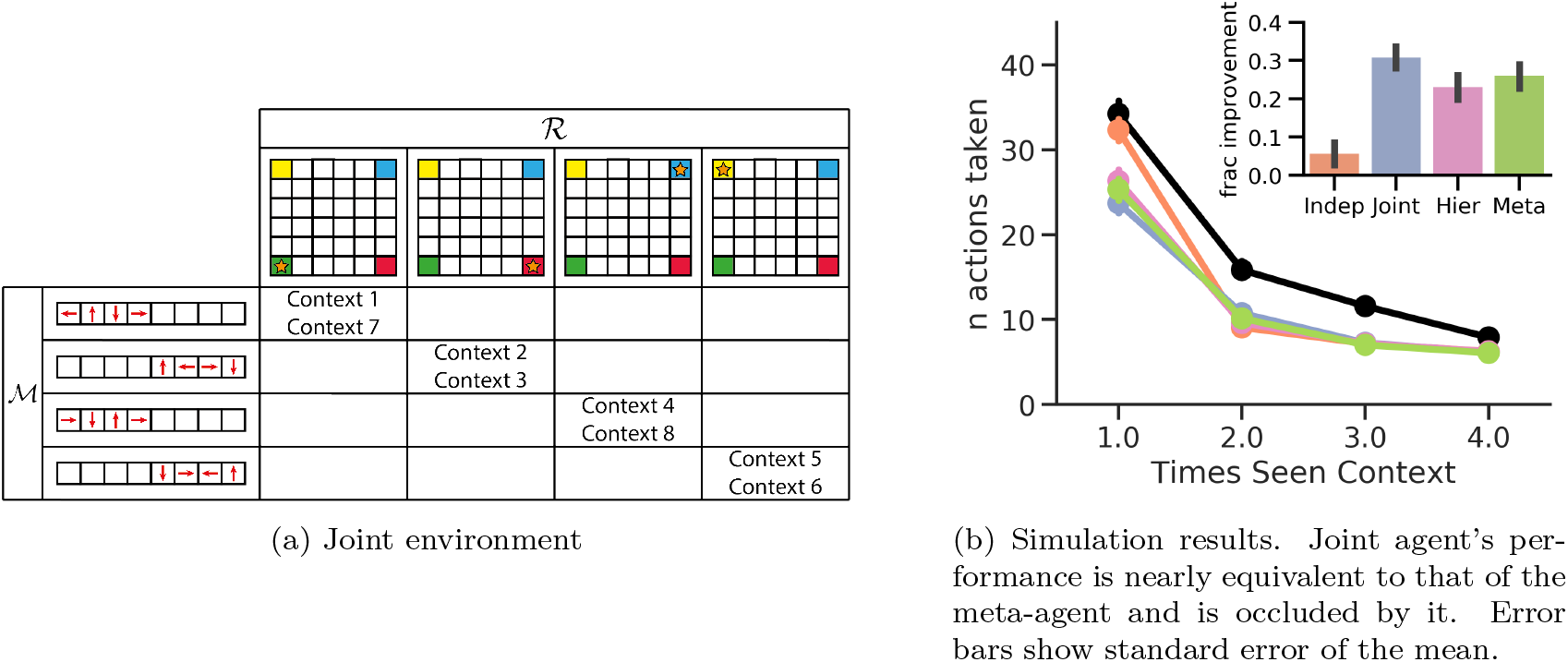
Joint environment. (a) Mappings ℳ and ℛ rewards are now paired in a one-to-one manner. Goals, mappings, context numberings, and order of context appearance are again randomised across trials. (b) Compared to the independent agent, the joint, hierarchical, and meta-agent’s fractional improvement were significantly greater (one-sided Welch’s t-test: *t* = 12.2, 7.9, 9.5; *p* = 1.0 × 10^−33^, 1.6 × 10^−21^, 2.3 × 10^−15^; *df* = 2397, 2342, 2377, respectively).

Fig. 5b shows this to be the case. The joint clustering agent transfers most effectively in this environment, while the independent agent is almost as suboptimal as the flat agent. The hypothesis spaces of both the meta-agent and the hierarchical agent subsume that of the joint agent as well, so their near-optimal performance suggests that they are successfully converging on hypotheses that capture the joint statistics between ℳ and ℛ.

#### 2.3.3 Mixed environment

So far, only the meta and hierarchical agents have been adaptive across both environments. But these environments are far from naturalistic. Independent and joint statistics represent the two extremes: in one case, the mappings and goals are in a strict one-to-one correspondence, and in the other, they are completely independent of each other. The next two environments we consider are arguably more representative of naturalistic tasks. Both require recognising that several different types of generative structures can exist alongside each other within the same environment.

Indeed, it is often the case that special joint structures will exist alongside independent ones. For instance, in navigation, one might always take a boat to go waterskiing, but be equally likely to drive, boat or bicycle to reach other goals. In music, one might always play bluegrass on a banjo but be equally likely to play classical, jazz or folk on other stringed instruments. In these cases, the waterskiing-boat and bluegrass-banjo pairings comprise joint structures that co-exist with otherwise independent structures. Our next *m*ixed environment (Fig. 6a) explicitly combines joint structures alongside independent ones. In Fig. 6a, contexts 1-3 and 4-6 each combine one mapping with exactly one goal, making these contexts with transferable joint statistics. The remaining contexts, contexts 7-16, have independent statistics as the goals are not predictable from mappings. They combine five additional mappings with two additional goals, giving a total of 10 contexts such that every possible goal-mapping combination appears exactly once. As before, contexts appear in random order and are visited four times.

**Figure 6:**
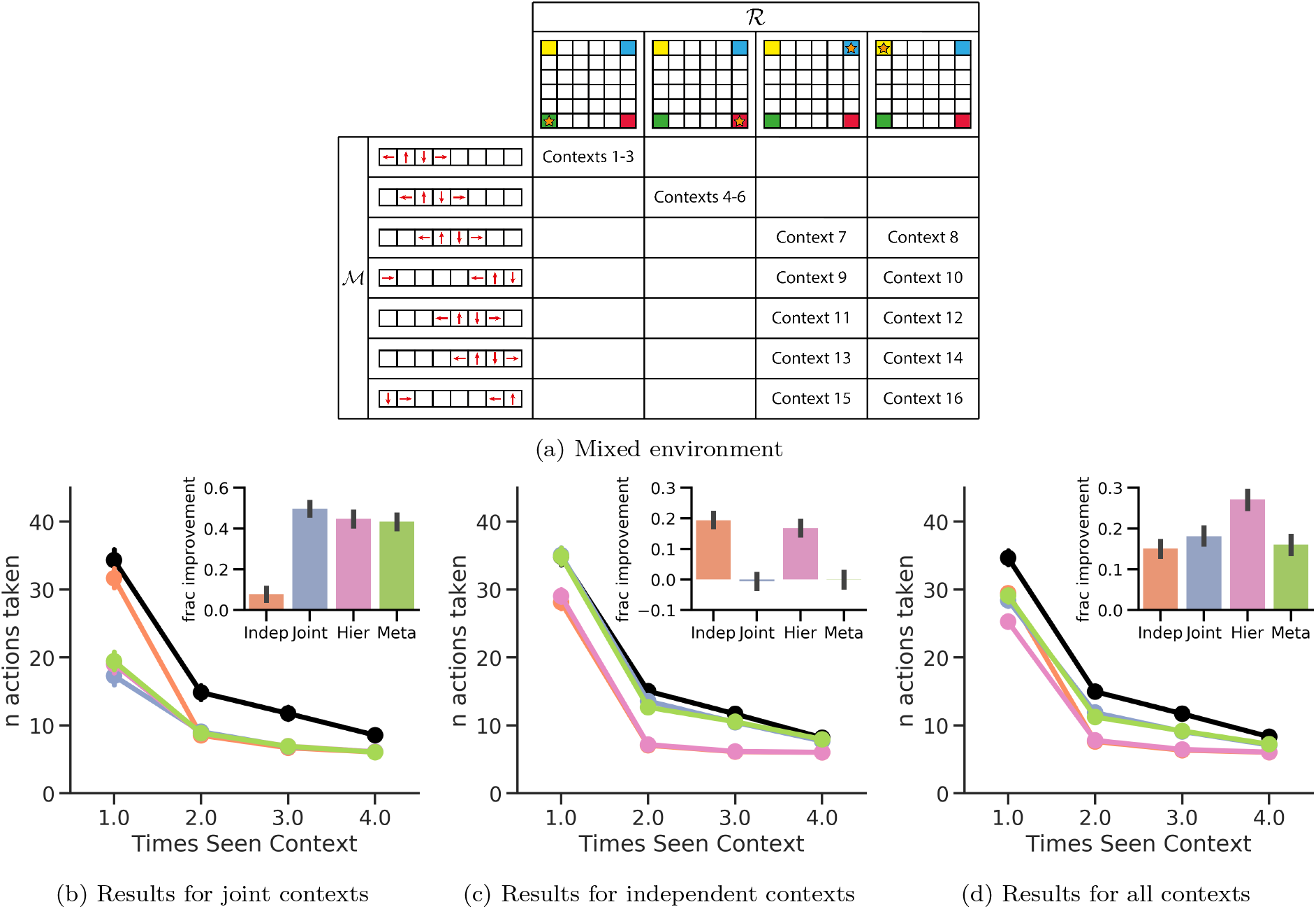
Mixed environment. (a) The environment consists of two sets of joint contexts (contexts 1-3 and 4-6) alongside a set of independent contexts (contexts 7-16). (b) In the joint contexts, the joint, hierarchical, and meta-agent’s fractional improvement were significantly better than the independent agent’s (one-sided Welch’s t-test: *t* = 18.6, 15.1, 15.3; *p* = 2.5 × 10^−71^, 6.6 × 10^−49^, 3.8 × 10^−50^; *df* = 1795, 1738, 1780, respectively). (c) In the independent contexts, the independent and hierarchical agent’s fractional improvement were significantly better than the joint agent’s (one-sided Welch’s t-test: *t* = 11.9, 10.4; *p* = 3.2 × 10^−32^, 5.0 × 10^−25^; and *df* = 2993, 2993, respectively). (d) Across all contexts, the hierarchical agent showed the most effective transfer. Compared to the next best agent (the joint agent), it had a significantly greater fractional improvement (one-sided Welch’s t-test: *t* = 6.0, *p* = 1.3 × 10^−9^, and *df* = 4789) indicating it was the most effective at transferring structural knowledge when visiting novel contexts. All error bars show standard error of the mean.

Intuitively, the joint agent shows the most effective transfer in new joint contexts (Fig. 6b), while the independent agent shows the worst; this situation is reversed in the independent contexts (Fig. 6c). The meta-agent approximates the joint agent’s behaviour in this environment. Importantly though, only the hierarchical agent is adaptive across all contexts (Fig. 6d), almost matching the joint agent’s performance in the joint contexts and the independent agent’s performance in the independent contexts. Thus, the hierarchical agent successfully learns and exploits room types that separate out the two types of joint contexts and the independent contexts from each other.

#### 2.3.4 Conditionally independent environment

Another common situation in naturalistic settings is for structural statistics to be conditionally independent. For instance, in the city, the choice of transport method may be completely independent of the choice of destination. Along waterways, the choice of transportation, whether it be speedboat, canoe, kayak or even swimming, can also be independent of the choice of destination. However, when aggregated overall, we only have conditionally independent task statistics because choosing land transportation methods will still be predictive of city rather than water destinations, and vice versa.

In the final *conditionally independent environment*, we consider such structure. Contexts are organised into two sets (Fig. 7a), each with four mappings and two goals. However, the mappings and goals in the two sets are mutually exclusive: mappings that appear in one set do not show up in the other and vice versa; likewise for the goals. Within each set, mappings and goals have independent statistics; that is, each of the four mappings in a set gets paired with each of the two goals to give eight contexts each. So the environment actually consists of two “independent structures” alongside each other – that is, a conditionally independent structure.

**Figure 7:**
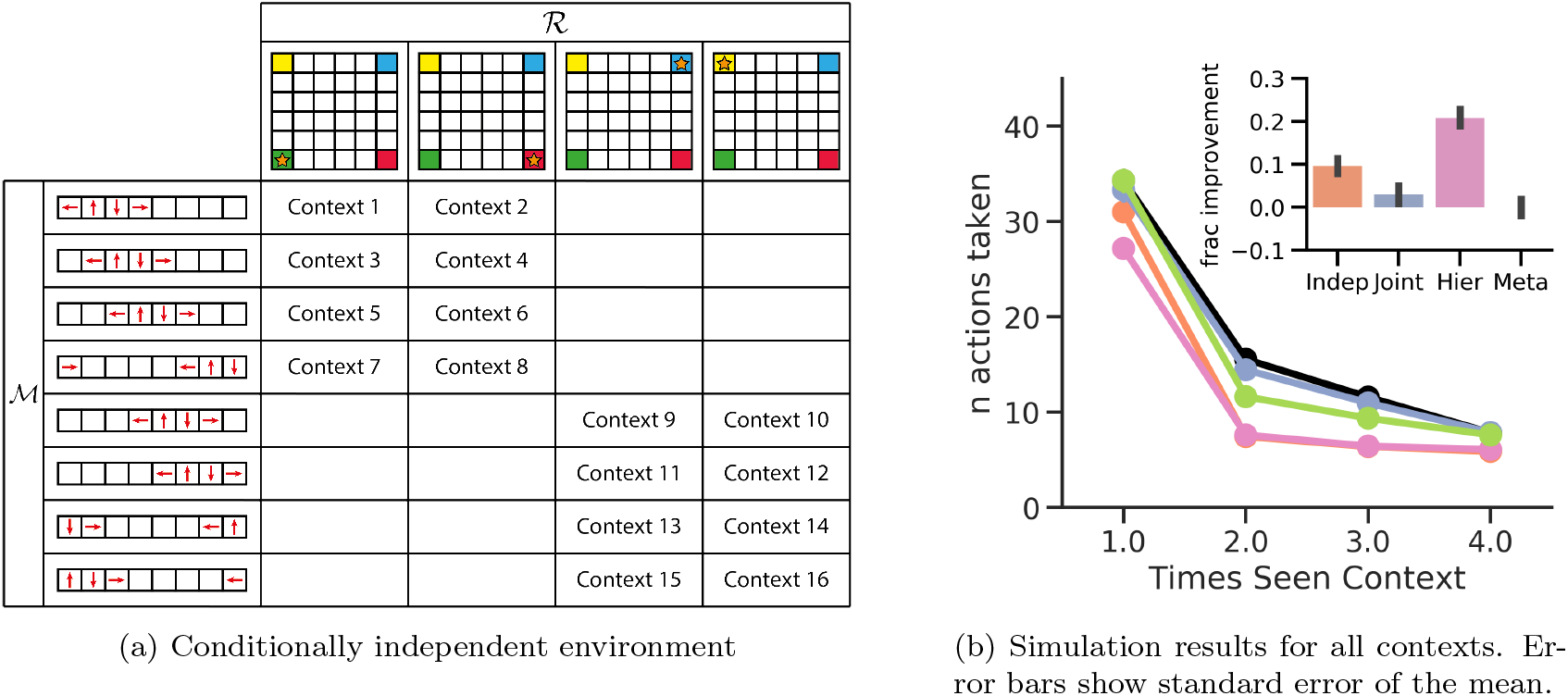
Conditionally independent environment. (a) Contexts are organised into two sets that use mutually exclusive mappings and goals. In each set, contexts have independent statistics, making the environment overall conditionally independent. (b) In new contexts, the hierarchical agent showed the best transfer of structural knowledge, as demonstrated by its significantly higher fractional improvement compared to the next best agent, the independent agent (one-sided Welch’s t-test: *t* = 8.2, *p* = 1.7 × 10^−16^, and *df* = 4766).

Because of the conditionally independent structure, an independent agent cannot be optimal in this environment because it will try to mix mappings in one set with goals in the other. Naturally, the independent statistics implies that the joint agent cannot be optimal either. In fact, only the hierarchical agent can be adaptive in this environment; it is the only agent capable of recognising the conditionally independent structure and will learn to represent it using two distinct room types that do not mix goals and mappings between rooms (Fig. 7b).

## 3 Scaling compositionality to hierarchical environments

Above we showed that the hierarchical agent can discover and transfer structures across a wider range of task statistics, and is thus more expressive and general than any of the other agents. But in the environments we considered, agents had to discover just a single goal and transition function in a given context. In contrast, the compositional structure of naturalistic tasks are often more complex and hierarchical, with goal/subgoal structures to them. In such cases, compositional generalisation becomes even more critical. Indeed, as structures increase in complexity, the number of possible structures explodes combinatorially, making naive memorisation of joint structures unsustainable over an increasing number of tasks. Furthermore, most task structures that an agent encounters across its entire lifetime will likely appear only once in the exact same way, further weakening the transfer value of exact structural memorisation. Rather, smaller substructures are more likely to recur, and an agent that can learn and transfer and recompose combinations of such substructures in new ways will likely be more adaptive. In this section, we demonstrate the greater expressivity of hierarchical agents to tasks with a more hierarchical, compositional structure.

The tasks we consider below are structured hierarchically with goal/subgoal structure, a situation commonly studied within the framework of *hierarchical reinforcement learning* [24, 12]. *As a real world example*, let us return again to our earlier navigation example. Suppose a person were visiting a city for the first time. She may hope to do several things during this trip, such as sightsee, sample the local delicacies, and enjoy the local nightlife. The success of the trip depends on how many of these activities she manages to do, but each activity takes place at a different locale around the city. Thus, the overall trip can be broken down hierarchically into first deciding the sequence in which to do things, and then for each activity, the subgoal of travelling there. The ordering of activities, however, can often be constrained by what one knows about cities and activities in general. For instance, one will generally want to have breakfast in the morning, to sightsee during the day, and to visit bars, night markets, and nightclubs in the evening. Additionally, the mode of transport can depend not only on the destination but also on higher-order contexts like the city itself, constraining modes of public transport that are most effective for given destinations. As another example, suppose one were travelling to a foreign land to conduct some business. At the start of a meeting in a conference room, the businessperson will realise that, in this context, they should first be introducing themselves and thanking their hosts. This decision about the general way to behave is inherited from their experience across many other business meetings in other cities. But the specific social norms that apply as to *how* to introduce, thank and greet the hosts should be informed from the local culture, and they would need to adjust their greeting “policy” accordingly, whether it be to shake hands or bow or, in the context of a global pandemic, bump elbows. Hence, the statistical dependence between task components (e.g., between the choice of transport and the destination in the first example, or the choice of social behaviors in the second) now depends on a hierarchy of contexts, including not just local contexts but also higher-order contexts like the city or country that everything takes place in.

In these cases, solving the overall task requires implementing a hierarchical policy, which can usually be broken up into subpolicies for the different subtasks, which can in turn be recombined in novel ways to quickly solve novel macro tasks. Developing approaches to learn such hierarchical policies has been the focus of hierarchical reinforcement learning [24, 12, 23, 25, 26]. However, such approaches have not considered how to transfer distinct components of knowledge across levels of the hierarchy depending on both global and local contexts. Additionally, these approaches tend to focus on model-free learning of both the subpolicies and overall macro policy, whereas our approach is motivated more from the perspective of model-based reinforcement learning where the agent must instead first determine the correct compositional model of the task and then use it to solve the task through model-based planning. In the next series of tasks, we explore the possibility of scaling our approach up to tasks with such hierarchical, compositional structure.

### 3.1 Castle – dungeon problems: Sequences of hierarchical gridworld rooms

We now consider environments where the agent must navigate a series of hierarchically structured rooms in a castle (Fig. 8). Each room has an upper level and three sublevels (reminiscent of “underground” levels in the “Super Mario Bros” video game or “dungeons” in “Zelda”), and each level is a gridworld. The agent starts in the upper level and must navigate to a sequence of four doors in the correct order (Fig. 8b). The first three doors leads the agent to each of the sublevels (orange arrows in Fig. 8a), where the agent has to navigate to a target location to return to the upper level. When the agent reaches the fourth door, it finishes this room “level” and advances to the next room (or ends the task if it is the final room), but only if the agent has first completed the first three sublevels in that room. To avoid having a non-Markovian reward function, the agent’s state information in the upper level includes how far along has it progressed in the door sequence.

**Figure 8:**
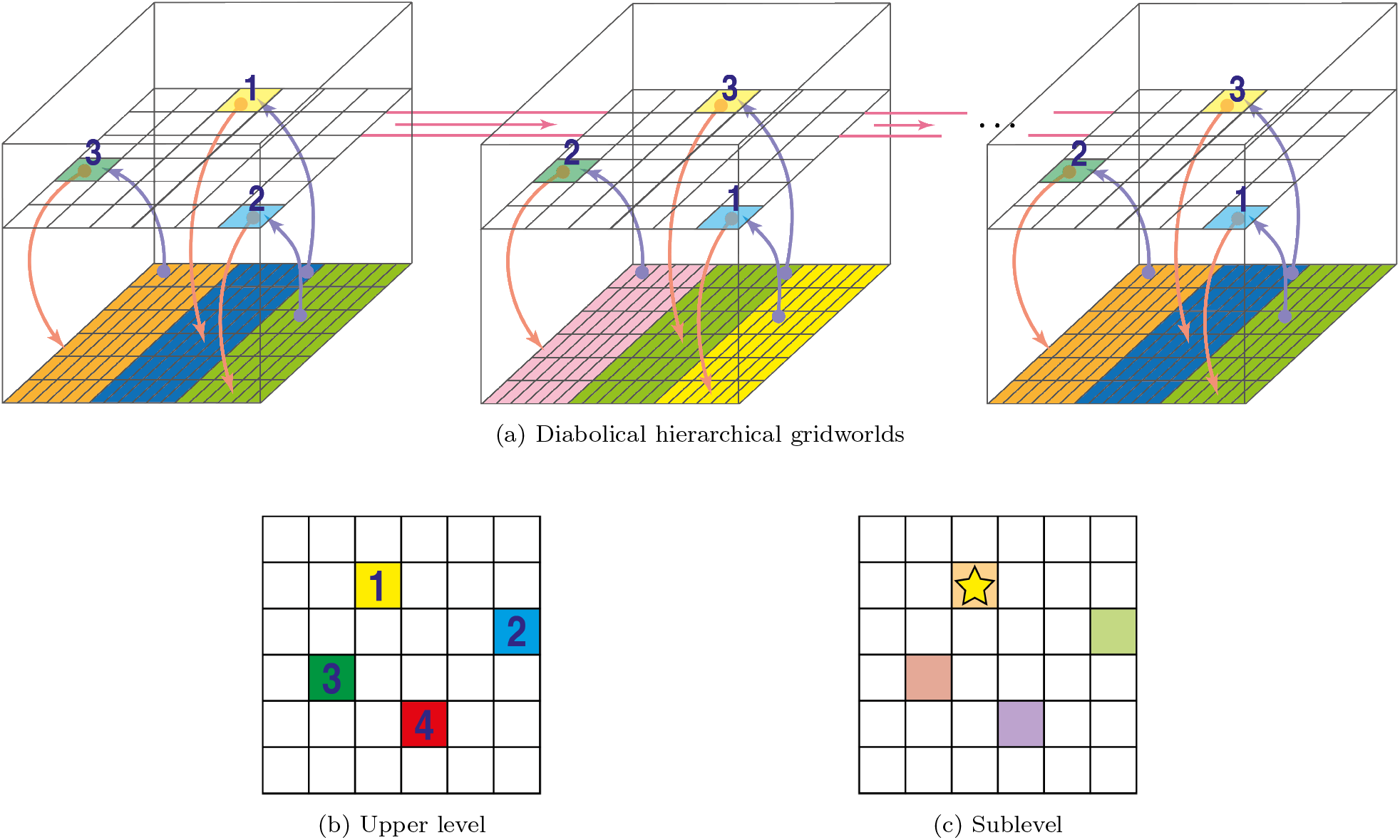
Hierarchical gridworlds environment. (a) The agent has to navigate through a series of rooms in a castle. The agent starts off in a room’s upper level and must pass through doors to visit each of the room’s three sublevels. To solve a room, the agent must visit the three sublevels in the correct order (indicated by the numbered sequence over the door squares). The agent enters a sublevel by stepping on the correct coloured square, which leads the agent into the sublevel (orange arrows). Upon successfully completing a sublevel, the agent is taken back to where it left off in the upper level (blue arrow). After visiting all sublevels, the agent can pass through a fourth door to advance to the next room (pink bridge) or end the task if in the final room. Each upper level has the same four doors (b) but the order in which they must be visited changes from room to room (numbered sequence). Each sublevel has four possible goals (c), but only one of these is correct (star) and leads back to the upper level. Which goal that is changes from sublevel to sublevel. Each upper level or sublevel also has its own mapping function (indicated for sublevels by the colour scheme in (a)), and this also changes between upper and sublevels; some of these repeat will repeat though and can therefore be reused to facilitate transfer (repeated colours in the sublevels). In the example here, the final room repeats the same set of sublevel mappings as in the first room but the same upper level door sequence as in the second room. In the diabolical version of this environment, if the agent attempts the wrong door (i.e., it attempts a door out of order) or if it goes to the wrong sublevel exit, it is sent back to the beginning of the task, making mistakes extremely costly.

Each sublevel itself (Fig. 8c) comprises another room with four exits, but where only one is correct and leads the agent back to the upper level (blue arrows in Fig. 8a).

We consider two versions of this hierarchical environment. The first is essentially a modification of the “diabolical rooms” environment from [14] to include hierarchical structure. In the original environment, the agent had to navigate through a sequence of compositional gridworld rooms (like those studied in the previous section), where the mappings and reward function (goals; i.e., which door to enter to reach the next room) in any given room might be similar to one of the previous rooms. But the diabolical nature of the environment refers to the additional challenge whereby if the agent goes to the wrong door in any room, it will return back to the start of the first room, making mistakes extremely costly. There is thus a large advantage towards agents that can transfer prior knowledge compositionally; this advantage grows with the number of rooms and the size of each one [14]. Here we significantly enhance the challenge imposed on the agent further: the environment is both diabolical and hierarchical. Thus, any mistake (within either upper or sub levels) will send the agent back to the beginning of the entire environment, again putting pressure on the agent to leverage prior knowledge about rooms and sublevels effectively.

In second version of the environment, to better examine transfer in individual rooms, we remove the diabolical part of the environment. The agent can keep exploring a room until it finds the correct exit, whereupon it enters the next room (or the task ends if it is the final room). The set-up is similar to the non-hierarchical environments studied earlier. Each room is now associated with a context, each context is visited four times across the entire environment, and contexts are visited in random order.

All environments used a total of 16 hierarchical rooms, giving a total of 64 gridworlds (including the upper level and three sublevels for each room). For both diabolical and non-diabolical versions, we manipulated the generative statistics of the environment to generate three environment types:

1. Independent statistics. This environment used a total of four distinct mappings, four distinct door sequences, and all four possible sublevel exits. To generate independent statistics, the applicable mappings, door sequences in the upper rooms, and exits in the sublevels were all randomly paired with each other; that is, knowledge of the mapping or reward function in any level was not informative of the others in that level or the other levels. Nevertheless, to ensure that there was a benefit to leveraging prior knowledge, some mappings, door sequences, and sublevel exits were more prevalent than others. Thus an agent could improve performance by, for example, reusing popular mappings in a room or sublevel that it had encountered in combination with a different door sequence.
2. Joint statistics. In this case, the upper level mapping was entirely predictive of both the door sequence and the mappings and exits in each of the room’s sublevels. Again, some of the mappings, and by extension the doors and sublevels they predict, were more prevalent than others. Transfer could be improved by recognizing and reusing this joint association.
3. Mixed statistics. This environment had three rooms with strictly joint statistics, four where the upper level mappings were predictive of doors and sublevel exits but not sublevel mappings, and nine with completely independent statistics between all mappings and reward functions.

### 3.2 Agent architectures for the hierarchical environments

We ran a flat agent (as in the earlier simulations, the flat agent had no clustering or generalization abilities, but was still a model-based agent), an independent clustering agent, and a hierarchical agent on the hierarchical environment.^3^ As above, the agents are endowed with knowledge of the general structure of the tasks: in this case, that there are rooms consisting of upper levels and sub-levels, with goals and mappings to be learned for each. We generalized the agents described in the earlier sections to allow for clustering of these higher level structures. Indeed, our hierarchical environments can be reduced to the non-hierarchical ones if the first door in the sequence for each upper level led directly to the next room; our extended agents for the hierarchical environments could then be directly run on the non-hierarchical ones.

The independent agent clusters each task component independently of the others (Fig. 9). One DP is used to cluster mappings across the entire environment, and another is used to cluster exits for all sublevels. The agent also maintains distinct DPs to cluster reward functions for each position in a door sequence.

**Figure 9:**
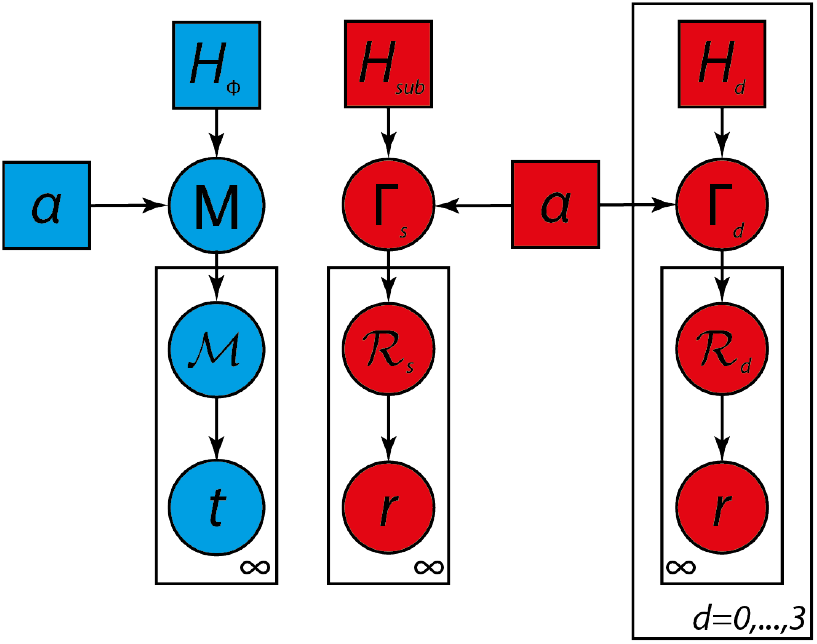
Independent clustering agent for the hierarchical environment. The blue DP clusters mappings, with the base distribution *H*_ℳ_ generating clusterings *M* of mappings ℳ, which in turn generates the observed transitions *t*. The red DPs independently cluster goals for doors and sublevel exits. A base distribution *H*_*sub*_ generates clusterings Γ_*s*_ of exit reward functions ℛ_*s*_ for sublevels, which in turn generates observed rewards *r*. A set of four independent DPs, one for each door sequence position, generate clusterings Γ_*d*_ of door reward functions ℛ_*d*_ from base distributions *H*_*d*_. ℛ_*d*_ in turn generates observed rewards *r*.

The hierarchical agent, in contrast, can simultaneously represent independent and joint structure, as it clusters task components at multiple levels of hierarchy (Fig. 10). Just as in the non-hierarchical environment, it maintains one clustering of room clusters (Fig. 10, dark green nodes) that capture higher-order structural information, though these now describe the structure of hierarchical rooms. In addition to mapping and reward functions, the hierarchical agent recognises two additional structural components that go into composing hierarchical rooms: the sublevel structures and upper level door sequences. These components are composite substructures that hierarchical rooms can reuse, and are themselves decomposable into smaller, more primitive structural components. Note that, unlike previously, this composition gives the agent the ability to transfer structural knowledge not just at the macroscopic room-level, but also at other levels of the structural hierarchy; indeed, substructures like sublevels and door sequences are more likely to recur than entire room structures, and the agent can recognise and reuse these substructural motifs across different rooms whenever they occur. As with the previous hierarchical agent, this hierarchical agent also clusters each structural component in a hierarchical manner:

**Figure 10:**
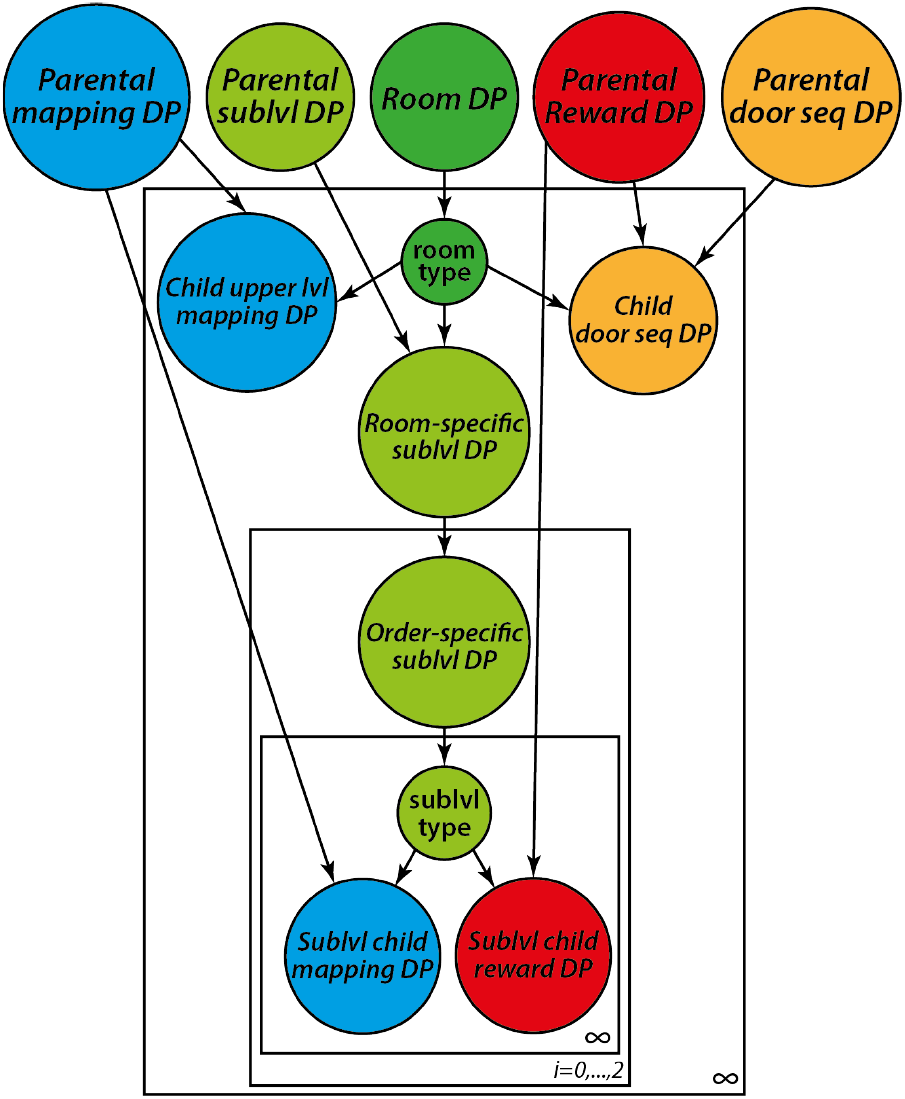
A schematic of the hierarchical agent’s generative model for the hierarchical environment. This consists of a DP for clustering hierarchical rooms by room type (dark green nodes), as well as several hierarchical DPs for clustering structural components that make up rooms, including hDPs for mapping functions (blue), sublevel structures (lime green nodes), reward functions (red nodes), and door sequence structures (orange nodes). Sublevel structures and door sequence structures are composite substructures that can appear across a variety of different hierarchical rooms, and thus the hierarchical agent will try to learn and transfer these substructural motifs. Sub-levels are composed of mapping and reward functions while door sequences are composed of four reward function, one for each position in the sequence. A full graphical model can be found in the Supplement.

1. *sublevel structures* (Fig. 10, inner-most plate) are like the room structures of the previous hierarchical agent in that they consist of one DP for clustering sublevel mappings (blue node) and a separate DP for clustering sublevel reward functions (red node), making them composite structures. These structures are clustered in a hierarchical manner (Fig. 10, lime green nodes) starting from a parental DP that clusters sublevel structures across the entire environment followed by a room-specific DP that clusters sublevels independent of their order in the room; this latter clustering induces a distribution over possible sublevel structures specific to that room. This is then followed by three further order-specific DPs (Fig. 10, middle plate) that induce different distributions over sublevel structures according to the sublevel’s position in a room (i.e., the first DP models the distribution over different possible structures that appear as the room’s first sublevel, the second DP for the room’s second sublevel, etc).
2. door sequences can be regarded as a composition of four reward functions, one for each door in a sequence. *Door sequence structures* consist of four DPs that cluster reward functions, one for each position in a door sequence (not shown in figure for conciseness). These structures are intended to model statistical correlations between door locations in a sequence, with each DP modelling the distribution over reward functions for the corresponding sequence position. The structures are themselves clustered in a hierarchical manner (Fig. 10, orange nodes) starting with a parental DP that clusters sequence structures across the entire environment followed by a room-specific DP that captures the distribution over sequence structures for that particular room type.
3. mapping functions, as in the previous hierarchical agent, are clustered in a hierarchical manner (Fig. 10, blue nodes) starting with a parental DP independent of any room followed by several children DPs inside the room structures. One of these models room-specific distributions over possible mapping functions for the room’s upper level. The others also model distributions over possible mapping functions for the sublevels.
4. reward functions are also clustered in a hierarchical manner (Fig. 10, red nodes), as in the previous hierarchical agent. Again, hierarchical clustering starts with a parental DP independent of the rooms followed by several children DPs inside room structures. Several of these lie within the door sequence structure described above, while the others lie within the sublevel structures. Each models distributions over possible reward functions conditioned, respectively, on the door position within a door sequence structure or on the type of sublevel.

Note that the hierarchical clustering of all structural components always starts with a parental DP that is independent of any hierarchical room structure, allowing for the factorisation of task components so that they can be recomposed in novel combinations inside novel room structures. While there are alternative generative models for this environment, based on different hierarchies that the hierarchical agent could have used, our primary interest was not to determine the most optimal architecture but rather to demonstrate that endowing a clustering agent with hierarchical structure enhances expressivity because of the potential for compositional transfer at different levels of the hierarchy.

Finally, we note that for this agent, it was sufficient to set all concentration hyperparameters {*α*} to 1.0 to obtain good performance.

### 3.3 Results for the diabolical environments

In all three diabolical environments, the independent and hierarchical agents greatly outperformed the flat agent baseline, needing far fewer steps on average to solve these environments (Fig. 11), indicating that they successfully leveraged experience from previous rooms to solve new ones more quickly. Moreover, the hierarchical agent significantly outperformed the independent agent, as it was able to exploit higher order statistical structures in the rooms, while retaining the independent components. Surprisingly, the hierarchical agent even outperformed the independent agent in the environment with independent statistics. As we will see below, this improvement was simply due to imperfect independence of the generative statistics: unlike in the non-hierarchical task, the combinatorial explosion in complexity arising from the multiple task components made it difficult to ensure complete independence of task statistics within only a limited number of rooms. The better performance of the hierarchical agent shows that it is sensitive enough to pick up on any residual structure that may lurk within the task statistics.

**Figure 11:**
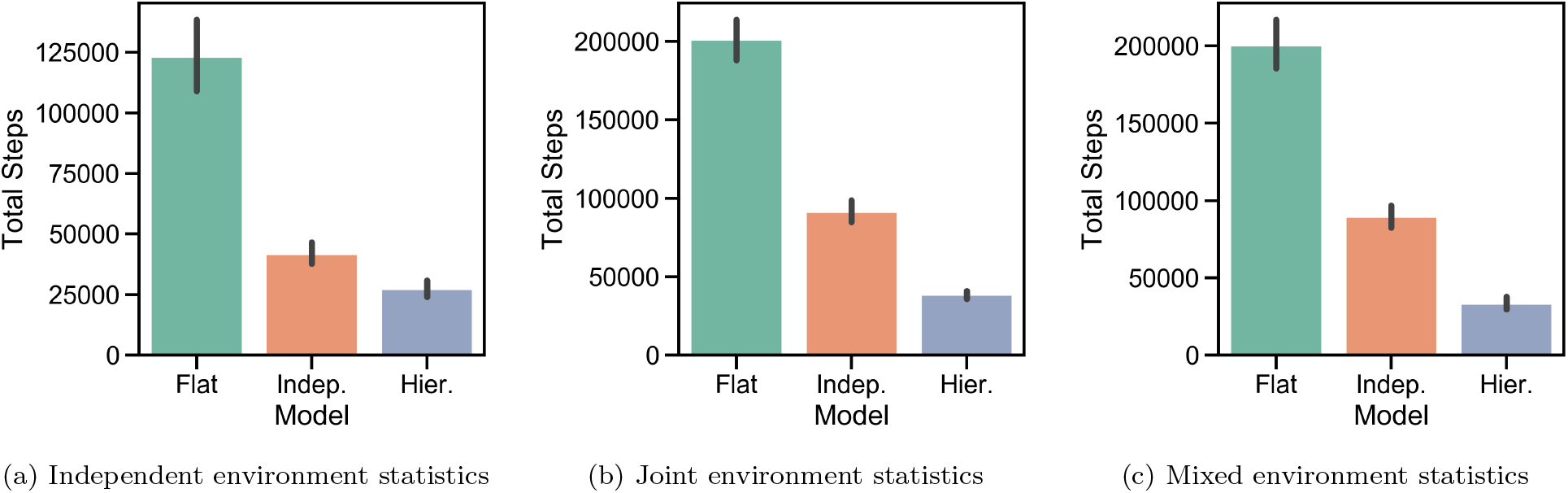
Hierarchical diabolical rooms environments. Total number of steps each agent needed to solve each environment. Results are averaged across 50 runs; errorbars indicate standard error of the mean. The hierarchical agent outperformed the independent agent in all three environments (one-sided Welch’s t-test: *t* = 4.9, *p* = 1.7×10^−6^, *df* = 86 for independent environment; *t* = 13.5, *p* = 3.0 × 10^−20^, *df* = 61 for joint environment; *t* = 12.9, *p* = 1.7 × 10^−20^, *df* = 71 for mixed environment). The independent agent, in turn, outperformed the flat agent (one-sided Welch’s t-test: *t* = 10.3, *p* = 4.4 × 10^−15^, *df* = 59 for independent environment; *t* = 14.3, *p* = 1.9 × 10^−23^, *df* = 75 for joint environment; *t* = 12.5, *p* = 5.2 × 10^−20^, *df* = 71 for mixed environment).

If the hierarchical agent performs better than the independent agent, it must be able to predict the correct goal whenever it sees a gridworld room for the first time. Figs. 12a-b show the fraction of times each of the agents predicted the correct goal in the first opportunity to do so (i.e, for a novel position in a door sequence or the correct exit for a novel sublevel). We see that, as expected, the flat agent’s performance is near chance level of 0.25, but the independent and hierarchical agents’ performances are much higher. Notably, the hierarchical agent’s better performance is driven by its ability to predict the next door in a sequence; Figs. 12c-f show that this advantage only emerges from the second door onwards. This result suggests that the hierarchical agent is exploiting residual structure in the door sequences.

**Figure 12:**
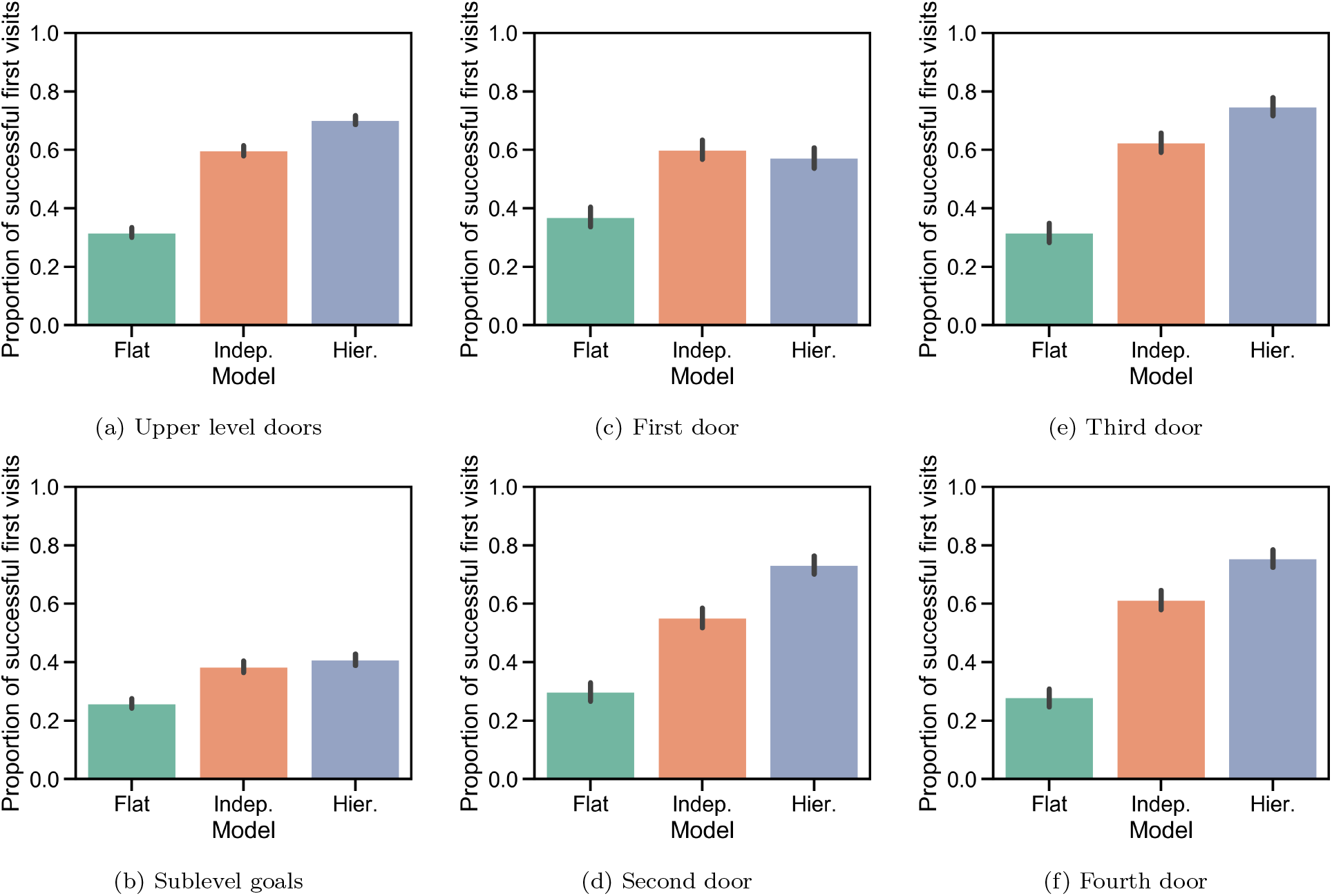
Probability of successful first visits. (a)-(b): The hierarchical agent significantly outperformed the independent agent when predicting doors but not sublevel goals (one-sided Welch’s t-test: *t* = 8.8, *p* = 9.3 × 10^−19^, *df* = 6331 for upper level doors; and *t* = 1.6, *p* = 0.053, *df* = 4759 for sublevel goals). (c)-(f): For the upper level doors, the hierarchical agent significantly outperformed the independent agent when predicting the second, third, and fourth doors (one-sided Welch’s t-test: *t* = −1.1, *p* = 0.14, *df* = 1586 for door 1; *t* = 7.6, *p* = 2.1 × 10^−14^, *df* = 1570 for door 2; *t* = 5.4, *p* = 4.7 × 10^−8^; *df* = 1572 for door 3; and *t* = 6.1, *p* = 5.0 × 10^−10^, *df* = 1566 for door 4).

To better understand this, we quantify the information theoretically available to an agent about structure between task components. Let us suppose the agent has already seen the following sequences of mappings and doors in the upper levels,

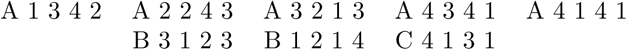

where letters denote different mappings and numbers denote doors. When an agent enters a new room, it will learn the mapping first before it gets any goal information. If it learns that the mapping is A, it will not be able to tell whether the first door will be 1, 2, 3, or 4, as all possibilities follow mapping A. If it then learns the first door is 1, 2, or 3, it can immediately infer the rest of the sequence, since knowing any of these is sufficient to narrow down the possibilities to just one unique sequence. But if it sees door 4 instead, it will need to learn the identity of the next door before it can uniquely determine the rest of the sequence. Alternatively, if the agent learned the mapping was B, it would only need to know the identity of the first door before it could figure out the entire sequence. And if it learned the mapping was C, it could immediately infer the entire sequence right away, although such a coincidence of a mapping paired with just one door sequence was highly unlikely in our environments. The amount of information that a partial sequence *s* tells an agent about the complete sequence can be quantified using the Shannon information

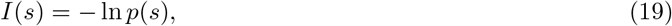

where the probability is computed with respect to the dictionary of all sequences. Since dictionaries can be of varying sizes across different simulations, we normalise this with respect to the total information obtained when the entire sequence is learned; that is,

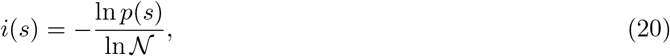

where 𝒩 is the total number of unique sequences.

To see whether the hierarchical agent was exploiting residual regularities in the door sequences left in the independent task, we performed several tests on the fractional information content of the partial sequences. We divided the partial sequences into two sets, one where the hierarchical agent succeeded at guessing the correct door but the independent agent failed, and one where either the independent agent succeeded or the hierarchical agent failed (that is, where the hierarchical agent was not more successful than the independent agent). We found that when the hierarchical agent was more successful, the fractional information content of the partial sequences was significantly higher (one-sided Welch’s t-test: *t* = 9.8, *p* = 5.7 × 10^−22^, *df* = 911). We also ran a logistic regression to predict the probability that the hierarchical agent will be more successful at guessing the door from the fractional information available (Fig. 13). A likelihood ratio test showed that this model was significantly better than one where the probability was the same across all levels of fractional information (*p* = 2.99 × 10^−18^). These results indicate that the agent is likely exploiting structure in the door sequences, which it may do by having more joint-like representations in its door sequence clusters. As we discussed above, most naturalistic tasks of sufficient complexity will rarely repeat in the exact same form twice within an agent’s lifetime; instead, one is more likely to find substructural motifs repeating across different structures. The finite dictionary of sequences used in this environment makes that even more the case here, so an agent that can learn and reuse repeated joint sequences will be more optimal. Our results indicate the hierarchical agent is doing this as evidenced by its better ability to predict the next door in a sequence.

**Figure 13:**
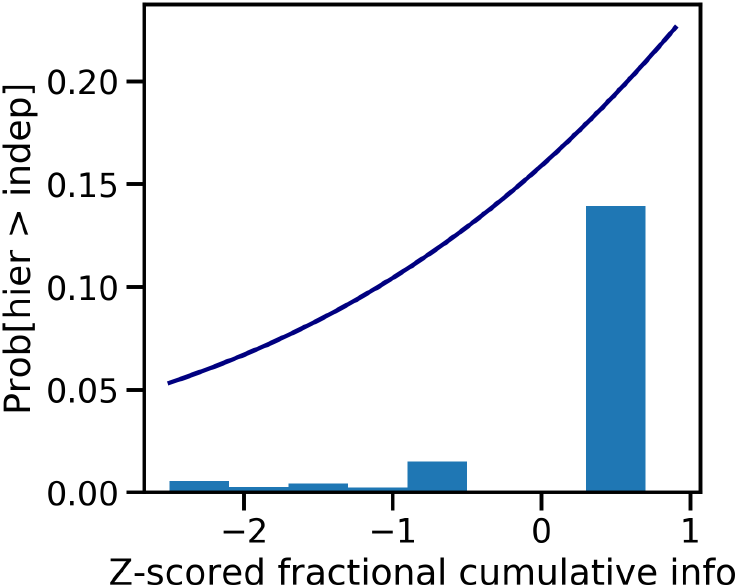
Probability of the hierarchical agent being more successful at guessing the correct door plotted against the (z-scored) fractional information content available about door sequences. Bars show the fraction of times the hierarchical agent was more successful for different bins of fractional information; the navy line shows the logistic fit. Data was aggregated across all rooms and all 50 simulations.

The Supplement provides additional histograms showing the distribution in the total number of steps each agent needed to solve each of the tasks described above.

### 3.4 Results for the non-diabolical environments

As another means of diagnosing agent transferability, we subjected them to non-diabolical versions of the hierarchical environment. Across all types of task statistics, the hierarchical and independent agents demonstrated significant positive transfer in novel contexts compared to the flat agent baseline, with the hierarchical agent being the better of the two (Fig. 14). In the independent environment, the difference between the hierarchical and independent agent was less pronounced.^4^ In the mixed environment, we also examined agent performance in different settings, including upper or sub levels only, or independent, joint, or mixed contexts only, and found the hierarchical agent to still outperform the independent agent regardless of the setting (Fig. 15 in Supplement). Finally, the Supplement provides additional violin plots showing the distribution in the number of steps each agent needed to solve these environments.

**Figure 14:**
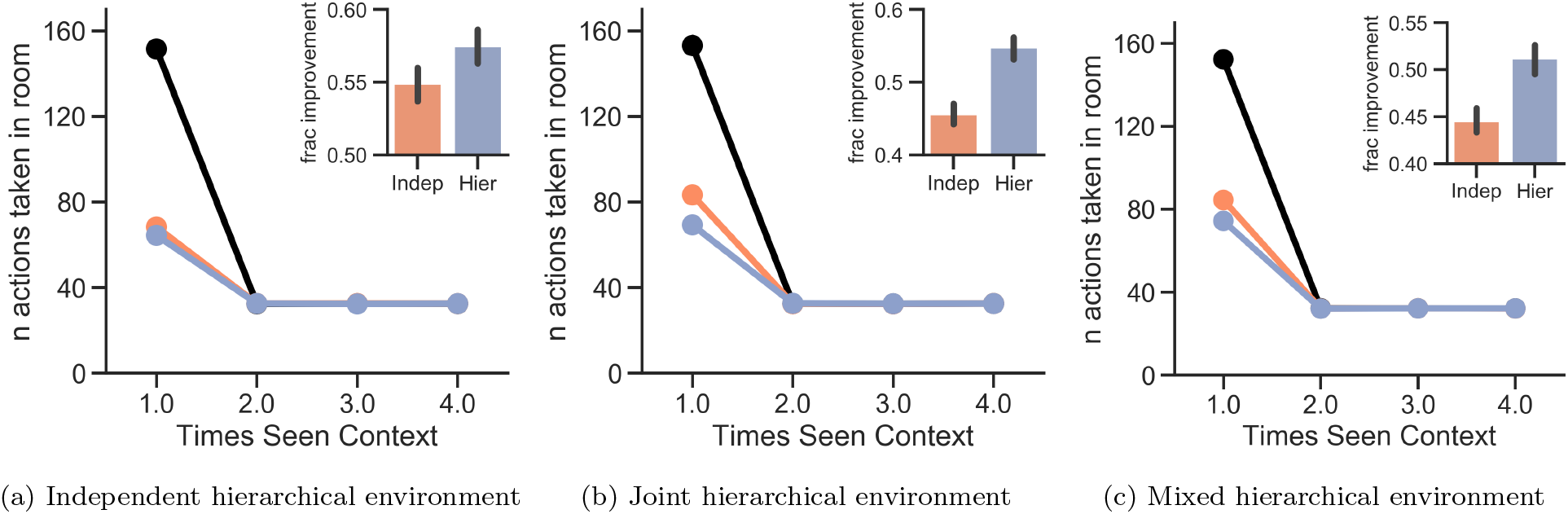
Results for the non-diabolical hierarchical task environments. In all cases, the hierarchical agent’s fractional improvement was significantly better than the independent agent’s (one-sided Welch’s t-test: *t* = 3.1, *p* = 0.0010, *df* = 1596 for the independent environment; *t* = 8.5, *p* = 1.5 × 10^−17^, *df* = 1547 for the joint environment; and *t* = 6.4, *p* = 1.5 × 10^−17^, *df* = 1518 for the mixed environment;).

## 4 Discussion

Taken together, our simulation results demonstrate the adaptive advantage of hierarchical clustering for generalization within a model-based reinforcement learning setting. Previous work has demonstrated the need to independently store and cluster task components of an MDP to support flexible compositional generalisation [14]. Yet there are cases when such a strategy can actually lead to suboptimal performance, especially when there are higher order relationships binding task components together; and in such cases, an agent that generalises jointly can actually perform better. One previous approach was to use a “meta-agent” that arbitrates between joint and independent clustering strategies [14]; but such an agent is unable to recognise clusters of joint and independent task structures existing alongside each other in the same environment. Thus it has remained an open question of how an agent can balance the need to compositionally generalise while also recognising that there are situations when transferring joint structures will actually serve the agent better. One of the principal ideas put forth by this work is that an agent must be able to do both simultaneously: it must be able to learn task components independently, but it must also be able to learn the higher order structures that combine them. Our hierarchical agent is intended to satisfy these competing demands in a principled manner. Its hierarchical structure maintains a clear delineation between the independent learning of task components, as implemented by parental DPs, and the learning of structures that combine these components together in a variety of novel ways, as implemented by room clusters. This set-up naturally allows different structures to share statistical strength by pooling knowledge about shared task components, supporting compositional generalisation. Additionally, by using structure-conditioned distributions over components, our framework also allows for a more probabilistic version of structures where component identity can be stochastic (Fig. 3). As validated by our simulations, our hierarchical agent framework leads to better and more flexible transfer across a wider range of task statistics, including those that have hierarchical goal-subgoal structure.

Indeed, theoretical formulations of compositional generalisation have often emphasised the importance of *systematicity* [16, 53], where to generalise effectively in compositional settings, one must learn not only the constituents of the composition but also the ways in which they can be combined together, what in natural languages corresponds to syntactic structures [16]. Recently, [53] has also emphasised the importance of systematicity to compositional generalisation in reinforcement learning; however, they only considered the simpler problem of compositional *interpolation* where the agent is asked to generalise to any novel combination of familiar task dimensions, which the independent clustering agent we considered above could readily handle. But as we explained above, naturalistic task domains might have additional structure to its compositions, such that certain pairings are more prevalent than others. This can be regarded as a stronger form of systematicity in compositional RL and is a closer analogue to the syntactic structures found in natural languages. As a linguistic analogy, consider the words “water”, ”kettle”, “on”, “the”, and “is”. There are 120 different permutations of these five words, but only a small subset will form meaningful propositions; for instance, the propositions “water is on the kettle”, “the kettle is on water”, and “the water kettle is on” are all meaningful, but “kettle is water on the” is not. Analogously, the stronger form of systematicity we consider here recognises that in the space of all possible combinations of task components, certain combinations will be far more prevalent than others. As such, systematicity requires learning not just the constituent components, which the independent agent is capable of doing, but also the structures that combine them. This is something that only the hierarchical agent, by its design, is capable of accomplishing. Although we have focused on compositions of rewards and transitions, our framework is quite general and can be applied to any type of compositional, hierarchical model structure. For example, the transition function itself might have further substructure where transition functions covering one region of state space might be composed with another transition function covering a different region (retaining our navigation example, one might drive across a city to a pedestrianised zone and then walk through that zone to reach a destination). Conceivably, these compositions could have independent, joint, or mixed statistics, and our framework would be able to discover and transfer such substructures in the mapping function. On a higher level of abstraction, high-dimensional MDPs, as one finds in sensory-rich naturalistic tasks, can often be compressed to discover regions of state spaces that are equivalent for the sake of planning. Discovery of such lower dimensional latent states allows an agent to transfer to novel MDPs with entirely novel reward and transition functions, as long as they retain the abstract latent structure [42]. Indeed, this work showed that one could discover useful abstractions in a guitar playing task (e.g., that all fret positions that yield a “C” note are equivalent) that allows an agent to much more rapidly learn new scales. This framework would be yet more powerful if it could compose and transfer substructures without requiring the previous abstraction to hold in all regions of the state space.

## Related work

### Compositional generalisation in other domains

Most prior work on compositional generalisation has been in the domains of natural language processing [16, 17, 18] and concept learning [19, 20, 21, 22]. In reinforcement learning, the primary focus has been on constructing new policies compositionally, the topic of hierarchical reinforcement learning [23, 24, 25, 26, 12, 27, 28, 29, 30, 31, 32]. But compositional generalisation over model structures – one important means by which we acquire models to support model-based planning – has received limited attention so far. Dayan [54] considered a factorisation of value functions into reward functions and a predictive representation of future state occupancies, known as the successor representation, which has received renewed interest in recent years [55, 56, 57, 58, 41, 59]. But this factorisation is constrained to one dimension and therefore supports limited compositionality. Moreover, the successor representation is policy-dependent and therefore cannot support transfer of model-based planning even in structurally similar environments [60, 42]. More recently, Whittington *et al*. [61] have considered a Tolman-Eichenbaum machine that factorises environments into objects and the abstract space they occupy, but this only supports the ability to associate different sets of objects with the same underlying Euclidean space, and not compositional generalization of the type we explore here.

Perhaps one of the first works to study compositional generalisation over task structures is [14]. In this work, agents learn task components, specifically reward and transition functions, independently of each other and, wherever possible, recombine them in novel ways to quickly solve new tasks. All these approaches to task decomposition, however, have the common limitation of treating task components as completely independent of each other; they are unable to learn and transfer higher order statistical structures between components whenever such structures exist. Indeed, the situation is somewhat analogous to that in natural language processing, where a distinction has been made between semantic and syntactic structures [16] – to generalise to new sentences, a language processor must be able to learn and transfer syntax as well as semantics. In task structure learning, the task components are analogous to semantics while the higher order structures are like syntaxes.

Recent work has also explored using sleep-wake algorithms to learn and infer compositional knowledge in neural networks [62]. The algorithm, known as DreamCoder, focused on learning compositions of simpler functions, building a hierarchy of increasingly complex functions to solve a increasingly challenging computational tasks. One similarity to our work was their algorithm’s ability to identify reusable computational motifs that appear across a wide variety of the solutions it generated. It packages these motifs into into reusable subroutines that are then added to its library of functions that it uses for composing together new task solutions. The value of reusing recurring compositional motifs was also demonstrated in our work, particularly with the door sequence substructure in the hierarchical environments. Yet there are several differences between DreamCoder and our approach. For one, the primitive structural components considered in DreamCoder were deterministic and well-defined mathematical operators; in our case, the agent also needed to learn the parameters of the primitive components (the mapping and reward functions). Using parametric components introduced the problem of interference in learning as different tasks could require different parametrisations for the same components, and this is something our clustering algorithm was intended to solve. Another difference is that the structures considered in DreamCoder were deterministic whereas those considered here allowed for some stochasticity in the structural components, and our higher-order clustering, in the form of room-types, allowed for the possibility of separating out distinct types of structures with distinct patterns of stochasticity in its components. Yet DreamCoder may provide a solution to one unsolved problem in our approach. The hierarchical agent works by inverting a hierarchical generative model of the tasks, yet how it acquires such models remains an open question. Perhaps the model itself can be inferred compositionally through a sleep-wake algorithm much like DreamCoder’s. Indeed, its dream-like abstraction phase provides a natural mechanism for discovering substructures, such as door sequences or sublevels, that are reused across more complex structures. We leave the possibility for such a hybrid approach to future work.

### Neural circuit implementations

Previous work using neural network models of the PFC-BG circuitry has shown that such models are capable of approximating non-parametric Bayesian inference over task-sets and other hierarchical task structures using a mixture of experts scheme and gating [43, 40], allowing the appropriate networks (and levels of hierarchy) to be deployed for a given task. The authors only considered modelling a single DP, but several of these circuits can be stacked together in a hierarchical relationship to yield an approximation of a hierarchical DP, with the higher circuit sending gating outputs to the lower circuit and which thereby influences the lower circuit’s own gating operations. Several other related neural network schemes have been advanced to model cortical recurrent networks for adaptive learning [63, 64] and even for guiding waves of dopamine across the striatum to support adaptive task learning in mice [65]. In models of category learning, neural networks have successfully learned clusters of substructures somewhat analogous to those studied here [66]. However, to date such networks have not been applied to sequential decision making in the model-based RL setting, which remains a considerable challenge for neural networks.

### Multi-task and continual learning with mixture of expert models

Successful multi-task learning hinges on the ability to transfer effectively, that is, to abstract out latent structure that does not depend on the particularities of the context and recognise when it applies again in novel contexts. To support such transfer, we have relied on a mixture-of-experts model, where each expert specialises in one of these abstract structures. Learning tunes the parameters of these experts, and transfer is cast as a problem of Bayesian inference, where the agent must infer which expert to use. Such an approach to multi-task learning has been considered previously [67, 39, 43, 40, 41, 42]. Indeed, one of the earliest examples in reinforcement learning doing this is [67]. Here, they also invert a hierarchical generative process underlying the MDPs, and then infer at the top level which of previously seen MDP classes facilitates transfer to new MDPs with familiar reward and transition functions. But this work only considered transferring task structures as a whole (i.e., the joint reward-transition function). Abstract structures are often compositional, and our work examines how individual components can also be transferred into new structures. We accomplish this by applying Bayesian mixture-of-experts across the entire model hierarchy (as opposed to just the top-level), thereby endowing our agents with the ability to generalize MDP components individually while also separately learning the structural relationships between them. Related approaches to multi-task learning have also been considered in the cognitive sciences where they have been used to model transfer of latent task structures in natural intelligence. [39] considered the use of a Dirichlet process [50, 52] to model contextual Pavlovian conditioning, where animals must infer the correct stimuli-outcome contingencies depending on the latent context. [40] extended this to instrumental conditioning, where agents must infer latent “task set” rules that govern the relationship between stimuli, actions, and outcomes in simple bandit tasks. In sequential decision-making tasks, [41] used a hierarchical generative model with a Dirichlet process to learn and transfer successor representations across tasks. [42] used Dirichlet processes to learn and transfer compressed, reward-predictive abstractions of higher-dimensional state spaces, allowing transfer even when neither reward or transition functions are familiar. Finally, [14] extended the work of [40] to allow for compositional generalisation of task models. Our work directly builds on [14] by introducing a unified framework that allows an agent to learn and transfer individual components at the same time as higher order structures. We do this by introducing a hierarchical Dirichlet process to model the task’s generative process. This framework gives the agent the flexibility needed to adapt to a much wider range of task statistics and, as we have shown, can even scale to more complex, hierarchical tasks with goal/subgoal structures.

These mixture-of-expert approaches to multi-task learning considered the setting whether the number of task structures (and hence the number of experts needed) cannot be known in advance and may even be infinite. Like us, these approaches used Dirichlet processes to incrementally expand the number of experts as more tasks are seen. Non-parametric methods have also been used to model complex, multi-modal data distributions and to support transfer in a few-shot setting [68, 69, 70, 71]. The main advantage of these approaches is that they adapt model capacity to the data distribution, thereby avoiding both underfitting due to insufficient capacity as well as overfitting due to an excess of capacity.

Mixture-of-expert approaches are also better adapted to situations where the set of tasks is sufficiently heterogeneous that some tasks will be significantly different from each other. In recent years, gradient-based meta-learning has proven to be one of the most powerful approaches to multi-task learning, demonstrating state-of-the-art performance when adapting few-shot to novel tasks [72, 64, 73, 74, 75]. These approaches focus on learning a single set of features to transfer across all tasks. But this can be suboptimal if the set of tasks is sufficiently heterogeneous that there are tasks significantly dissimilar to each other, as the agent will struggle to find a common set of features that is adequately suited to the task diversity [71]. Moreover, when transferring to novel tasks not seen during training, these latent features will only support positive transfer if the novel tasks also share these features; they will lead instead to negative interference if the test tasks are sufficiently different to the training tasks. Previous work has leveraged Dirichlet processes to address this problem [69, 70, 71]. These works have considered the multi-task setting where the set of tasks is heterogeneous and better characterised by smaller clusters of related tasks. Much like in our work, they have used DPs to group tasks together into distinct clusters. Tasks that are sufficiently similar to each other and that share similar latent features get assigned to the same cluster while tasks that are sufficiently different get assigned to different clusters, much like our room-type clusters. And as in our work, this approach supports feature sharing and positive transfer amongst tasks within a cluster while preventing interference between unrelated tasks. However, these approaches take an all-or-nothing approach to feature sharing and do not allow for the possibility of sharing in a compositional manner. Generally, two sets of tasks can be related along some dimensions but highly unrelated along others. These approaches will generally group such tasks into separate clusters, preventing feature sharing along the related dimensions. In contrast, our compositional approach allows different clusters to still share features along related dimensions while simultaneously separating out features along unrelated dimensions. This approach furthermore allows learning of common features to be shared across different clusters.

Bayesian non-parametrics also offers one possible solution to two of the challenges inherent in continual learning. Continual learning requires the agent to continually adapt to an ever-evolving task distribution, introducing additional challenges not present in the static, multi-task setting. Specifically, in the static setting, the agent learns a fixed set of features that can be transferred across tasks at test time. The features will reflect their relative prevalences in the training task set and allow the agent to adapt rapidly to new tasks when sampled from the same task distribution. Indeed, one can understand static meta-learning as learning not only the shared features themselves but also an implicit prior distribution over the features [75, 76]. So should the distribution change significantly, as may be the case in continual learning, this bias can lead to a form of negative transfer as the most likely set of relevant features under the new task distribution might be under-weighted by the prior. As we have seen, Bayesian non-parametrics offers one possible solution to this problem by allowing for priors over shared task features that can continually adapt and update as more tasks are encountered. Continual learning also requires that learning of novel tasks not interfere with knowledge of familiar ones, and Bayesian non-parametrics offers a solution by assigning different tasks to different task clusters. Essentially, Bayesian non-parametrics can address the challenges of continual learning by using a mixture of experts approach: distinct tasks get assigned to different experts, which helps avoid catastrophic interference, but each expert can also pool knowledge about common features from the tasks it handles and thereby encourage transfer across tasks. Indeed, DP-based approaches have been used to address these issues [71], and our work continues in a similar vein.

### Task design

We have tested our agents on tasks adapted from those in [14]. To our knowledge, only these tasks have been explicitly designed to explore compositional generalisation of task structures. These tasks are themselves a modification of the grid-sailing tasks of Fermin *et al*. [77], which introduced an explicitly compositional MDP, but the grid-sailing tasks were not designed to study generalisation to new structures. Our work uses the original tasks from [14] as well as modified versions designed to study more graded statistics in task structures. We have also combined the grid-sailing tasks with the classic “rooms” problem of hierarchical reinforcement learning [24, 12]) to demonstrate the benefits to more hierarchically-structured compositional environments.

### Limitations and future directions

Compositional generalisation has generally been regarded as an essential feature of natural intelligence. Beyond the navigational examples we have referenced throughout this paper, compositional structure shows up in a myriad of different naturalistic task domains. In musical domains, fingerings can be transferred across different, sufficiently-related instruments while songs can be transferred across nearly any instrument. Social behaviour can also be compositional in structure: when conducting meetings in foreign lands, one might recognise that one should greet one’s hosts first, but how one *how* one goes about doing that may vary according to local norms and can involve shaking hands, bowing, or in the context of a global pandemic, bumping elbows. And perhaps the best example of compositionality are languages, both natural and artificial (such as mathematics in the latter case). Compositional structure therefore shows up regularly in naturalistic tasks, and it is thought that natural intelligence readily leverages this to rapidly solve new tasks. Indeed, previous work [15] has shown that humans do reasonably well at some of the tasks that we have examined above and in a manner broadly consistent with the hierarchical model. However, more work is still needed to test the specific predictions of our model in human learning. This would validate whether natural intelligence learns compositional structures in a manner consistent with the hierarchical agent.

One limitation of our approach is that the hierarchical agent does have an initial disadvantage in terms of sample complexity. Credit assignment becomes harder as the model becomes more with greater levels of hierarchy (see also [40] for an example of how hierarchical structure in neural networks give rise to the same credit assignment issue, hindering early learning but facilitating transfer). If the agent does not get enough training data on old contexts before moving on to new ones, it is more liable to throw out correct clustering hypotheses during the particle filtering step, and this manifests as the occasional outlier where the agent takes an inordinately long time to solve the environment in a simulation (see violin plots in the Supplement). Nevertheless, the greater flexibility in transfer that the hierarchical model affords outweighs the cost of the increased complexity, especially when there is a heterogeneity of structures present in the environment, as is the case in the real world.

Yet, there are other limitations that future work will also need to address. Currently, our approach relies on signalled context changes (e.g., that the agent has entered a new room), whereas such changes can be more latent in naturalistic tasks and must therefore be inferred from observations. A natural way to address this issue would be to introduce hidden Markov model-like inference (or resource-constrained approximations thereof) to determine context boundaries. Another limitation is that in naturalistic tasks, familiar contexts can change over time, making the initial cluster assignment no longer appropriate after a certain point. In such cases, it may make more sense to reassign the context to a different cluster rather than have cluster assignments be static and irreversible.

Given the model-based setting of our work, as in other such schemes, agents were endowed with knowledge that there are task components such as transition functions and reward functions that can be used for planning. In this sense, the hierarchical structure given to our agents can be regarded as part of the task model that a model-based agent would have access to. Similarly, in the hierarchical task with sublevels, the agents also knows of the task’s hierarchical structure (e.g., that doors in upper levels may lead to sublevels, which in turn have exits). Yet, in both hierarchical and non-hierarchical settings, the agents do not know the parameters of such transition and reward functions, or how they combine them, and have to infer them to plan correctly. Indeed, as we mentioned above, the agents that operate on hierarchical tasks can be regarded as a generalisation of the agents that operate in the non-hierarchical tasks. If these more general agents were to solve the non-hierarchical tasks, they would simply not recruit the parts of their hierarchy used to represent the sublevels or the unused doors in the door sequences. Analogously, human cognitive neuroscience studies have shown that humans can flexibly shift between hierarchical and non-hierarchical tasks by recruiting increasingly anterior regions of the prefrontal cortex-basal ganglia (PFC-BG) circuitry and that they can learn to do so via reinforcement learning [44].

Ultimately, we expect any intelligent agent, whether natural or artificial, to implement an approximation, as constrained by computational resources, to the algorithm outlined in our framework, and it remains an open question of how our hierarchical model can be implemented in neural circuits. In artificial agents, previous work has used hybrid architectures wherein deep networks implement likelihood functions and network parameters are clustered together through a DP [41, 78, 71], essentially yielding a non-parametric mixture of experts model. Such an approach can certainly be adapted to our framework; in our case, the likelihood functions 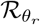and 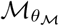 would simply be implemented by deep networks as well, and our hierarchical DP clusters would simply encode network parameters *θ*_*r*_ and *θ*_ℳ_. A neural network implementation of the DP is still lacking, however, although hypernetworks [79] may offer one solution, as they may, in principle, learn to modify the functions implemented by the downstream networks in a context-dependent manner. Alternatively, one could also look to neurobiology for some clues. As discussed earlier, hierarchical neural network models of the PFC-BG circuitry have shown that it is capable of approximating non-parametric Bayesian inference over task structures [43, 40], and we have speculated that this circuitry may even approximate the sort of hierarchical clustering we have considered here. Yet how such models can be scaled to more complex tasks, or even translated into gradient-based deep network models, remains something that future work will need to explore.

## 5 Conclusion

Our framework offers a normative perspective on what is necessary computationally for an agent to learn hierarchical, compositional task structures – in terms of Marr’s three levels of analyses [80], the framework can be understood as addressing both the computational and algorithmic levels. And through our empirical results, we have validated our approach, showing that only an agent that learned structures hierarchically was capable of supporting compositional generalisation as well as effective transfer of structural knowledge to a wider range of tasks.

## Code availability

All simulation code can be found at https://github.com/RexGLiu/HierPolicies.

## Acknowledgements

The authors would like to thank Nicholas Franklin, Cris Buc Calderon and Matt Nassar for very helpful comments in improving the quality of the manuscript. We also gratefully acknowledge the use of Nicholas Franklin’s code [14] (https://github.com/nicktfranklin/IndependentClusters), upon which we have built our work. This work was conducted using computational resources and services at the Center for Computation and Visualization, Brown University, which is supported by NIH grant S10OD025181. RGL is supported by funding from the Carney Institute for Brain Sciences. MJF is supported by NIH grants MH084840-08A1 and MH119467-01.

## Footnotes

1 We note that the parent CRPs also express a different type of prior compared to their children. The children CRPs are much like their counterparts in the non-hierarchical agents – in the absence of any data, clusters that were more popular in a given room type will be favoured for other novel contexts associated with that room type. A parent CRP, instead, has a bias towards diversity. It will prefer clusters that appear across a wider range of children CRPs over ones that might be far more frequent overall but special to a smaller number of children CRPs. This captures the intuition that a task component for a novel structure is more likely to be one that is generic to all structures rather than idiosyncratic to a few.

2 To see that the goals and mappings are statistically independent, note that given a randomly chosen context, the probability that its goal is, for instance, the blue square is 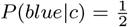 regardless of context *c*, and this probability remains the same if the context’s mapping is also given; that is, 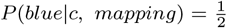 regardless of *c* or mapping. Analogous arguments follow both for the other goal as well as for showing that *P* (*mapping*|*c*) = *P* (*mapping*|*c, goal*). These equalities do not hold for the joint environment that we consider later.

3 Unlike previously, we did not include a fully joint agent as jointness can be defined on multiple levels, such as across the entire room, at the level of individual upper and sublevels, at the level of sublevel sequences within a room, and at the level of exit sequences amongst a room’s sublevels, to name just a few possibilities. The task structure involves many subcomponents which affords many possibilities of jointness, and selecting one or a subset of agents that assume different types of jointness would involve a certain degree of arbitrariness. For this reason, we have chosen not to include a joint agent baseline for these tasks. Indeed, as we mentioned earlier, the combinatorial explosion in the number of different possible structures due to the increased complexity of task structure implies that many structures are never seen twice in exactly the same way in an agent’s lifetime. In such cases, memorising all possible structures an agent sees throughout the course of its lifetime would be a highly suboptimal strategy to follow.

4 This may be because the diabolical nature of the previous environments amplified any performance advantage one agent has over the other, because over repeated resets, any small performance difference in individual gridworlds will eventually accumulate into a large final difference by the environment’s end.

## References

[1] Collins AGE, Frank MJ. Neural signature of hierarchically structured expectations predicts clustering and transfer of rule sets in reinforcement learning. Cognition. 2016;152:160–169.

[2] Baram AB, Muller TH, Nili H, Gavert MM, Behrens TEJ. Entorhinal and ventromedial prefrontal cortices abstract and generalize the structure of reinforcement learning problems. Neuron. 2020;109:1–11.

[3] Harlow H. The formation of learning sets. Psychological Review. 1949;56:51–65.

[4] Mnih V, et al. Human-level control through deep reinforcement learning. Nature. 2015;518:529–533.

[5] Badia AP, et al. Agent57: Outperforming the Atari Human Benchmark. 200313350. 2020.

[6] Silver D, et al. Mastering the game of Go with deep neural networks and tree search. Nature. 2016;529:484–489.

[7] Silver D, et al. Mastering the game of Go without human knowledge. Nature. 2017;550:354–359.

[8] Taylor ME, Stone P. Transfer Learning for Reinforcement Learning Domains: A Survey. JMLR. 2009;10:1633–1685.

[9] Kansky K, et al. Schema Networks: Zero-shot Transfer with a Generative Causal Model of Intuitive Physics. 170604317. 2017.

[10] Schultz W, Dayan P, Montague R. A Neural Substrate of Prediction and Reward. Science. 1997;275:1593–1599.

[11] Collins AGE, Frank MJ. Opponent actor learning (OpAL): Modeling interactive effects of striatal dopamine on reinforcement learning and choice incentive. Psychological Review. 2014;121(3):337–366. Available from: http://doi.apa.org/getdoi.cfm?doi=10.1037/a0037015.

[12] Botvinick MM, Niv Y, Barto AC. Hierarchically organized behavior and its neural foundations: A reinforcement learning perspective. Cognition. 2009;113:262–280.

[13] Sutton RS, Barto AG. Reinforcement Learning: An Introduction. Cambridge, MA: MIT Press; 2018.

[14] Franklin NT, Frank MJ. Compositional clustering in task structure learning. PLoS Comput Biol. 2018;14:e1006116.

[15] Franklin NT, Frank MJ. Generalizing to generalize: Humans flexibly switch between compositional and conjunctive structures during reinforcement learning. PLoS Comput Biol. 2020;16:e1007720.

[16] Chomsky AN. Syntactic Structures. Berlin, Germany: Mouton & Co.; 1957.

[17] Marcus GF. Rethinking Eliminative Connectionism. Cognitive Psychology. 1998;37:243–282.

[18] Lake B, Baroni M. Generalization without Systematicity: On the Compositional Skills of Sequence-to-Sequence Recurrent Networks. In: Dy J, Krause A, editors. Proceedings of the 35th International Conference on Machine Learning. PMLR; 2018. p. 2873–2882. Available from: https://proceedings.mlr.press/v80/lake18a.html.

[19] Fodor JA, Pylyshyn ZW. Connectionism and Cognitive Architecture: A Critical Analysis. Cognition. 1988;28:3–71.

[20] Marcus GF. The Algebraic Mind: Integrating Connectionism and Cognitive Science. Cambridge, USA: MIT Press; 2003.

[21] Lake BM, Salakhutdinov R, Tenenbaum JB. Human-level concept learning through probabilistic program induction. Science. 2015;350:1332–1338.

[22] Frankland SM, Greene JD. Concepts and Compositionality: In Search of the Brain’s Language of Thought. Ann Rev Psychol. 2020;71:273–303.

[23] Dayan P, Hinton GE. Feudal reinforcement learning. In: Hanson S, Cowan J, Giles C, editors. Advances in Neural Information Processing Systems. vol. 5. Morgan-Kaufmann; 1993. Available from: https://proceedings.neurips.cc/paper/1992/file/d14220ee66aeec73c49038385428ec4c-Paper.pdf.

[24] Sutton RS, Precup D, Singh S. Between MDPs and semi-MDPs: A framework for temporal abstraction in reinforcement learning. Artificial Intelligence. 1999;112:181–211.

[25] Parr R, Russell S. Reinforcement Learning with Hierarchies of Machines. In: Jordan M, Kearns M, Solla S, editors. Advances in Neural Information Processing Systems. vol. 10. MIT Press; 1998. Available from: https://proceedings.neurips.cc/paper/1997/file/5ca3e9b122f61f8f06494c97b1afccf3-Paper.pdf.

[26] Diettrich TG. Hierarchical reinforcement learning with the MAXQ value function decomposition. JAIR. 2000;13:227–303.

[27] Vezhnevets AS, Osindero S, Schaul T, Heess N, Jaderberg M, Silver D, et al. FeUdal Networks for Hierarchical Reinforcement Learning. In: Precup D, Teh YW, editors. Proceedings of the 34th International Conference on Machine Learning. PMLR; 2017. p. 3540–3549. Available from: https://proceedings.mlr.press/v70/vezhnevets17a.html.

[28] Kulkarni TD, Narasimhan KR, Saeedi A, Tenenbaum JB. Hierarchical Deep Reinforcement Learning: Integrating Temporal Abstraction and Intrinsic Motivation. In: Lee D, Sugiyama M, Luxburg U, Guyon I, Garnett R, editors. Advances in Neural Information Processing Systems. vol. 29. Curran Associates, Inc.; 2016. Available from: https://proceedings.neurips.cc/paper/2016/file/f442d33fa06832082290ad8544a8da27-Paper.pdf.

[29] Silver D, Ciosek K. Compositional Planning Using Optimal Option Models. In: Langford J, Pineau J, editors. Proceedings of the 29th International Conference on Machine Learning. PMLR; 2012. p. 1267–1274. Available from: https://arxiv.org/abs/1206.6473.

[30] Wingate D, Diuk C, O’Donnell TJ, Tenenbaum JB, Gershman SJ. Compositional Policy Priors; 2013. Available from: https://dspace.mit.edu/handle/1721.1/78573.

[31] Eschenbach B, Gupta A, Ibarz J, Levine S. Diversity is All You Need: Learning Skills without a Reward Function. In: International Conference on Learning Representations; 2019. Available from: https://openreview.net/forum?id=SJx63jRqFm.

[32] Tirumala D, Galashov A, Noh H, Hasenclever L, Pascanu R, Schwarz J, et al. Behavior Priors for Efficient Reinforcement Learning; 2020. Available from: https://arxiv.org/abs/2010.14274.

[33] Hessel M, Modayil J, v Hasselt H, Schaul T, Ostrovski G, Dabney W, et al. Rainbow: Combining Improvements in Deep Reinforcement Learning. In: The Thirty-Second AAAI Conference on Artificial Intelligence (AAAI-18); 2018. Available from: https://aaai.org/ocs/index.php/AAAI/AAAI18/paper/view/17204.

[34] Mnih V, Badia AP, Mirza M, Graves A, Harley T, Lillicrap TP, et al. Asynchronous Methods for Deep Reinforcement Learning. In: Balcan MF, Weinberger KQ, editors. Proceedings of the 33rd International Conference on Machine Learning. PMLR; 2016. p. 1928–1937. Available from: http://proceedings.mlr.press/v48/mniha16.pdf.

[35] Schulman J, Levine S, Moritz P, Jordan M, Abbeel P. Trust Region Policy Optimization. In: Bach F, Blei D, editors. Proceedings of the 32nd International Conference on Machine Learning. PMLR; 2015. p. 1889–1897. Available from: http://proceedings.mlr.press/v37/schulman15.pdf.

[36] Schulman J, Wolski F, Dhariwal P, Radford A, Klimov O. Proximal Policy Optimization Algorithms; 2017. Available from: https://arxiv.org/abs/1707.06347.

[37] Haarnoja T, Zhou A, Abbeel P, Levine S. Soft Actor-Critic: Off-Policy Maximum Entropy Deep Reinforcement Learning with a Stochastic Actor. In: Dy J, Krause A, editors. Proceedings of the 35th International Conference on Machine Learning. PMLR; 2018. p. 1861–1870. Available from: http://proceedings.mlr.press/v80/haarnoja18b/haarnoja18b.pdf.

[38] Lillicrap T, Hunt JJ, Pritzel A, Heess N, Erez T, Tassa Y, et al. Continuous control with deep reinforcement learning. In: International Conference on Learning Representations; 2016. Available from: https://openreview.net/forum?id=tX_O8O-8Zl.

[39] Gershman SJ, Blei DM, Niv Y. Context, learning, and extinction. Psychological Review. 2010;117:197–209.

[40] Collins AGE, Frank MJ. Cognitive Control Over Learning: Creating, Clustering, and Generalizing Task-Set Structure. Psychological Review. 2013;120:190–229.

[41] Madarász TJ, Behrens TEJ. Better Transfer Learning with Inferred Successor Maps. In: Wallach H, Larochelle H, Beygelzimer A, d’Alché-Buc F, Fox E, Garnett R, editors. Advances in Neural Information Processing Systems. vol. 32. Curran Associates, Inc.; 2019. Available from: https://proceedings.neurips.cc/paper/2019/file/274a10ffa06e434f2a94df765cac6bf4-Paper.pdf.

[42] Lehnert L, Littman ML, Frank MJ. Reward-predictive representations generalize across tasks in reinforcement learning. PLoS Comput Biol. 2020;16:e1008317.

[43] Frank MJ, Badre D. Mechanisms of hierarchical reinforcement learning in corticostriatal circuits 1: Computational analysis. Cereb Cortex. 2012;22:509–526.

[44] Badre D, Frank MJ. Mechanisms of hierarchical reinforcement learning in corticostriatal circuits 2: Evidence from fMRI. Cereb Cortex. 2012;22:527–536.

[45] Collins AGE, Cavanagh JF, Frank MJ. Human EEG Uncovers Latent Generalizable Rule Structure during Learning. J Neurosci. 2014;34:4677–4685.

[46] Tomov MS, Dorfman HF, Gershman SJ. Neural Computations Underlying Causal Structure Learning. J Neurosci. 2018;38:7143–7157.

[47] Luyckx F, Nili H, Spitzer B, Summerfield C. Neural structure mapping in human probabilistic reward learning. eLife. 2019;8:e42816.

[48] Teh YW, Jordan MI, Beal MJ, Blei DM. Hierarchical Dirichlet Processes. Journal of the American Statistical Association. 2006;101:1566–1581.

[49] Hallak A, Di Castro D, Mannor S. Contextual Markov Decision Processes. 161102779. 2015.

[50] Ferguson TS. A Bayesian analysis of some nonparametric problems. Ann Statist. 1973;1:209–230.

[51] Aldous DJ. Exchangeability and related topics. In: Hennequin PL, editor. École d’Été de Probabilités de Saint-Flour XIII – 1983. Berlin, Heidelberg: Springer Berlin Heidelberg; 1985. p. 1–198.

[52] Teh YW. Dirichlet Process. In: Sammut C, Webb GI, editors. Encyclopedia of Machine Learning. Boston, MA: Springer US; 2011. p. 280–287.

[53] Kirk R, Zhang A, Grefenstette E, Rocktäschel T. A survey of generalisation in deep reinforcement learning. 211109794. 2021.

[54] Dayan P. Improving Generalization for Temporal Difference Learning: The Successor Representation. Neural Computation. 1993;5:613–624.

[55] Kulkarni TD, Saeedi A, Gautam S, Gershman SJ. Deep Successor Reinforcement Learning; 2016. Available from: https://arxiv.org/abs/1606.02396.

[56] Barreto A, Dabney W, Munos R, Hunt JJ, Schaul T, v Hasselt HP, et al. Successor Features for Transfer in Reinforcement Learning. In: Guyon I, Luxburg UV, Bengio S, Wallach H, Fergus R, Vishwanathan S, et al., editors. Advances in Neural Information Processing Systems. vol. 30. Curran Associates, Inc.; 2017. Available from: https://papers.nips.cc/paper/2017/file/350db081a661525235354dd3e19b8c05-Paper.pdf.

[57] Russek EM, Momennejad I, Botvinick MM, Gershman SJ, Daw ND. Predictive representations can link model-based reinforcement learning to model-free mechanisms. PLoS Comput Biol. 2017;13:e1005768.

[58] Barreto A, Borsa D, Quan J, Schaul T, Silver D, Hessel M, et al. Transfer in Deep Reinforcement Learning Using Successor Features and Generalised Policy Improvement. In: Dy J, Krause A, editors. Proceedings of the 35th International Conference on Machine Learning. vol. 80 of Proceedings of Machine Learning Research. PMLR; 2018. p. 501–510. Available from: https://proceedings.mlr.press/v80/barreto18a.html.

[59] Vértes E, Sahani M. A neurally plausible model learns successor representations in partially observable environments. In: Wallach H, Larochelle H, Beygelzimer A, d’Alché-Buc F, Fox E, Garnett R, editors. Advances in Neural Information Processing Systems. vol. 32. Curran Associates, Inc.; 2019. Available from: https://proceedings.neurips.cc/paper/2019/file/dea184826614d3f4c608731389ed0c74-Paper.pdf.

[60] Lehnert L, Littman ML. Successor Features Combine Elements of Model-Free and Model-based Reinforcement Learning. JMLR. 2020;21:1–53.

[61] Whittington JCR, Muller TH, Mark S, Chen G, Barry C, Burgess N, et al. The Tolman-Eichenbaum Machine: Unifying Space and Relational Memory through Generalization in the Hippocampal Formation. Cell. 2020;183:1249–1263.e23.

[62] Ellis K, et al. DreamCoder: Growing generalizable, interpretable knowledge with wake-sleep Bayesian program learning. 200608381. 2020.

[63] Tsuda B, Tye KM, Siegelmann HT, Sejnowski TJ. A modeling framework for adaptive lifelong learning with transfer and savings through gating in the prefrontal cortex. PNAS. 2020;117:29872–29882.

[64] Wang JX, Kurth-Nelson Z, Kumaran D, Tirumala D, Soyer H, Leibo JZ, et al. Prefrontal cortex as a meta-reinforcement learning system. Nat Neurosci. 2018;21:860–868.

[65] Hamid AA, Frank MJ, Moore CI. Wave-like dopamine dynamics as a mechanism for spatiotemporal credit assignment. Cell. 2021;184:2733–2749.e16.

[66] Love BC, Medin DL, Gureckis TM. SUSTAIN: A Network Model of Category Learning. Psychological Review. 2004;111:309–332.

[67] Wilson A, Fern A, Ray S, Tadepalli P. Multi-task reinforcement learning: A hierarchical bayesian approach. In: Ghahramani Z, editor. Proceedings of the 24th International Conference on Machine Learning. Omni Press; 2007. p. 1015–1022. Available from: https://icml.cc/imls/conferences/2007/proceedings/papers/463.pdf.

[68] Allen K, Shelhamer E, Shin H, Tenenbaum J. Infinite Mixture Prototypes for Few-shot Learning. In: Chaudhuri K, Salakhutdinov R, editors. Proceedings of the 36th International Conference on Machine Learning. vol. 97 of Proceedings of Machine Learning Research. PMLR; 2019. p. 232–241. Available from: https://proceedings.mlr.press/v97/allen19b.html.

[69] Xue Y, Liao X, Carin L, Krishnapuram B. Multi-Task Learning for Classification with Dirichlet Process Priors. JMLR. 2007;10:35–63.

[70] Gupta S, Phung D, Venkatesh S. Factorial Multi-Task Learning: A Bayesian Nonparametric Approach. In: Dasgupta S, McAllester D, editors. Proceedings of the 30th International Conference on Machine Learning. vol. 28 of Proceedings of Machine Learning Research. PMLR; 2013. p. 657–665. Available from: https://proceedings.mlr.press/v28/gupta13a.html.

[71] Jerfel G, Grant E, Griffiths T, Heller K. Reconciling meta-learning and continual learning with online mixtures of tasks. In: Wallach H, Larochelle H, Beygelzimer A, d’Alché-Buc F, Fox E, Garnett R, editors. Advances in Neural Information Processing Systems. vol. 32. Curran Associates, Inc.; 2019. Available from: https://proceedings.neurips.cc/paper/2019/file/7a9a322cbe0d06a98667fdc5160dc6f8-Paper.pdf.

[72] Wang JX, Kurth-Nelson Z, Tirumala D, Soyer H, Leibo JZ, Munos R, et al. Learning to reinforcement learn. 161105763. 2016.

[73] Duan Y, Schulman J, Chen X, Bartlett PL, Sutskever I, Abbeel P. RL$^2$: Fast Reinforcement Learning via Slow Reinforcement Learning. 161102779. 2016.

[74] Finn C, Abbeel P, Levine S. Model-Agnostic Meta-Learning for Fast Adaptation of Deep Networks. In: Precup D, Teh YW, editors. Proceedings of the 34th International Conference on Machine Learning. vol. 70 of Proceedings of Machine Learning Research. PMLR; 2017. p. 1126–1135. Available from: http://proceedings.mlr.press/v70/finn17a.html.

[75] Finn C, Xu K, Levine S. Probabilistic Model-Agnostic Meta-Learning. In: Bengio S, Wallach H, Larochelle H, Grauman K, Cesa-Bianchi N, Garnett R, editors. Advances in Neural Information Processing Systems. vol. 31. Curran Associates, Inc.; 2018. Available from: https://papers.nips.cc/paper/2018/file/8e2c381d4dd04f1c55093f22c59c3a08-Paper.pdf.

[76] Ortega PA, et al. Meta-learning of Sequential Strategies. DeepMind Tech Report. 2019;1.1:1–15.

[77] Fermin A, Yoshida T, Ito M, Yoshimoto J, Doya K. Evidence for Model-Based Action Planning in a Sequential Finger Movement Task. J Mot Behav. 2010;42:371–379.

[78] Nagabandi A, Finn C, Levine S. Deep Online Learning Via Meta-Learning: Continual Adaptation for Model-Based RL. In: International Conference on Learning Representations; 2019. Available from: https://openreview.net/forum?id=HyxAfnA5tm.

[79] Ha D, Dai AM, L. QV. HyperNetworks. In: International Conference on Learning Representations; 2017. Available from: https://openreview.net/forum?id=rkpACe1lx.

[80] Marr DC. Vision: A Computational Investigation into the Human Representation and Processing of Visual Information. San Francisco: W. H. Freeman & Co.; 1982.

