## Supplement for "Hierarchical clustering optimizes the tradeoff between compositionality and expressivity of task structures for flexible reinforcement learning"

### 1 Agent implementation details

Actions were selected for each agent as follows. Because the reward and mapping functions in any given context are unknown, we use those given by the MAP clustering hypothesis for evaluating the value of the cardinal actions and for the policy. We first applied $\mathcal{R}_{MAP}$ to solve the Bellman optimality equation for cardinal actions,

$$Q_c^*(s, a_{card}) = \sum_{s'} p(s'|s, a_{card}) \left[ \mathcal{R}_{MAP}^{k_{\mathcal{R}}}(s') + \max_{a'_{card}} \gamma \, Q_c^*(s', a'_{card}) \right],$$

where $c$ is the current context and $k_{\mathcal{R}}$ indexes the reward cluster to it has been assigned. Solutions were found through policy iteration. We next constructed a policy for the cardinal actions by taking a soft-max over $Q_c$,

$$\pi_{card}^c(a_{card}|s) = \frac{e^{\beta \, Q_c(s, \, a_{card})}}{\sum_{a_{card} \in \mathcal{A}_{card}} e^{\beta \, Q_c(s, \, a_{card})}},$$

where $\beta$ is an inverse temperature parameter. Similarly, we took the MAP hypothesis' mapping function $\mathcal{M}_{MAP}^{k_{\mathcal{M}}}$ to convert this into a policy in terms of literal actions ("key presses"),

$$\pi^c(a|s) = \sum_{a_{card} \in \mathcal{A}_{card}} \mathcal{M}_{MAP}^{k_{\mathcal{M}}}(a|a_{card}) \, \pi_{card}^c(a_{card}|s),$$

where $k_{\mathcal{M}}$ is the cluster to which context $c$ has been assigned. Keys are then sampled randomly from this policy.

We encouraged initial optimistic exploration by initialising all reward probabilities to $\mathcal{R}^c(r|s) = 1$, as is common in RL (e.g. [1]). Mapping probabilities were initialized to $P(a_{card}|a, c) = 1/n_{actions}$, where $n_{actions} = 8$ is the number of keys. As the agent interacted with the environment, probabilities were updated to reflect the maximum likelihood estimate given the data assigned to the relevant cluster.

As the agent sees more contexts or rooms, the hypothesis space of possible clusterings expands exponentially, making it impossible to both store or perform computations over the full posterior distribution. For this reason, the DPs are instantiated as a set of particles sampled from the hypothesis space. The particles are then filtered every time the hypothesis space needs to be expanded. Specifically, in the non-hierarchical environments, each time we add a new context to the clusterings, we first discard all but the 300 *maximum a posteriori* hypotheses; we then expand out the remaining hypotheses in all possible ways that the new context can be added to these clusterings but retained only the top 10000 hypotheses immediately afterwards. It has been shown that keeping the *maximum a posteriori* (MAP) hypotheses will minimize the Kullback-Leibler divergence between the particle representation and the true posterior [2]. In the hierarchical environments, we instead retained only the best 100 hypotheses. However, to speed up computation, we also discarded all but the top 20000 *maximum a posteriori* hypotheses immediately after expanding the hypotheses.

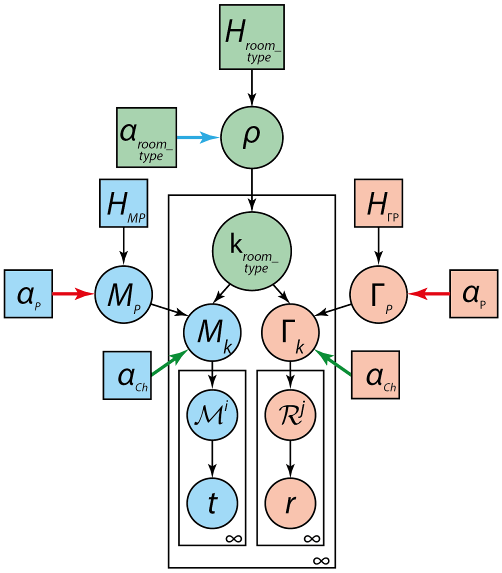


Figure 1: By manipulating the concentration hyperparameters $\{\alpha\}$ (associated with coloured arrows), the hierarchical agent can be reduced to behave like either the independent or joint agent.

Finally, it is important to note that at the level of reward and mapping clusters, all agents operate in exactly the same way. Ultimately, the only difference between agents lies in the structure of the hypothesis space of context clusterings; this determines the possible Bayesian priors used to support transfer and is, alone, responsible for any differences in performance between the agents.

#### 2 Hierarchical agent subsumes the independent and joint agents

The hierarchical agent subsumes the hypothesis spaces of both the independent and joint clustering agents, making the hierarchical agent more general than either of the two. Indeed, by an appropriate choice of hyperparameters, we can actually force the hierarchical agent to behave exactly like either of the two agents, showing that they are special cases. We can force the hierarchical agent to mimic the independent agent by setting $\alpha_{room\_type}$ to zero (Fig. 1, blue arrow) and setting $\alpha_{Ch}$ (Fig. 1, green arrows) to $\alpha$ in the independent agent. Setting $\alpha_{room\_type}$ to zero forces the agent to assign all contexts to only one room cluster, in which the reward and mapping functions across all gridworlds would get clustered independently of each other. Conversely, we can force the hierarchical agent to mimic the joint agent by setting $\alpha_{Ch}$ to zero (Fig. 1, green arrows), so that only one mapping and reward cluster gets created inside a room cluster. Each room cluster would then act like a joint cluster, and $\alpha_{room\_type}$ would act like $\alpha$ in the joint clustering agent. For both agents, we would also need $\alpha_P$ to be infinite (Fig. 1, red arrows) so that the children CRPs always get a new cluster from the parent CRPs, rather than reusing a cluster already existing in the parent. We have summarised the hyperparameters for these two special cases in Table 1.

| | Independent | Joint |
|---|---|---|
| $\alpha_{room\_type}$ | 0 | arbitrary |
| $\alpha_{Ch}$ | arbitrary | 0 |
| $\alpha_P$ | $\infty$ | $\infty$ |

Table 1: Hyperparameters for forcing the hierarchical agent into either the independent or joint clustering agents.

#### 3 Details of the non-hierarchical environments

For these environments, goals were located at $\{(0,0),(0,5),(5,0),(5,5)\}$. In each gridworld, agents always started in a random location drawn randomly from $\{(x,y)\,|\,x \in [1,5] \text{ and } y \in [1,5]\}$. All environments were repeated for a total of 150 simulations.

The tables that follow indicate the number of distinct contexts for each pair of mapping and goal. Mappings were generated without overlap; that is, no keys will map to the same cardinal action if the mappings are different.

| | Goal $(0,0)$ | Goal $(5,5)$ |
|---|---|---|
| Mapping A | 1 | 1 |
| Mapping B | 1 | 1 |

Table 2: Independent environment statistics

| | Goal $(0,0)$ | Goal $(0,5)$ | Goal $(5,0)$ | Goal $(5,5)$ |
|---|---|---|---|---|
| Mapping A | 2 | 0 | 0 | 0 |
| Mapping B | 0 | 2 | 0 | 0 |
| Mapping C | 0 | 0 | 2 | 0 |
| Mapping D | 0 | 0 | 0 | 2 |

Table 3: Joint environment statistics

| | Goal $(0,0)$ | Goal $(0,5)$ | Goal $(5,0)$ | Goal $(5,5)$ |
|---|---|---|---|---|
| Mapping A | 5 | 0 | 0 | 0 |
| Mapping B | 0 | 5 | 0 | 0 |
| Mapping C | 0 | 0 | 1 | 1 |
| Mapping D | 0 | 0 | 1 | 1 |
| Mapping E | 0 | 0 | 1 | 1 |
| Mapping F | 0 | 0 | 1 | 1 |
| Mapping G | 0 | 0 | 1 | 1 |

Table 4: Mixed environment statistics

| | Goal $(0,0)$ | Goal $(0,5)$ | Goal $(5,0)$ | Goal $(5,5)$ |
|---|---|---|---|---|
| Mapping A | 1 | 1 | 0 | 0 |
| Mapping B | 1 | 1 | 0 | 0 |
| Mapping C | 1 | 1 | 0 | 0 |
| Mapping D | 1 | 1 | 0 | 0 |
| Mapping E | 0 | 0 | 1 | 1 |
| Mapping F | 0 | 0 | 1 | 1 |
| Mapping G | 0 | 0 | 1 | 1 |
| Mapping H | 0 | 0 | 1 | 1 |

Table 5: Conditionally independent environment statistics

### 4 Additional mixed statistics environments

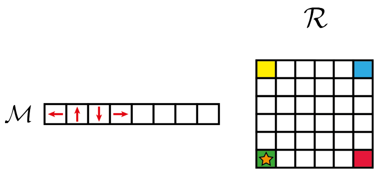

(a) Joint contexts consist of a single mapping-goal pairing, such that the mapping is perfectly predictive of the goal.

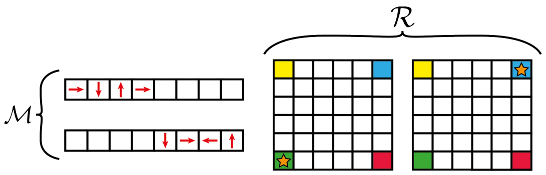

(b) Independent training contexts consist of two distinct mappings (different from that in the joint contexts), and two distinct goals (one of which is the same as in the joint context). The mappings are not always predictive of the goals.

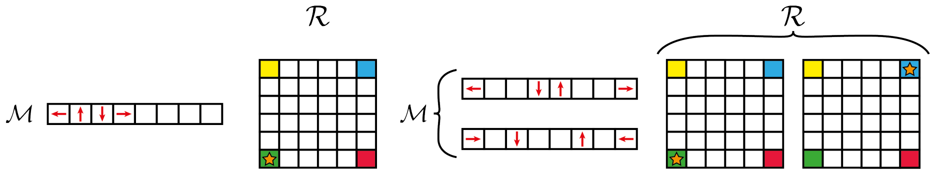

(c) Test contexts consist of two types. One context (left) is a joint context and repeats the same mapping-goal pair as in Fig. 2a. The other two contexts (right) are more independent contexts and use two novel mapping functions but the same two goals as in (Fig. 2b).

Figure 2: Additional mixed environment

While one might naturally assume that the hierarchical agent should favour the more popular goals and mappings across the entire environment, in actuality, clustering preferences depend on a complex interplay between its various CRPs. Indeed, as we shall show in two additional environments with mixed statistics, it is possible to alter the preference and have the agent favour less popular goals.

The two environments are organised somewhat differently from before (Fig. 2). The agent first visits a series of "training" contexts followed by a series of "test" contexts. The test contexts are intended to assess how the agent transfers structures and components learnt during training. Contexts within each set are visited in random order. This training-test split provides another means of assessing transfer, since we can study how the agent transfers in the test set based on what it has seen in the training set.

Training contexts consist of both *joint* contexts, where the mappings and goals are in an one-to-one relation (Fig. 2a), and *independent* contexts where mappings do not entirely predict goals (Fig. 2b). Test contexts also consist of joint and independent contexts (Fig. 2c). Joint contexts in training and test sets all use the same mapping-goal pair. Independent contexts use two goals, but importantly, one appears more frequently than the other in the training set. The goal that is more frequent is unique to the independent contexts while the other goal is shared with the joint contexts but less frequent. By manipulating the relative prevalence of these two goals in the training set, we can alter which goal the hierarchical agent will favour in independent test contexts, and this is the difference between the two versions of this mixed environment.

In one version, the hierarchical agent will favour the more popular goal, but in the other version, we can bias it to favour the less popular goal. Between the two versions, only the training set is changed; the test set remains the same. Independent training contexts use two mapping functions, but these are different from that of the joint context. Independent test contexts also introduce two new mappings, not seen in any training context. But because of this, the hierarchical agent will also need to infer which room cluster to assign these new contexts to, and that affects how it will transfer goal information. The contexts present in the two versions of the environment has been summarised in the Table 6. We shall refer to the environment where the agent favours the more popular goal at test time as "Environment 1" and the other version as "Environment 2".

##### Training contexts

Before examining agent transfer in the test contexts, we shall first summarise their behaviour in the training contexts. Again, only the hierarchical agent is optimal across all contexts here (Fig. 3a and 3d). As expected, the joint agent performs well in the joint contexts but not the independent ones, while the converse is true of the independent agent (Figs. 3c, 3b, 3f, 3e). The independent agent has a prior that favours the most popular goal, but because the most popular goal has never been associated with a joint context, the independent agent cannot recognize this and will therefore attempt the wrong goal first.

By the end of training, the hierarchical agent would have learned to assign all joint contexts to one room cluster with joint structure, and all independent contexts to another room cluster with more independent structure (Fig. 4). This clustering provides an inductive bias that determines how the agent transfers goal knowledge in the test contexts.

##### Test contexts

The agents' performance in the two environments have so far been qualitatively similar. But as we shall now see, the hierarchical agent shows different transfer behaviour in the independent test contexts.

First, we note that the joint agent is again optimal in the joint context only (Figs. 5a and 5d), showing it is successfully transferring the joint structure learnt during training. And it is again suboptimal in the independent contexts (Figs. 5b, 5c, 5e and 5f). Indeed, because the joint agent is unable to generalise compositionally, the novel mapping forces it to relearn familiar reward functions rather than transfer; thus its performance mimics that of the flat agent in these contexts.

| | Less popular goal $(0, 0)$ | More popular goal $(5, 5)$ |
|---|---|---|
| Mapping A | 4 training, 1 test | 0 |
| Mapping B | 1 training | 14 training |
| Mapping C | 1 training | 14 training |
| Mapping D | 1 test | 0 |
| Mapping E | 0 | 1 test |

(a) Environment 1. More popular goal favored in test contexts

| | Less popular goal $(0, 0)$ | More popular goal $(5, 5)$ |
|---|---|---|
| Mapping A | 4 training, 1 test | 0 |
| Mapping B | 1 training | 5 training |
| Mapping C | 1 training | 5 training |
| Mapping D | 1 test | 0 |
| Mapping E | 0 | 1 test |

(b) Environment 2. Less popular goal favoured in test contexts

Table 6: Mixed statistics environments. In both tables, the first row corresponds to the joint contexts (both training and test), the second and third rows correspond to the independent training contexts, while the last two rows correspond to the independent test contexts.

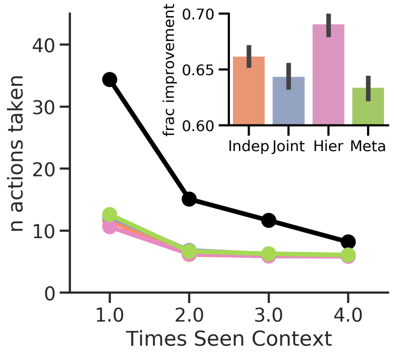


(a) Environment 1 – all training contexts

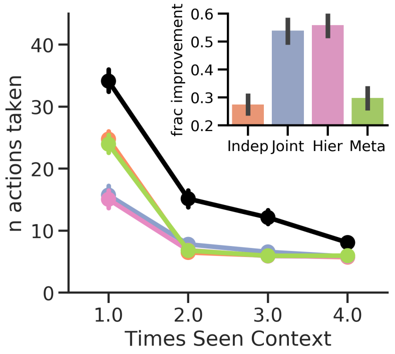


(b) Environment 1 – joint training contexts

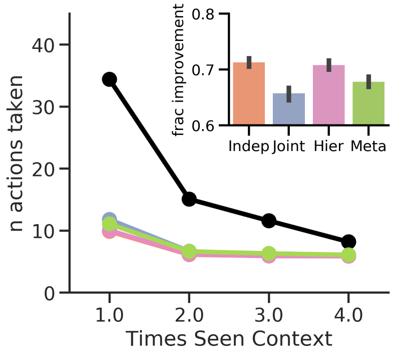


(c) Environment 1 – independent training context

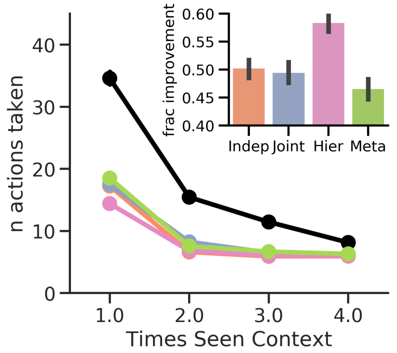


(d) Environment 2 – all training contexts

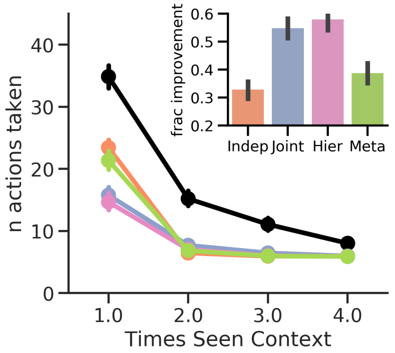


(e) Environment 2 – joint training contexts

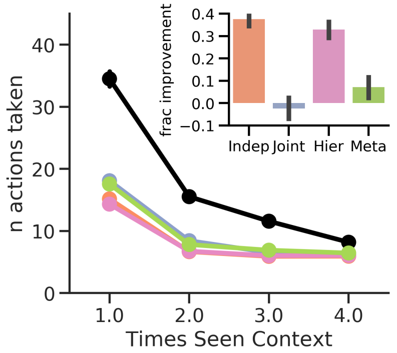


(f) Environment 2 – independent training context

Figure 3: Agent performance in the training contexts for the two environments. Errorbars show standard error of the mean.

On the other hand, the independent agent can compositionally generalise. It is suboptimal in the joint context simply because its CRP prior is biased against this goal which is less popular (Figs. 5a and 5d); it will instead attempt the incorrect (more popular) goal on its first attempt. For the same reason, it transfers poorly when independent contexts use the same goal (Figs. 5b, and 5e). But it shows positive transfer when the correct goal is the more popular one (Figs. 5c and 5f). Regardless of the context though, the agent is always optimal from the second visit onward; even when the agent attempted the incorrect goal first, it would immediately attempt the other (correct) goal next, as this is the only other goal it knows about. Notably, this recovery is something the joint agent is incapable of displaying, as it cannot transfer compositionally.

The hierarchical agent, however, shows a distinctly different pattern of behaviour in the two environments. In the joint context, it is near optimal, showing that it is successfully associating this context with the joint room cluster. But in the independent contexts, it shows a bias towards the popular goal in Environment 1, but a bias towards the less popular one in Environment 2.

As we alluded to earlier, this follows from an interplay between the various CRPs in the hierarchical agent and their different predilections for the two goals. When the agent is trying to reach a goal for the first time, it relies on its prior to decide which goal to attempt first. Specifically, the choice is determined by which room cluster the agent assigns the context to. Even though independent contexts involve novel mappings, nothing prevents the agent from assigning it to the joint cluster, thereby "breaking" the joint structure. If the context is assigned to the independent room cluster, the cluster's reward CRP will favour the more popular goal, which is what we see in Environment 1. But if the context is assigned to the joint room cluster, then the agent will favour the less popular goal, because only that goal has been associated with the joint cluster; this is what we see in Environment 2. The interplay between the CRPs determines which room cluster is ultimately chosen.

Naturally, the room-type CRP will favour the more popular room cluster, which happens to be the

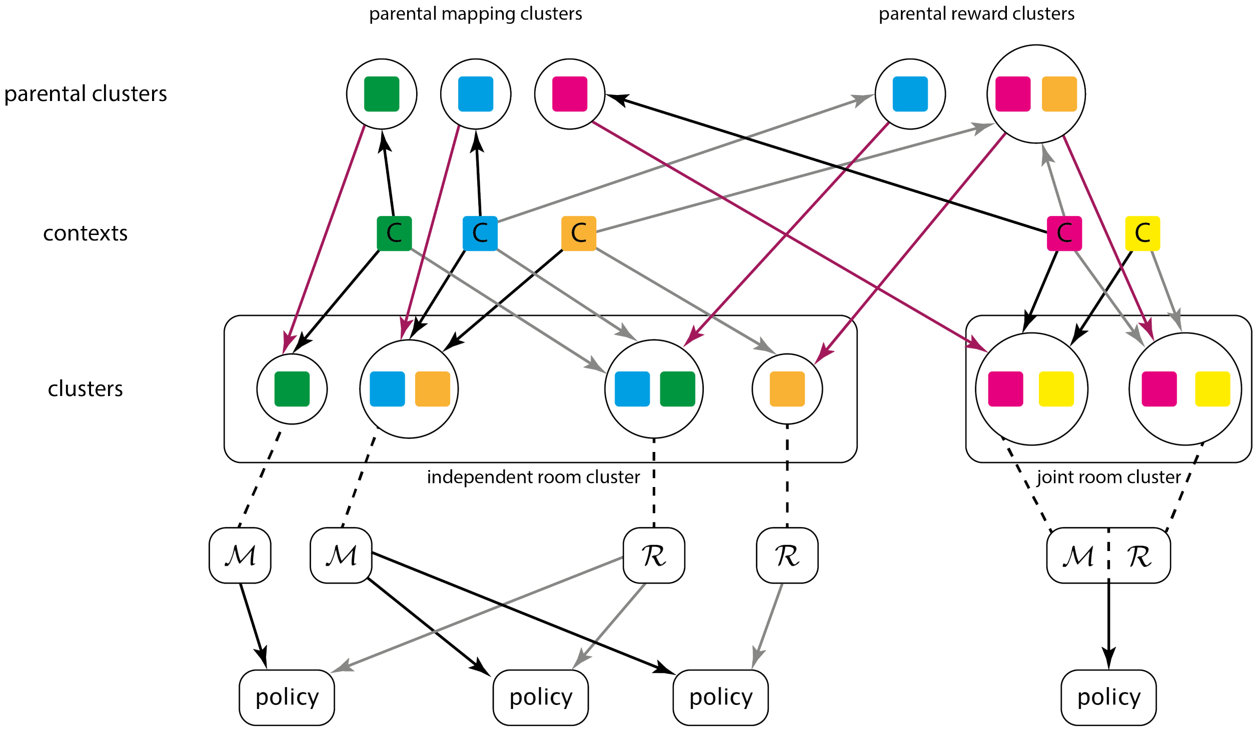


Figure 4: Hierarchical agent's clusterings after training in the mixed statistics environment. The agent discovers two room structures: an independent room cluster in which $\mathcal{M}$ and $\mathcal{R}$ are clustered independently, and a joint room cluster in which each context that forms part of the cluster is linked to a single $\mathcal{M}$ and $\mathcal{R}$ pair. Children clusters inherit from parent clusters (burgundy arrows). Contexts are assigned to a mapping cluster (black arrows) and to a reward cluster (grey arrows). Note, however, that the parental clusters at the top only receive a context assignment when a child cluster first inherits from them; all subsequent assignments to the child cluster will not involve the parent cluster. In general, the parental clusters can contain further contexts, with other children clusters inheriting from them (e.g. as in the top-right parental reward cluster), but in this example, most of the parental clusters had only one child cluster.

independent cluster in both versions of the environment. But their children CRPs will favour the joint cluster instead. This is because the decision of which room cluster to assign a context to simultaneously involves a decision of which pair of children CRPs (for the mapping and reward functions) to assign the context to. While assignment to one room cluster may be favoured by the room-type CRP, it might be disfavoured by its children CRPs. This is because the children CRPs must assign the context to one of its clusters (including possibly a new cluster), so the agent will prefer CRPs for which the probability of assignment is highest. If the context is assigned to a pre-existing cluster, then the agent will prefer the other clusters in the CRP to be far less popular or even non-existent. This is what happens with the reward CRPs. Before the agent has learnt anything about the goal in a novel context, it will prefer assigning the context to the most popular cluster in a reward CRP. From (9) in the main text, the log prior of such an assignment is

$$\log \frac{N_+}{N_+ + N_{other} + \alpha_\gamma} = \log \frac{1}{1 + (N_{other} + \alpha_{Ch})/N_+},$$

where $N_+$ is the number of contexts already assigned to the cluster, and $N_{other}$ is the number of contexts assigned to all other clusters. This is maximised not just when $N_+$ is large but specifically when the ratio $(N_{other} + \alpha_{Ch})/N_+$ is also small. If one interprets $\alpha_{Ch}$ as effectively a "prior" number of contexts assigned to a novel cluster, what this means is that when choosing between alternative children CRPs, there will be a preference for those CRPs where the contexts in the most popular cluster far outnumber the contexts in all other clusters. The reward CRP in the joint room has only one cluster, so there are no alternative reward clusters. On the other hand, there is one alternative cluster in the independent room's reward CRP. So the reward CRPs will actually favour the joint room cluster, in contrast to the room-type CRP. Indeed, this

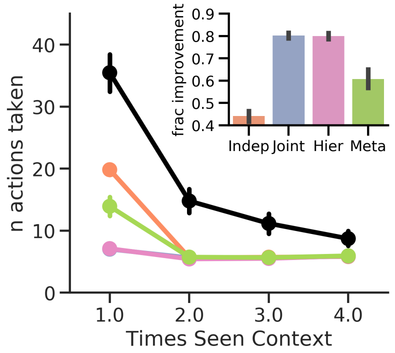


(a) Environment 1 – joint test context. The joint agent's performance is equivalent to that of the hierarchical agent and is occluded by it.

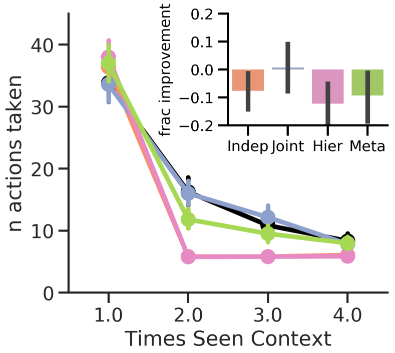


(b) Environment 1 – independent test context with less popular goal. The independent agent's performance is equivalent to that of the hierarchical agent and is occluded by it.

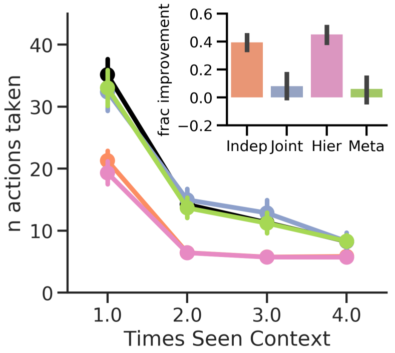


(c) Environment 1 – independent test context with more popular goal.

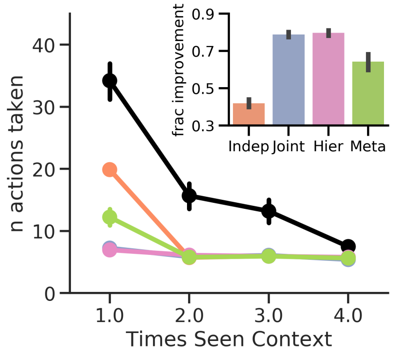


(d) Environment 2 – joint test context. The joint agent's performance is equivalent to that of the hierarchical agent and is occluded by it.

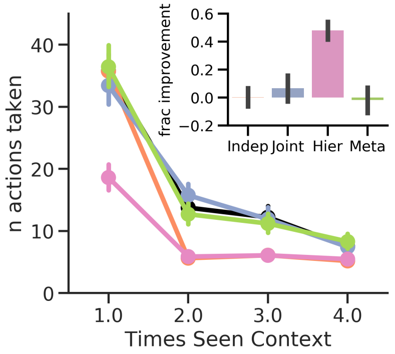


(e) Environment 2 – independent test context with less popular goal.

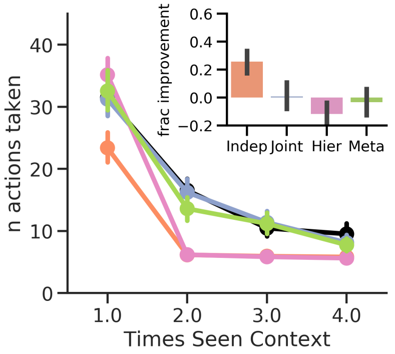


(f) Environment 2 – independent test context with more popular goal.

Figure 5: Mixed environment results. In this version, the hierarchical agent favours the more popular goal at test time, as shown by its positive transfer in (f). Graphs show average number of steps taken by each agent in the training contexts plotted against the number of visits to that context. Insets show the average fractional improvement of each agent over the flat agent baseline (black graph in main plot) during the first visit to a context. Note that only the hierarchical agent is adaptive across all contexts (g). Also note that many agents exhibit negative transfer in the independent test context that uses the less popular goal, as revealed by the inset in (e). These agents have priors that favour the more popular goal and will therefore navigate to the incorrect goal on the first attempt. However, the hierarchical and independent agents show positive transfer once they have found the correct goal, as demonstrated by their better performance from the second visit onwards in the graph. Errorbars show standard error of the mean.

is a general behaviour of the hierarchical agent whenever it assigns novel contexts to pre-existing children clusters: it will prefer CRPs for which there is only one cluster, thus encouraging jointness.

But there is also a preference for jointness when assigning contexts to novel children clusters, as is the case here with the mapping clusters. The agent quickly discovers that the mapping function is new, so it must assign the context to a new mapping cluster. From (9) in the main text, log prior of such an assignment is given by

$$\log \frac{\alpha_{Ch}}{N + \alpha_{Ch}} = \log \frac{1}{1 + N/\alpha_{Ch}},$$

where $N$ is the total number of contexts already assigned to this CRP. Like with the reward CRPs, $\alpha_{Ch}$ can be interpreted as effectively pre-assigning a "prior" number of contexts to the novel cluster. But we see that there will be a preference for CRPs where the ratio $N/\alpha_{Ch}$ is as small as possible, which translates to having $N$ be as small as possible. In other words, there will be a preference for CRPs with the fewest number of contexts already assigned to it (or even with no clusters altogether). Because the joint room cluster was far less popular overall, the mapping CRPs favour assignment to this room cluster.

Indeed, this analysis reveals several interesting effects in the interaction between the room CRP and the children CRPs within them. When a context is being assigned to a pre-existing cluster, there will be a preference for structures where the relevant cluster is more prevalent than any alternative. This has the effect of encouraging individual CRPs to represent only one type of cluster. And if both reward and mapping CRPs have only one cluster in each room, then these room clusters will have joint structures. On the other hand, whenever a new cluster is being created to represent a new reward or mapping, there will be a preference to assign this to less popular structures, thus balancing out the popularity of the different structures. In fact, there will actually be a preference for creating completely new children CRPs, which requires generating a new structure altogether. This propensity for balancing out structures will counteract the room CRP's preference to always assign a novel context to the most popular structure, while the propensity for creating new structures entirely will further encourage the generation of joint structures. In sum, there is an overall propensity for the room-type CRPs to encourage parsimony in the number of structures and hence to favour structures with more independent statistics; but there is a counteracting propensity amongst children CRPs to encourage more joint structures.

##### All contexts

Finally, we note that while some contexts favoured the joint agent and others the independent agent, only the hierarchical agent consistently showed the better performance across all contexts, demonstrating the greater flexibility of its hierarchical hypothesis space of context clusterings (Fig. 6). In Figs. 7 and 8, we have provided violin plots showing the total number of steps taken by each agent in each context type in the two environments.

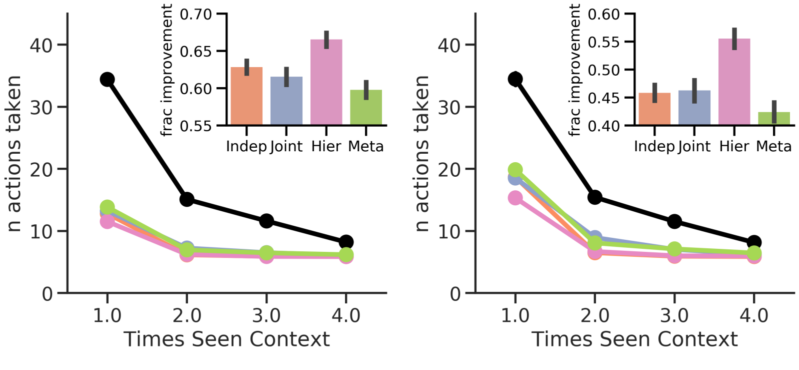


(a) Environment 1 – all contexts (b) Environment 2 – all contexts

Figure 6: Results for all contexts

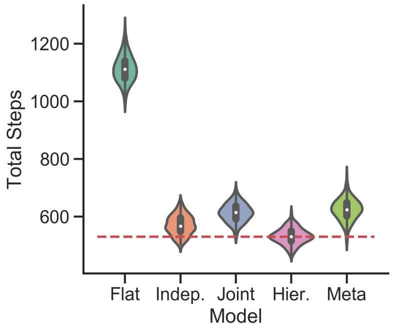


(a) All training contexts

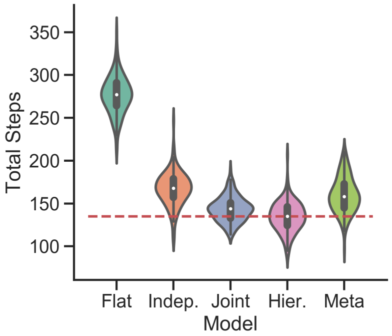


(b) Joint training contexts

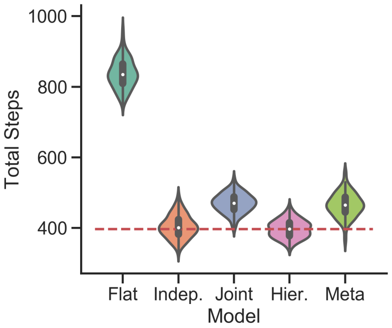


(c) Independent training context

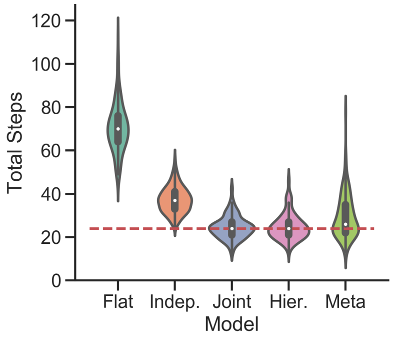


(d) Joint test context

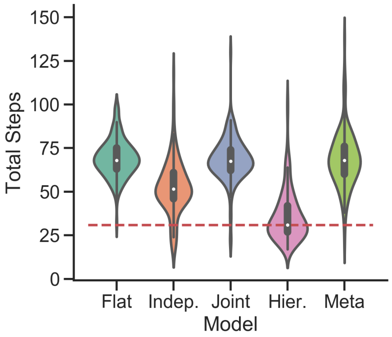


(e) Independent test context with less popular goal

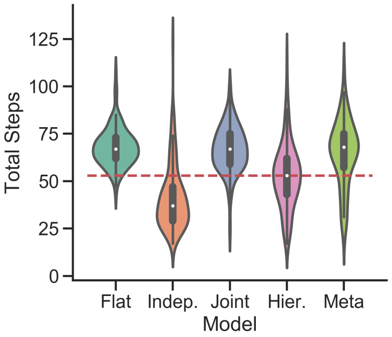


(f) Independent test context with more popular goal

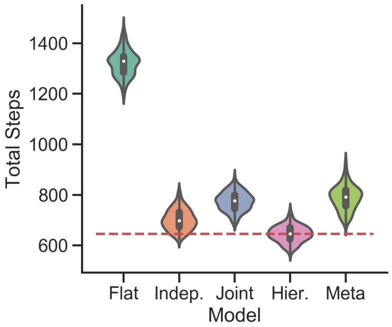


(g) All contexts

Figure 7: Non-hierarchical environment with mixed statistics (less popular goal favored by the hierarchical agent). Violin plots showing the total number of steps taken by each agent. Red dashed line shows average number of steps taken by the hierarchical agent.

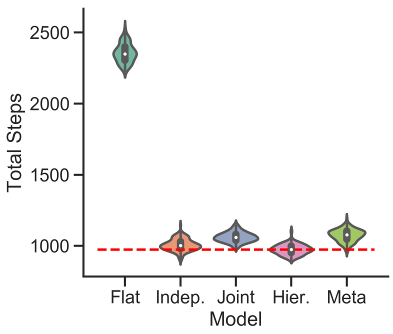


(a) All training contexts

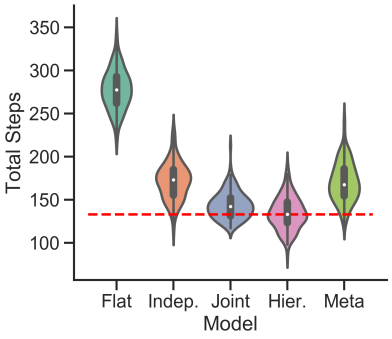


(b) Joint training contexts

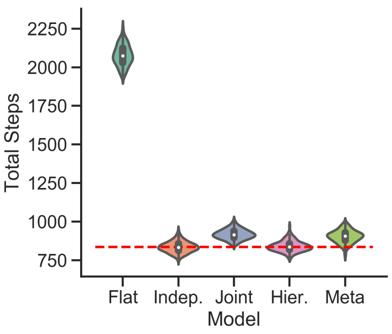


(c) Independent training context

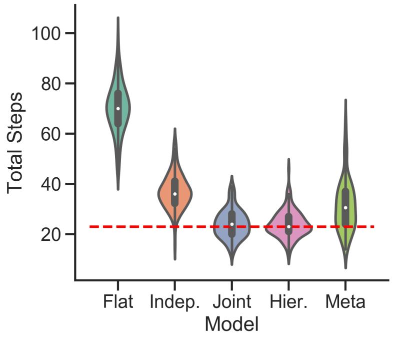


(d) Joint test context

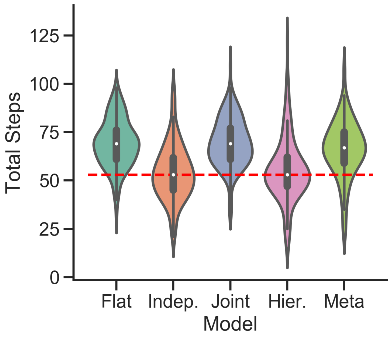


(e) Independent test context with less popular goal

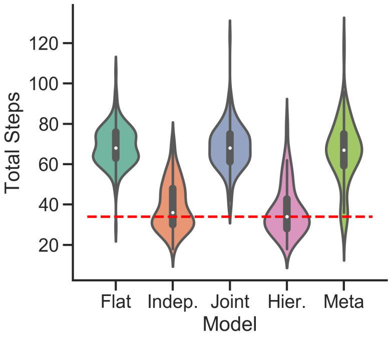


(f) Independent test context with more popular goal

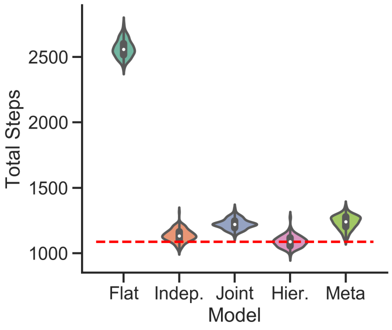


(g) All contexts

Figure 8: Non-hierarchical environment with mixed statistics (more popular goal favored by the hierarchical agent). Violin plots showing the total number of steps taken by each agent. Red dashed line shows the average number of steps taken by the hierarchical agent.

### 5 Details of the hierarchical environments

In this environment, all upper level and sublevels were $6 \times 6$ grid with doors and exits located at $\{(1,2),(2,5),(3,1),(4,3)\}$. These four possible goal locations are known to the agent, but it is of course not given information as to which location is correct. This means the reward function will now model the reward probability at the four possible goal locations only. The agent always starts in the bottom left corner, specifically $(0,0)$. Whenever the agent returns from a sublevel, it would reappear in the square that it was in immediately before moving onto the door's square. The agent receives a reward signal for reaching the correct door in a sequence or the correct sublevel exit.

A set of eight mapping functions were randomly generated for each simulation. Mappings were mutually exclusive in the sense that no two mappings shared keys that would map onto the same cardinal direction. For each simulation, a set of eight mappings would be generated. To do this, an initial random assignment of keys to actions was generated – specifically, we generated a random permutation of the keys 0 to 7 and assigned the first four keys to cardinal movements left, up, right, down, in that order. All subsequent mappings would be generated by cyclically permuting the assignment – as an example, if an initial assignment were $[stay, U, R, stay, stay, D, stay, L]$, the next two mappings would be $[L, stay, U, R, stay, stay, D, stay]$ and $[stay, L, stay, U, R, stay, stay, D]$. Finally, mappings for individual rooms and sublevels would be chosen randomly from this set according to the environment statistics. Some environments used fewer than eight mappings, so only the first few mappings in the set would be used. We note that both upper levels and sublevels used the same set of mappings. This allows agents to potentially reuse familiar mappings across all parts of the task hierarchy.

At the expense of naturalism, for the independent environments only, we allowed doors to repeat within a sequence but with different visits leading to different random sublevel environments. This was to remove as much residual joint structure in the environment as possible. (In all other environments, door sequences are random permutations of all four doors).

For each simulation, a random door sequences was chosen from among the 4! sequences possible, if doors did not repeat, or from among the $4^4$ sequences if they did.

In each room, exits for the three sublevels were generated in a similar manner. In general, exits were chosen so that the three sublevels had different exits. But to remove as much residual structure as possible in the independent environment, sublevel exits were allowed to repeat. The sequence of exits was chosen randomly from the 4! sequences possible, if exits did not repeat, or from the $4^4$ sequences possible if they did.

All environments were repeated for a total of 50 simulations.

#### 5.1 Independent statistics environment

The independent environments used four mappings, four door sequences, and four exit sequences. The prevalences of these mappings and sequences are summarised in Table 7.

| | | | | |
|---|---|---|---|---|
| Mappings | 8 × mapping A | 5 × mapping B | 2 × mapping C | 1 × mapping D |
| Door sequences | 8 × sequence A | 5 × sequence B | 2 × sequence C | 1 × sequence D |
| Exit sequences | 8 × sequence A | 5 × sequence B | 2 × sequence C | 1 × sequence D |

Table 7: Independent environment. Prevalences of mappings, door sequences, and exit sequences.

Door sequences and exit sequences were randomly assigned to each room of the 16 rooms. The 16 mappings were randomly assigned to the 16 upper levels. For each sequence position, we also have a set of 16 sublevels. For each set, we used the same 16 mappings randomly assigned them to the sublevels in the set.

#### 5.2 Joint statistics environment

This environment used eight mappings, which we shall denote mappings A through H, four door sequences, which we shall denote door sequences A through D, and four exit sequences, which we shall denote exit sequences A through D. The particular combinations of mappings and sequences is given in Table 8.

| Room type | Prevalence | Door sequence | Exit sequence | Upper mapping | Sublevel mapping 1 | Sublevel mapping 2 | Sublevel mapping 3 |
|---|---|---|---|---|---|---|---|
| Room type 1 | 4 times | Door sequence A | Exit sequence A | Mapping A | Mapping D | Mapping G | Mapping B |
| Room type 2 | 3 times | Door sequence A | Exit sequence A | Mapping B | Mapping E | Mapping H | Mapping H |
| Room type 3 | 3 times | Door sequence B | Exit sequence B | Mapping C | Mapping F | Mapping F | Mapping C |
| Room type 4 | 3 times | Door sequence C | Exit sequence C | Mapping C | Mapping F | Mapping F | Mapping C |
| Room type 5 | 3 times | Door sequence D | Exit sequence D | Mapping C | Mapping F | Mapping F | Mapping C |

Table 8: Joint environment. Combinations of mappings, door sequences, and exit sequences across different room types.

| | Room type | Prevalence | Door sequence | Exit sequence | Upper mapping | Sublevel mapping 1 | Sublevel mapping 2 | Sublevel mapping 3 |
|---|---|---|---|---|---|---|---|---|
| Joint | Room type 1 | 3 times | Door sequence A | Exit sequence A | Mapping A | Mapping A | Mapping A | Mapping A |
| Independent | Room type 2 | 1 time | Door sequence B | Exit sequence B | Mapping B | Mapping B | Mapping B | Mapping B |
| | Room type 3 | 1 time | Door sequence B | Exit sequence B | Mapping B | Mapping C | Mapping C | Mapping C |
| | Room type 4 | 1 time | Door sequence B | Exit sequence B | Mapping B | Mapping D | Mapping D | Mapping D |
| | Room type 5 | 1 time | Door sequence B | Exit sequence B | Mapping B | Mapping E | Mapping E | Mapping E |
| Independent | Room type 6 | 1 time | Door sequence C | Exit sequence C | Mapping C | Random permutation of mappings {F,F,F, G,G,G, H,H,H} | Random permutation of mappings {F,F,F, G,G,G, H,H,H} | Random permutation of mappings {F,F,F, G,G,G, H,H,H} |
| | Room type 7 | 1 time | Door sequence C | Exit sequence C | Mapping D | | | |
| | Room type 8 | 1 time | Door sequence C | Exit sequence D | Mapping E | | | |
| | Room type 9 | 1 time | Door sequence C | Exit sequence D | Mapping F | | | |
| | Room type 10 | 1 time | Door sequence C | Exit sequence D | Mapping G | | | |
| | Room type 11 | 1 time | Door sequence D | Exit sequence C | Mapping C | | | |
| | Room type 12 | 1 time | Door sequence D | Exit sequence C | Mapping D | | | |
| | Room type 13 | 1 time | Door sequence D | Exit sequence D | Mapping E | | | |
| | Room type 14 | 1 time | Door sequence D | Exit sequence D | Mapping F | | | |

Table 9: Mixed environment. Combinations of mappings, door sequences, and exit sequences across different room types.

##### 5.3 Mixed statistics environment

Like the joint environment, this environment also used eight mappings, four door sequences, and four exit sequences. The particular combinations of mappings and sequences is given in Table 9.

### 6 Further agent details

| Hyperparameter | Value |
|---|---|
| $\alpha$ | 1.0 |
| discount factor ($\gamma$) | 0.75 |
| inverse temp. ($\beta$) | 5.0 |
| $N_{min\ particles}$ | 300 |
| $N_{max\ particles}$ | 10000 |

(a) Independent clustering agent

| Hyperparameter | Value |
|---|---|
| $\alpha$ | 1.0 |
| discount factor ($\gamma$) | 0.75 |
| inverse temp. ($\beta$) | 5.0 |
| $N_{min\ particles}$ | 300 |
| $N_{max\ particles}$ | 10000 |

(b) Joint clustering agent

| Hyperparameter | Value |
|---|---|
| $\alpha_P$ | 1.0 |
| $\alpha_{room}$, $\alpha_{Ch}$ | 0.5 |
| discount factor ($\gamma$) | 0.75 |
| inverse temp. ($\beta$) | 5.0 |
| $N_{min\ particles}$ | 300 |
| $N_{max\ particles}$ | 10000 |

(c) Hierarchical clustering agent

Table 10: Agent hyperparameters for non-hierarchical environments

| Hyperparameter | Value |
|---|---|
| $\alpha$ | 1.0 |
| discount factor ($\gamma$) | 0.80 |
| inverse temp. ($\beta$) | 5.0 |
| $N_{min\ particles}$ | 100 |
| $N_{max\ particles}$ | 20000 |

(a) Independent clustering agent

| Hyperparameter | Value |
|---|---|
| $\alpha_P$, $\alpha_{Ch}$ | 1.0 |
| discount factor ($\gamma$) | 0.80 |
| inverse temp. ($\beta$) | 5.0 |
| $N_{min\ particles}$ | 100 |
| $N_{max\ particles}$ | 20000 |

(b) Hierarchical clustering agent

Table 11: Agent hyperparameters for hierarchical environments

For the hierarchical environments, we imposed a cap on the total number of agent steps; if an agent exceeded the cap, the simulation was discarded. For the diabolical environments, the agent had to find the next goal within 500 steps. For the non-diabolical environment, we instead required the agent to complete an entire room in 2000 steps.

Fig. 9 shows the full graphical model of the hierarchical agent used in the hierarchical environments.

### 7 Information content of partial door sequences

Each room contributes four partial sequences, corresponding to when 0, 1, 2, or 3 doors are revealed to the agent. Across 16 rooms, this corresponds to a total of 64 partial sequences for one simulation. Aggregated across 50 simulations, this gives a total of 3200 partial sequences.

Partial sequences were divided into two sets. Sequences where the hierarchical correctly guessed the next door but not the independent agent were grouped into one set. All other sequences were placed in the second set; these correspond to sequences where either the hierarchical agent guessed incorrectly or the independent agent guessed correctly – in any case, the hierarchical agent cannot be considered to have performed better on these sequences.

We first compared the average fractional information content in the two sets using a one-tailed t-test, assuming equal variance. The two sets were found to have significantly different information content ($t = 7.5$, $p = 1.5 \times 10^{-13}$).

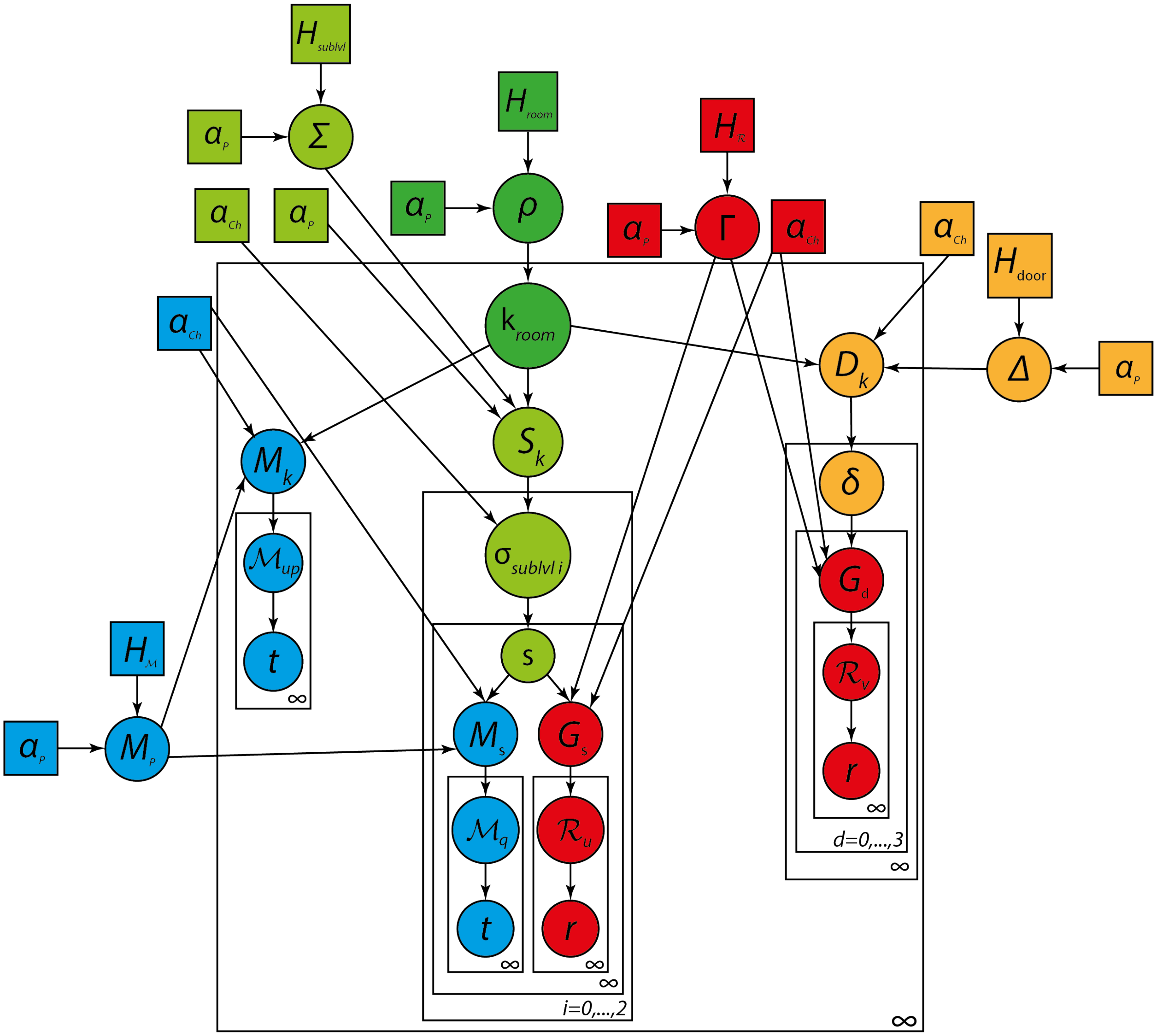


Figure 9: Hierarchical agent's generative model for the hierarchical environment. This consists of several interacting DPs and hDPs, including a DP for room clusters (dark green), an hDP for sublevel clusters (light green), an hDP for door sequence clusters (orange), an hDP for mappings (blue), and an hDP for goals (red). Base distributions $H_{room}$, $H_{sublvl}$, $H_{door}$, $H_{\mathcal{R}}$, and $H_{\mathcal{M}}$ generate environment-wide clusterings, respectively, of rooms $\rho$, sublevels $\Sigma$, door sequences $\Delta$, goals $\Gamma$, and mappings $M_P$. A room cluster $k_{room}$ is drawn from $\rho$ and contains room-specific clusterings of sublevels $S_k$, mappings for the upper level $M_k$, and door sequences $D_k$. Distribution $D_k$ generates door sequence clusters $\delta$, which in turn consists of four further goal clusterings $\{G_d\}$, one for each door position, with each cluster in $G_d$ representing the reward function $R_v$ for a door goal. Distribution $S_k$ generates a further three clusterings $\{\sigma_{sublvl\,i}\}$ of sublevel structures $s$ specific to each position in the door sequence. Each sublevel structure $s$ has further clusterings of mappings $M_s$ and goals $G_s$ specific to the sublevel. Finally, mapping clusters $\mathcal{M}$ and goal clusters $\mathcal{R}$, wherever they appear in the hierarchy, generate observed transitions $t$ and rewards $r$. All parental DPs use concentration hyperparameter $\alpha_P$, and all children DPs use concentration hyperparameter $\alpha_{Ch}$.

We then fitted a logistic model to predict the probability of the hierarchical agent performing better given a z-scored level of fractional information $\bar{i}(s)$ for a partial sequence $s$,

$$P[i(s)] = \frac{1}{1 + \exp[-(a + b \cdot \bar{i}(s))]},$$

where $a$ and $b$ are model parameters to be fitted and the z-scoring on $i(s)$ is across all simulations and all upper levels. In other words, this is the probability that $s$ belongs to the first set of partial sequences given its z-scored fractional information is $\bar{i}(s)$. The best fitting parameters were $a = -1.57$ and $b = 0.33$. This was compared against an alternative model that did not depend on the fractional information,

$$P[i(s)] = \frac{1}{1 + \exp[-a]}.$$

The best fitting parameter was found to be $a = -1.54$. We compared the models using a likelihood ratio test and found the first model to be a significantly better fit to the data ($\Lambda = 45.7$, $p = 1.35 \times 10^{-11}$).

### 8 Additional results

In the following figures, we present additional graphs for the simulations discussed in the main paper. In particular, we present violin plots showing the distribution of the total steps needed to solve an environment for each agent; distributions are over all simulations. In Fig. 13, we provide histograms showing the distribution of the total number of steps needed by each agent to solve each of the hierarchical gridworld environments.

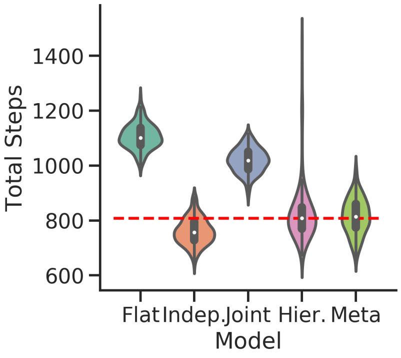


(a) Independent environment

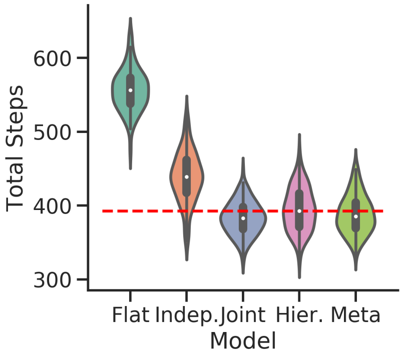


(b) Joint environment

Figure 10: Additional results for the independent and joint non-hierarchical environment. Violin plots of total steps taken across 150 simulations. Red dashed line shows average number of steps taken by the hierarchical agent.

### References

[1] Brafman RI, Tennenholtz M. R-Max – A General Polynomial Time Algorithm for Near-Optimal Reinforcement Learning. JMLR. 2002;3:213–231.

[2] Saeedi A, Kulkarni TD, Mansinghka VK, Gershman SJ. Variational Particle Approximations. JMLR. 2017;18:1–29.

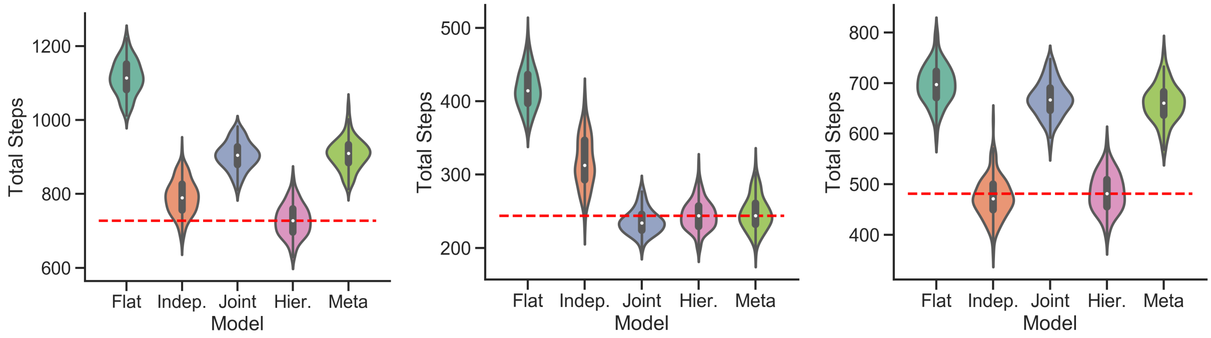


(a) All contexts (b) Joint contexts (c) Independent contexts

Figure 11: Additional results for the mixed non-hierarchical environment. Violin plots of total steps taken across 150 simulations. Red dashed line shows average number of steps taken by the hierarchical agent. Regardless of the type of context, the hierarchical agent is highly adaptive. The joint agent is only adaptive within joint contexts, while the independent agent is only adaptive in the independent contexts.

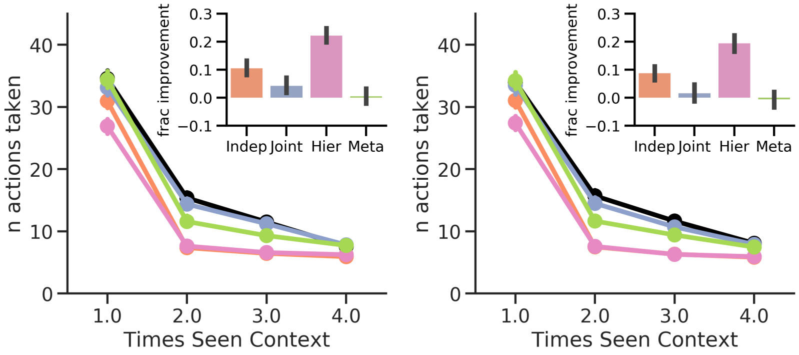


(a) Independent context set 1 (b) Independent context set 2

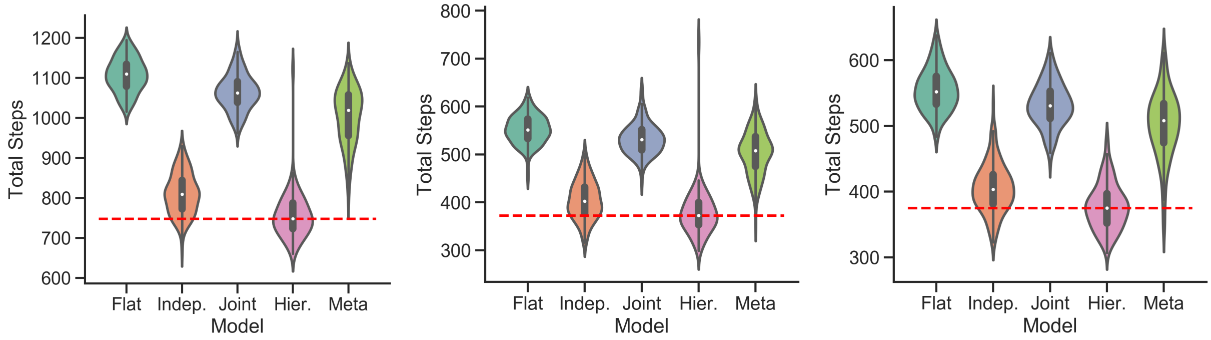


(c) All contexts (d) Independent context set 1 (e) Independent context set 2

Figure 12: Additional results for the conditionally-independent non-hierarchical environment. Red dashed lines in the violin plots show the average number of steps taken by the hierarchical agent. Conditionally independent prevents even the independent agent from being adaptive here. In this case, only the hierarchical agent is adaptive.

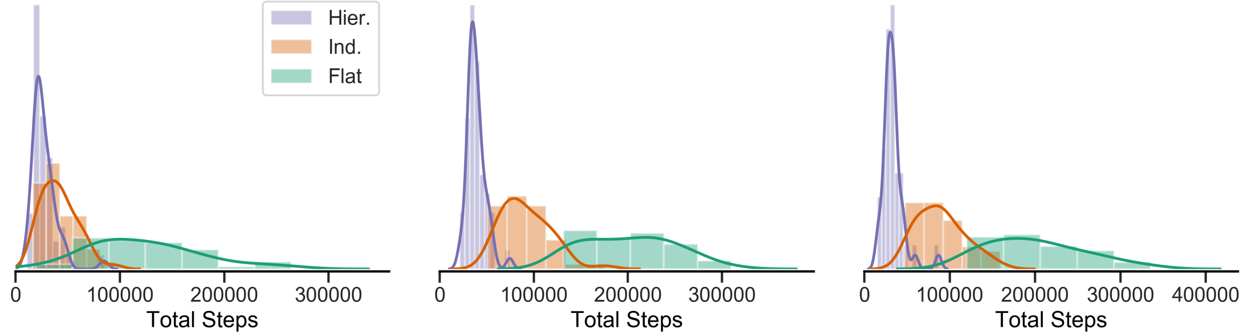


(a) Independent environment statistics (b) Joint environment statistics (c) Mixed environment statistics

Figure 13: Hierarchical diabolical rooms environments. Histograms showing the total number of steps taken by each agent in the environments with (a) independent, (b) joint, and (c) mixed statistics.

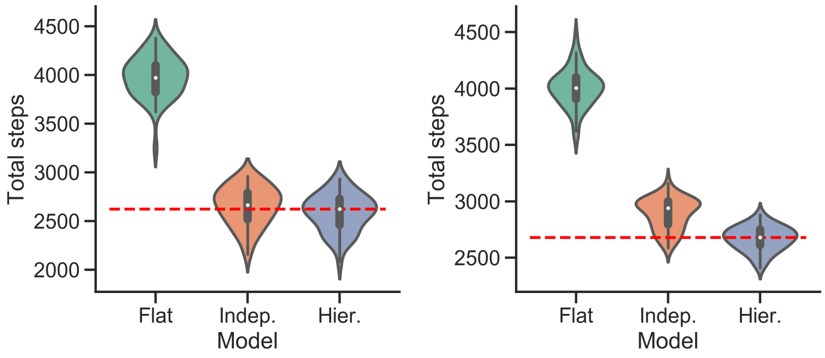


(a) Independent environment (b) Joint environment

Figure 14: Hierarchical, non-diabolical environments with (a) independent and (b) joint statistics. Violin plots of the total number of steps taken by each agent. Red dashed line shows the average number of steps taken by the hierarchical agent.

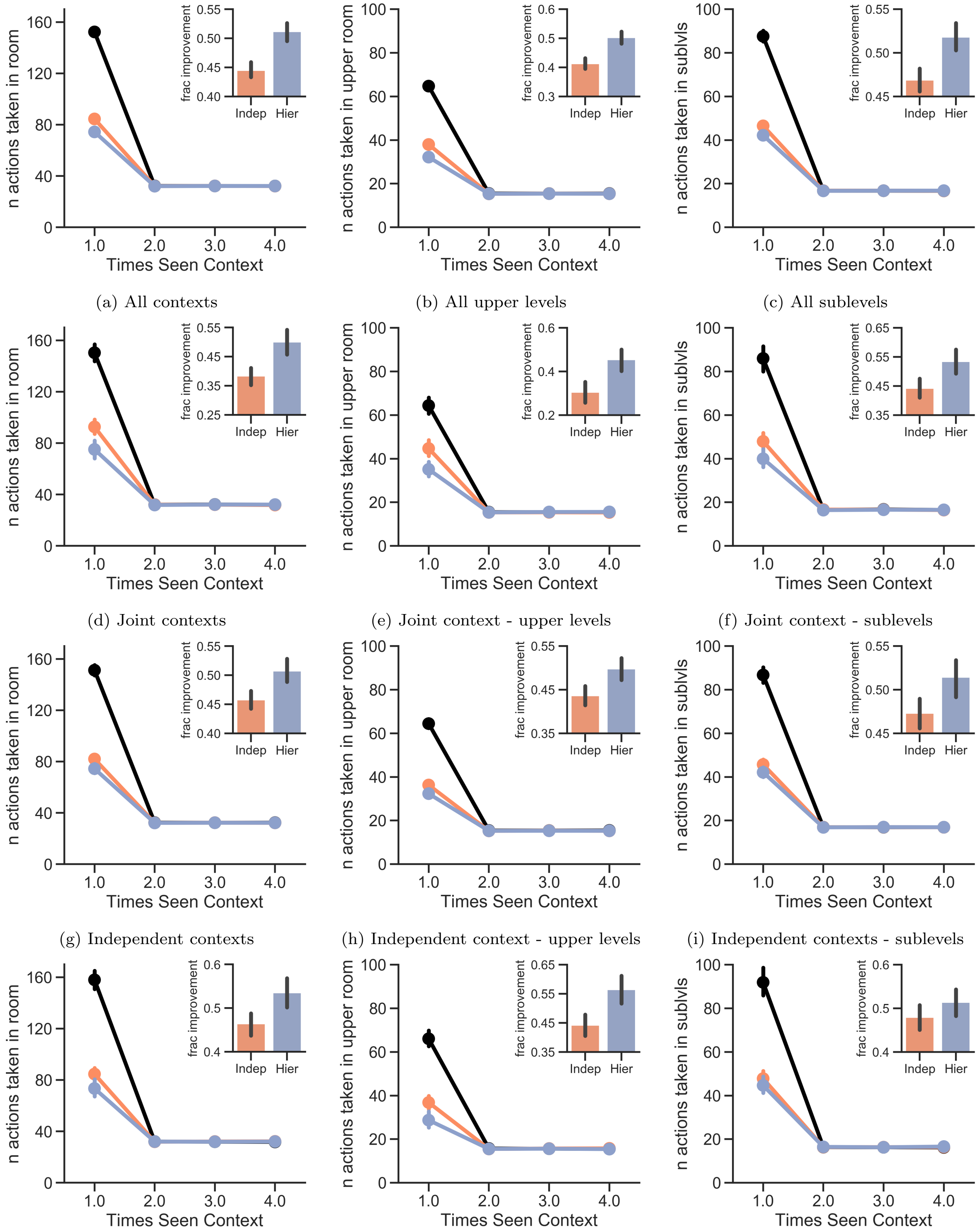


(a) All contexts (b) All upper levels (c) All sublevels

(d) Joint contexts (e) Joint context - upper levels (f) Joint context - sublevels

(g) Independent contexts (h) Independent context - upper levels (i) Independent contexts - sublevels

(j) Mixed contexts (k) Mixed context - upper levels (l) Mixed contexts - sublevels

Figure 15: Hierarchical environment with mixed statistics

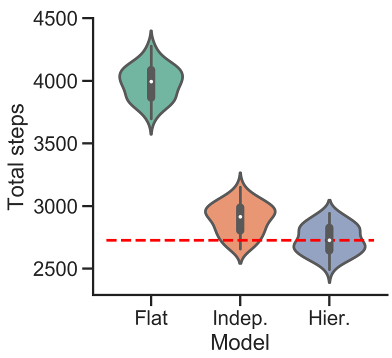


(a) All contexts

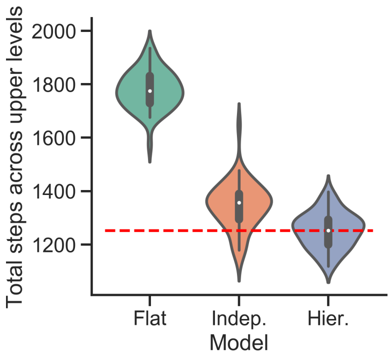


(b) All upper levels

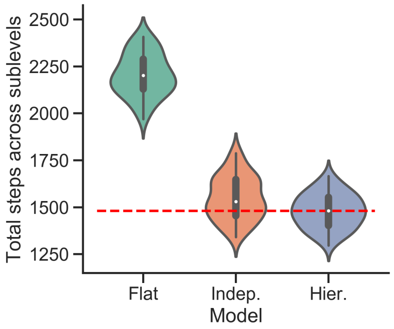


(c) All sublevels

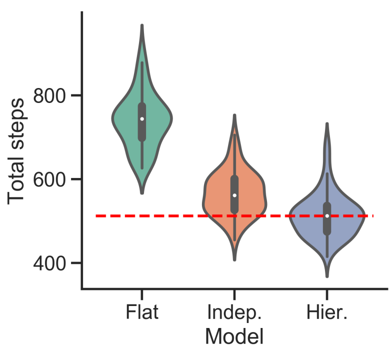


(d) Joint contexts

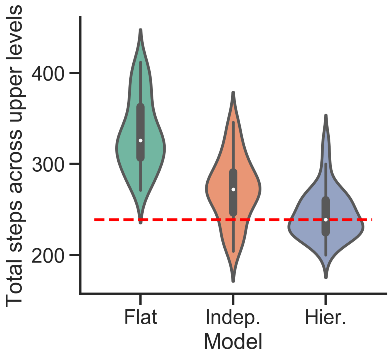


(e) Joint context - upper levels

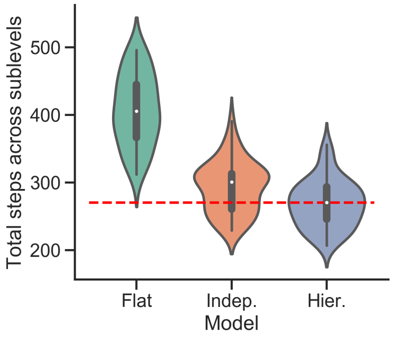


(f) Joint context sublevels

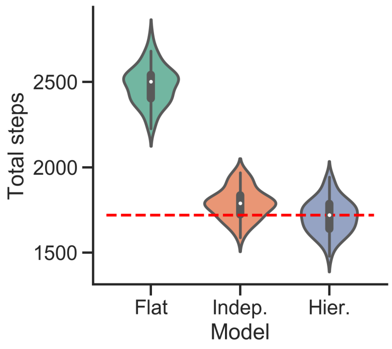


(g) Independent contexts

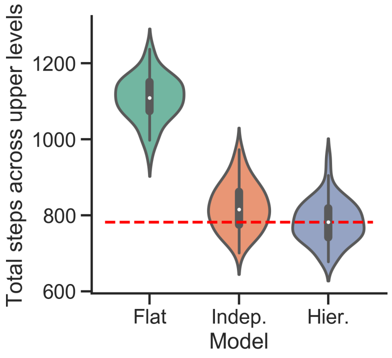


(h) Independent context - upper levels

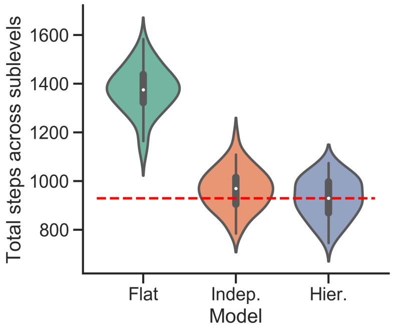


(i) Independent contexts sublevels

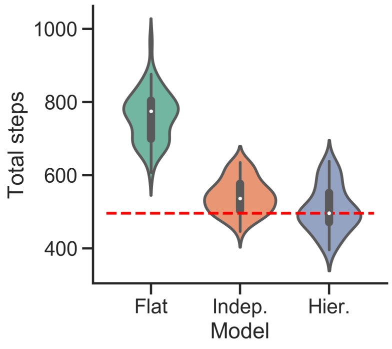


(j) Mixed contexts

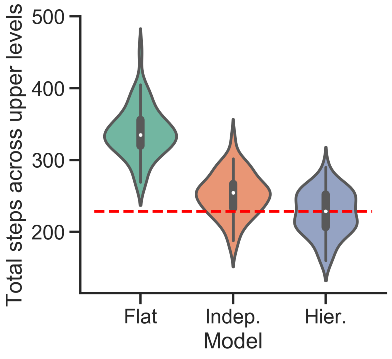


(k) Mixed context - upper levels

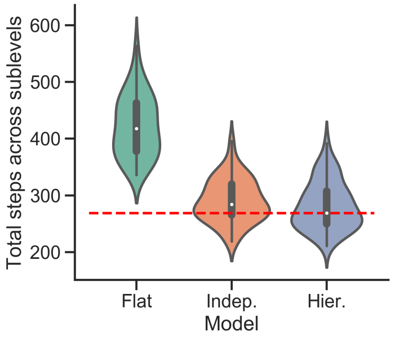


(l) Mixed contexts - sublevels

Figure 16: Hierarchical, non-diabolical environment with mixed statistics. Violin plots of the total number of steps taken by each agent. Red dashed line shows the average number of steps taken by the hierarchical agent.
